# Cross-species gene programs enable reference-free annotation of plant cells in single-cell and spatial transcriptomes

**DOI:** 10.64898/2026.11.02.763657

**Authors:** Shipeng Yang, Cheng Si, Yujiang Wu, Zhu Sun, Cai Yao, Xuemei Sun, Lihui Wang, Wei Li, Qiwen Zhong

## Abstract

Single-cell atlases span many crops, but their cells are annotated with markers from a few model plants that often lose specificity across species, leaving unclear whether plant cell types retain transferable molecular features. We present scPCIP, a reference-free method that learns cell-type programs over shared ortholog groups from 112,905 root, leaf, flower and seed cells in 68 datasets and traces each call to its genes. Across 26 held-out datasets, scPCIP outperformed existing plant annotation methods and placed spatial transcriptomic spots in anatomical tissues. In 11 held-out species, components shared by monocots and dicots discriminated cell types better than the rest of each program. Root programs trained without leaf data recognized matching tissue systems in leaves of nine species, endodermis most often marking bundle sheath; across 89 species from algae to angiosperms, the corresponding marker families were largely established in seed plants. We release all 19 programs as a gene-level atlas.

## Introduction

Single-cell and single-nucleus transcriptomes now span the life cycle of *Arabidopsis*^1^ and the major organs of rice^2,3^, soybean^4,5^ and other crops^6,7^, and comparative atlases have begun to align cell types across vascular plants^8,9^. Assigning each cell an identity, however, remains a bottleneck. Clusters are still annotated mainly with marker genes, markers validated by reporter lines or in situ hybridization exist for few species, mostly *Arabidopsis*^10,11^, and even a comprehensive *Arabidopsis* life-cycle atlas confidently annotated only three-quarters of its clusters^1^. Markers transfer poorly to crops because gene duplication, polyploidy and lineage-specific gene loss blur orthology, and cell-type transcriptomes diverge extensively even among grasses^6,8^. Protoplast isolation can add stress-induced artifacts and under-represent cell types that are difficult to release^12^, which single-nucleus profiling avoids^13^, and annotation terminology and resolution differ among studies, motivating shared cell-type ontologies^14^ and automated harmonization^15^. Beneath these practical difficulties lies a biological question: do plant cell types retain transferable molecular features despite the turnover of their individual markers?

Current approaches meet this need only in part. Marker-based methods score published gene lists^16^ or orthologous marker groups^17^ without a labelled reference from the query species but inherit the species bias and variable specificity of markers. Reference-based classifiers developed for human and animal data, such as CellTypist, scANVI and SingleR, transfer labels from annotated cells^18–20^, which most plant species lack, and cross-species integration methods bridge transcriptomes through gene homology^21–23^, with performance declining as evolutionary distance increases^24^. Plant foundation models and transformer-based annotators^25–27^ do not necessarily outperform simpler approaches without task-specific fine-tuning^28^, and their predictions are difficult to attribute to specific genes. What is missing is a method that transfers annotations to species without labelled references and shows which genes support each call.

Gene-expression programs, coordinated gene sets associated with particular cell identities or activities, offer such a basis: they can be learned from single-cell data^29^, aligned across datasets and modalities^30^ and scored in individual cells^31^. In animals, conserved regulatory components have been proposed to maintain cell-type identities while downstream effector genes diverge^32^; plant comparative atlases have identified conserved cell-type groups alongside extensive transcriptional rewiring^11^, and foundational genes shared among shoot-apex cell types of six vascular plants^9^. Three questions remain open. First, can cell-type programs learned across many studies recognize cells of species excluded from training? Second, which program components carry this cross-species information: those whose discrimination is shared between monocots and dicots, or the program as a whole? Third, can programs learned in one organ recognize related cell types in another, such as the leaf bundle sheath and the root endodermis?

Here we present scPCIP, a reference-free method that learns weighted cell-type programs in a shared ortholog-group space from root, leaf, flower and seed transcriptomes of up to 11 species per organ and reports, for each prediction, a confidence margin, the program scores and the contributing ortholog groups and query genes. We address the first question by benchmarking scPCIP across species, independent studies and spatial transcriptomes, and the second and third by using its programs to identify the components that carry cell-type identity between monocots and dicots and to relate leaf to root cell types. The programs are released as a gene-level atlas for 12 plant species, providing a compact, interpretable basis for annotating plant cells and for generating testable hypotheses about cell-type conservation.

## Results

### A shared ortholog-group space for cell-identity programs in four plant organs

Comparing cell types across species requires features that mean the same thing in every genome. Cross-species plant atlases have matched cell types through orthologous genes or by transferring annotations from model species^6–8^, and gene or protein homology also underlies cross-species integration of animal atlases^21,22^. We therefore represented each cell by the summed expression of eggNOG 5.0 Viridiplantae ortholog groups and trained one dictionary of cell-identity programs per organ in this shared feature space (Fig. 1a). Building on the concept of gene-expression programs^29^, each program is a weighted set of ortholog groups selected to distinguish one cell type consistently across training datasets and species. To limit proliferation- and stress-related signals, ribosomal, translational, histone, DNA-replication, cell-cycle and heat-response ortholog groups were excluded before the root, leaf and flower programs were learned (Supplementary Table 1).

**Fig 1.**
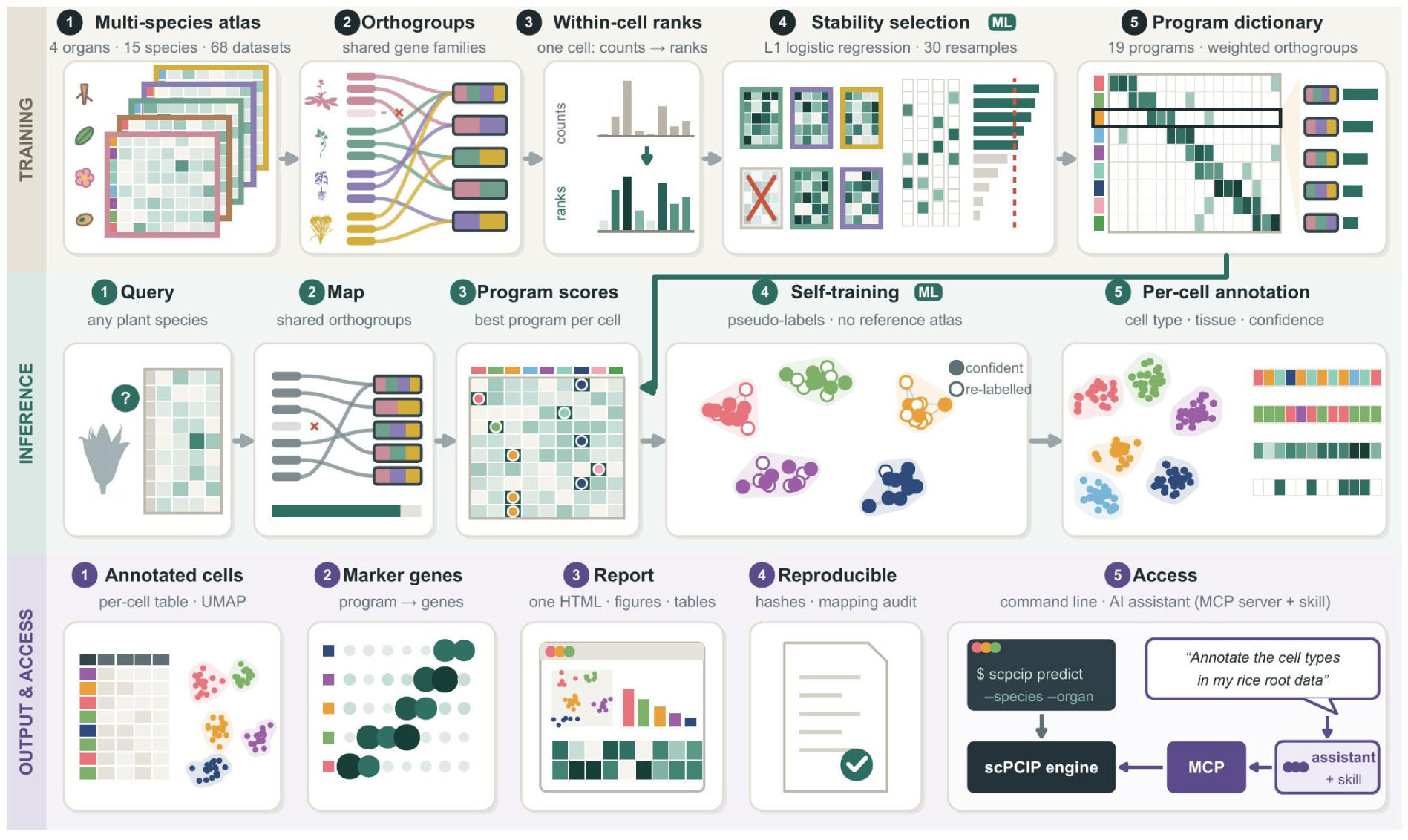
Overview of scPCIP training, inference and output. a, Training: labelled single-cell and single-nucleus datasets from four organs are represented by shared ortholog groups, from which one weighted program is learned for each cell type (Supplementary Table 3). b, Inference: cells of a query dataset are scored against the programs, and confident assignments provide pseudo-labels for one round of self-training. c, Outputs and access. ML, machine learning; MCP, Model Context Protocol. Plant silhouettes from PhyloPic.

The dictionaries were trained on 112,905 published single-cell and single-nucleus transcriptomes with raw counts and per-cell labels (Extended Data Fig. 1a, Supplementary Table 2a,b and Methods): 65,809 root cells from 31 datasets of nine species, covering eight cell types from epidermis to root cap; 30,745 leaf cells from 26 datasets of 11 species, covering epidermal, mesophyll and vascular types; 6,926 flower cells from four datasets of four Brassicaceae species in one study, covering four floral classes and a non-floral class; and 9,425 seed cells from seven datasets of three species, covering embryo, endosperm and seed coat. Labels came from the original studies or from the iPscDB re-annotation^33^, which supplied those of 44 of 68 training datasets (Extended Data Fig. 1b and Supplementary Table 2b,c). The 19 programs comprise 835 weighted program–ortholog-group entries representing 727 distinct ortholog groups (Extended Data Fig. 1c and Supplementary Table 3); with their member genes in 12 plant species, they form the scPCIP program atlas, which can be used independently of the software (Supplementary Table 3c). Stem was not included because published stem datasets sample different tissues, from internodes and culms to hypocotyls, at different developmental stages, and a stem dictionary did not exceed random gene sets in leave-one-species-out tests across three author-labelled stem datasets (Supplementary Note 3). For a new dataset, scPCIP maps genes to ortholog groups, scores programs from within-cell expression ranks without fixed absolute thresholds and returns for each cell a predicted label, a confidence margin, the program scores and ranked candidate marker genes (Fig. 1b,c and Extended Data Fig. 2a–f). Because the programs are defined on ortholog groups rather than species-specific genes, the same dictionary can be applied to species absent from training, which we tested next.

### scPCIP transfers cell-type annotations across plant species and independent datasets

We annotated 18 leaf, flower and seed datasets with dictionaries trained without the corresponding species and eight root datasets excluded from training with the released dictionary, supplemented by root leave-one-species-out tests because most of these root species were represented in training (Supplementary Table 4a and Methods). The primary metric was macro-F1, the mean F1 across cell types against published labels, compared across 18 study–organ sources (8 root and 10 leave-one-species-out). Against four plant annotation methods that require no species-specific labelled reference, OMG^17^, SPmarker^16^, scPlantLLM^26^ and scPlantAnnotate^27^, scPCIP achieved a mean macro-F1 of 0.66 across 26 held-out datasets, from 0.57 in seed to 0.71 in flower, and the same mean in the root leave-one-species-out evaluation across seven species (Fig. 2a,b and Supplementary Table 4b). It exceeded OMG by a mean of 0.17 (95% confidence interval (CI) 0.08–0.27; Holm-adjusted *P* = 0.005), in 14 of 18 sources and in all four organs (Extended Data Fig. 3a and Supplementary Table 4c,d), although not significantly in the 10 leave-one-species-out sources alone. The gain was largest in flower, where OMG assigned most annotated clusters to labels outside the evaluated floral types; OMG performed better in 5 of 26 datasets, most markedly in two soybean seed datasets with only 2–3% embryo cells (Supplementary Note 3). scPCIP also exceeded scPlantLLM and scPlantAnnotate by mean differences of 0.27 and 0.46 and, in seven root sources, the SPmarker baseline by 0.44, all significant after Holm correction (Fig. 2a and Extended Data Fig. 3a); with the released settings of both deep-learning methods, scPlantAnnotate assigned at least 99% of cells to a single type in 17 of 26 datasets (Extended Data Fig. 4d).

**Fig 2.**
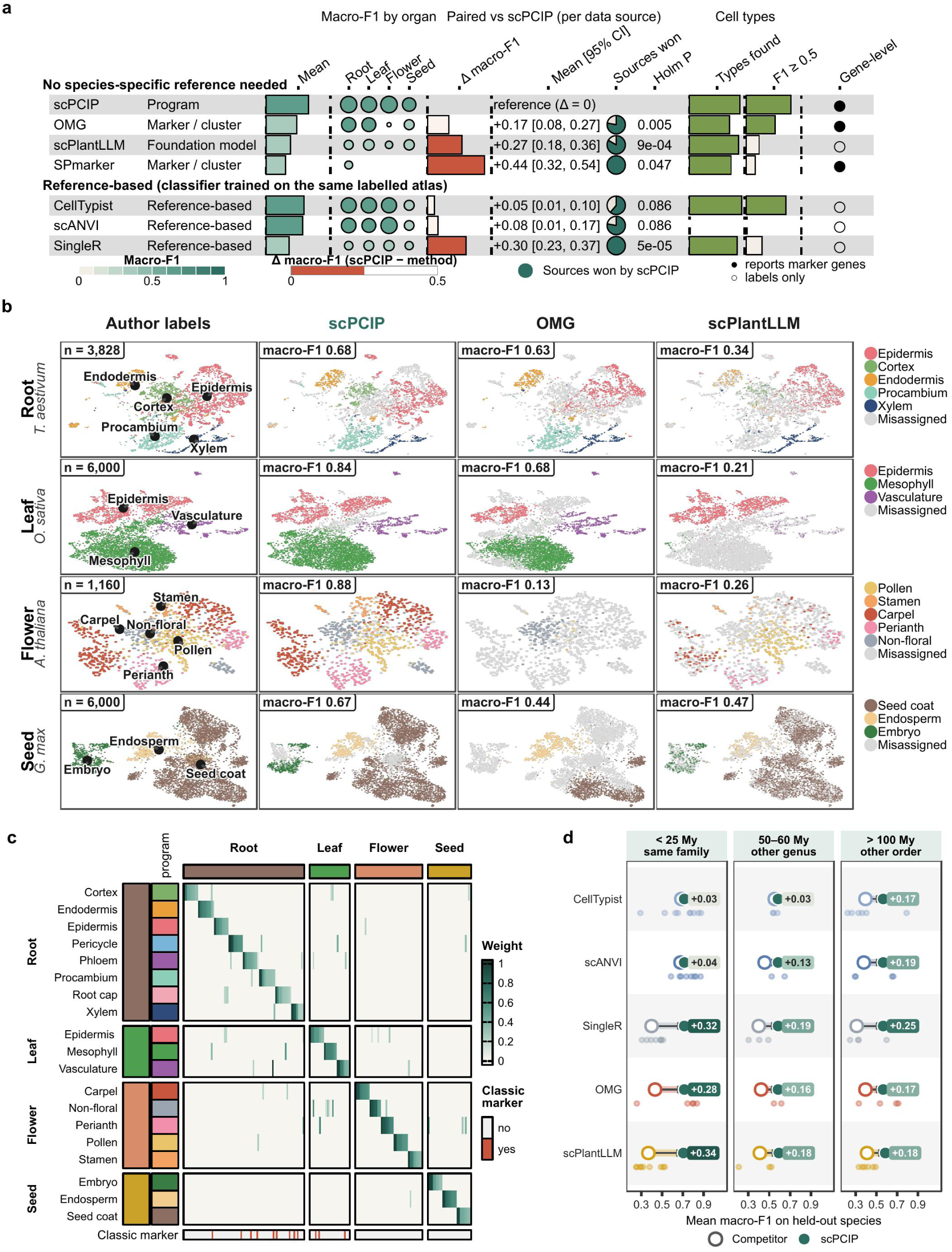
scPCIP transfers cell-type annotations across plant species and independent datasets. a, Benchmark performance across 26 held-out datasets. Paired macro-F1 differences are means across 18 data sources, or 7 for SPmarker, with 95% percentile bootstrap confidence intervals and Holm-adjusted two-sided Wilcoxon signed-rank *P* values; “Types found” and F1 ≥ 0.5 are proportions of 99 dataset–cell-type pairs, or 37 for SPmarker. Per-type results were not calculated for scANVI. b, UMAPs of representative datasets: wheat root, rice leaf, *Arabidopsis thaliana* flower and soybean seed at the cotyledon stage. The wheat dataset was scored with a dictionary that included wheat from another study, whereas the other three datasets were evaluated with dictionaries trained without the corresponding species. c, Ortholog-group weights of the 19 coarse cell-identity programs, showing 169 ortholog groups: the nine highest-weighted members of each program and additional members containing established marker genes. d, Mean macro-F1 across 18 leave-one-species-out datasets, grouped by divergence time from the closest training species. Large symbols, means; small points, individual datasets; labels, macro-F1 of scPCIP minus that of each method.

As a reference, we trained three supervised classifiers developed for human and animal data, CellTypist^18^, scANVI^19^ and SingleR^20^, on the assembled cross-species atlas in the same ortholog-group space; they used several thousand features, compared with 79–339 per organ in the coarse programs plus up to 2,000 query ortholog groups for refinement (Supplementary Note 3). scPCIP exceeded CellTypist and scANVI by mean macro-F1 differences of 0.05 and 0.08, not significant after Holm correction, and SingleR in all 18 sources, by a mean of 0.30 (Extended Data Fig. 4a,b and Supplementary Table 4c,d). The direction of these differences was unchanged across 16 independent studies, where the difference from scANVI became significant, with classifiers retrained on the root training cells of scPCIP, without scPlantDB-curated labels and with all author-provided root labels retained (Supplementary Note 3 and Supplementary Table 5c–g,j). In eight further sources from six studies outside the training data, scPCIP exceeded SingleR in all eight and OMG in 6 of 7 evaluable sources and performed similarly to CellTypist and scANVI (Supplementary Table 5a,b). Compact programs (Fig. 2c) thus matched supervised classifiers trained on the same atlas, although CellTypist had fewer dataset–cell-type pairs with very low F1 and scANVI a higher area under the receiver operating characteristic curve (AUROC) in root (Extended Data Figs. 3b and 4c).

Among the 18 leave-one-species-out datasets, scPCIP achieved a mean macro-F1 of 0.71 when the query species diverged from its closest training species within 25 million years (My), compared with 0.68 for CellTypist and scANVI, and 0.57 beyond 100 My, compared with 0.38– 0.40 for CellTypist, scANVI and OMG (Fig. 2d); the distant group comprised only seed datasets, so organ and divergence time cannot be separated. Of 99 dataset–cell-type pairs, 81% reached F1 ≥ 0.5 with scPCIP and 53% with OMG, and F1 was zero in two pairs for scPCIP and 22 for OMG (Extended Data Fig. 3b and Supplementary Table 4e); both scPCIP failures were maize leaf vascular classes consisting only of bundle-sheath cells, whose maize labels were excluded from leaf training (Supplementary Table 2c). Root cap and procambium were the most difficult root types (leave-one-species-out F1 of 0.28 and 0.51). Accuracy rose from 0.32 to 0.86 between the lowest and highest confidence-margin quartiles (Extended Data Fig. 3c), so the margin flags less reliable calls.

Finally, we withheld the entire pan-grass root atlas GSE225118^6^, including reprocessed copies of its cells, and retrained the root dictionary, leaving sorghum absent from training and maize represented only by another study. scPCIP reached macro-F1 values of 0.34 in sorghum and 0.46 in maize, above the best single ortholog group per cell type and the 95th percentile of 20 expression-matched random dictionaries, and exceeded OMG by 0.07 and 0.15 (cell-level bootstrap 95% CIs excluding zero; Extended Data Fig. 5a–c and Supplementary Table 4f). By accuracy, the primary measure specified for this test, scPCIP exceeded OMG and the random dictionaries in both species and every baseline in maize, but not the majority class or the best single ortholog group in sorghum, where cortex alone made up 39% of cells (Supplementary Note 3). Because all of these benchmarks rely on published labels, we next asked whether scPCIP annotations agree with tissue positions in spatial transcriptomes.

### Spatial transcriptomes support the anatomical placement of scPCIP annotations

Plant spatial transcriptomes are usually annotated through clustering and marker genes or by mapping to a same-species single-cell reference^3–5,7^. We applied scPCIP, with species-withheld dictionaries, to seven published spatial transcriptomic datasets of root, leaf and seed from soybean^4,5^, rice^3^ and *Arabidopsis*^34^ (Fig. 3 and Supplementary Table 6a). Cell-identity programs distinguished spots of their author-annotated tissue in 24 of 31 program–dataset evaluations (mean AUROC > 0.5 and Benjamini–Hochberg-adjusted q < 0.05 against 100 expression-matched random programs; Fig. 3a, Supplementary Fig. 1 and Supplementary Table 6b,c), with AUROCs of eight representative programs ranging from 0.60 for rice root cortex to 0.91 for soybean procambium (Fig. 3c). With the released annotation procedure, scPCIP exceeded both the majority-class baseline and the 95th percentile of 20 identically refined, expression-matched random dictionaries in 5 of 6 organ datasets other than soybean root, which was assessed by radial order (Fig. 3b,d and Supplementary Figs. 2–8).

**Fig 3.**
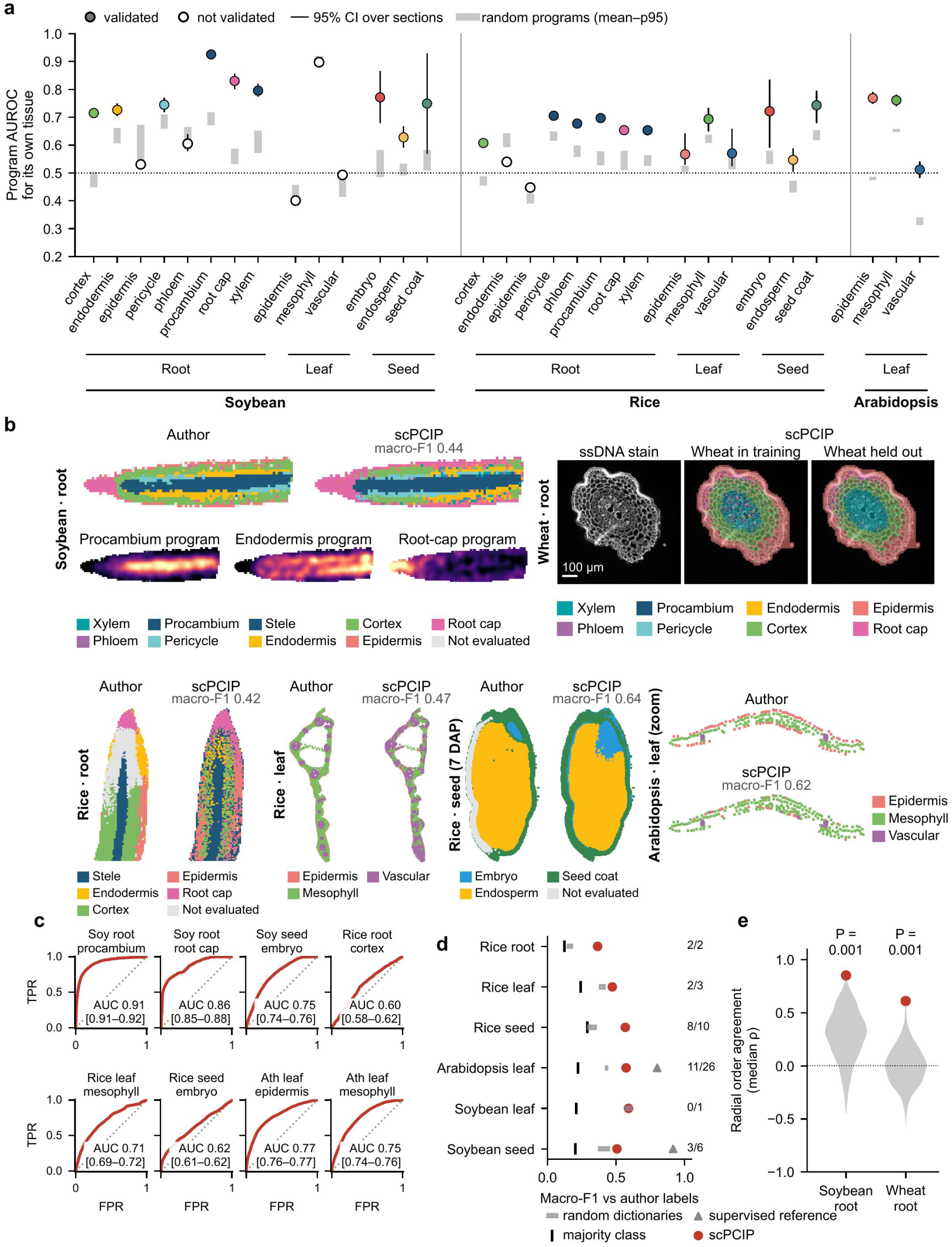
scPCIP annotates spatial transcriptomes of plant organs without a same-species reference. a, AUROC of cell-identity programs for their corresponding author-annotated classes across 31 program–dataset evaluations in seven spatial transcriptomic datasets of root, leaf and seed, scored with dictionaries trained without the corresponding species. Points, mean AUROC across sections, with bootstrap CIs for datasets with at least three sections; validated, mean AUROC > 0.5 and Benjamini–Hochberg-adjusted q < 0.05 against 100 expression-matched random programs. b, Author-provided annotations and scPCIP assignments in representative spatial sections, with macro-F1 values. The wheat root transverse section is shown with single-stranded DNA (ssDNA) staining and assignments from dictionaries trained with or without wheat. c, Spot-level receiver operating characteristic curves for eight representative programs, with AUROCs and DeLong 95% CIs from spots pooled across sections; rice seed curves pool all 16 sections, whereas panels a and d use 10. d, Macro-F1 against author-provided labels from spots pooled across sections. Random dictionaries are 20 expression-matched dictionaries with the same refinement as scPCIP; triangles, same-dataset supervised references evaluated by leave-one-section-out classification; numbers, sections in which scPCIP exceeded the section-specific random-dictionary 95th percentile. e, Median Spearman correlation ρ between inferred radial program order and anatomy across 18 soybean longitudinal and 14 wheat transverse root sections, compared with 1,000 random dictionaries, shown in grey; one-sided empirical *P* values.

Because root tissues form concentric layers^35^, spatial coordinates also provide an anatomical reference independent of author labels. In longitudinal sections of soybean root tips and transverse sections of wheat roots^7^, scored with species-withheld dictionaries (Fig. 3b, Extended Data Figs. 6a,b and 7, Supplementary Fig. 9 and Supplementary Table 6d–f), the radial order inferred from program scores correlated positively with anatomical order in all 18 soybean and 11 of 14 wheat sections (one-sided sign tests, *P* = 3.8 × 10^−6^ and *P* = 0.029; Extended Data Fig. 6c), although neither species met the pre-specified radial criteria (Extended Data Fig. 6d). Median correlations of 0.85 in soybean and 0.61 in wheat exceeded those of all 1,000 random dictionaries (Fig. 3e), and in soybean a median of 18% of high-scoring root-cap spots lay at the root tip, compared with 2.7% for the 95th percentile of random dictionaries (Extended Data Fig. 6e). Annotations transferred from single-cell data thus placed spatial spots in their expected tissues and reflected the radial organization of roots.

### Known markers, independent translatomes and phylogenetic support characterize cell-identity programs

We next asked which genes the programs contain and whether these agree with independent knowledge of plant cell types (Fig. 4a). Root program ortholog groups clustered by program when arranged by co-expression across training cell types, as expected from their selection for cell-type specificity (Fig. 4b). Of 76 literature-curated records of classic *Arabidopsis* cell-type markers, representing 70 genes and compiled independently of the training data (Fig. 4c and Supplementary Table 7), 40, including *WEREWOLF*, *GLABRA2* and *SUC2*, lay outside the shared feature space because their ortholog groups were not present in all training species. Of the remaining 36, 14 were selected by the program of their annotated cell type, two by other programs and 20 by none (Extended Data Fig. 8a), so retained classic markers were assigned to the expected cell type in 14 of 16 cases. Because such markers are commonly used to annotate plant cell populations^10^, published labels may partly depend on them; we therefore also used translating ribosome affinity purification (TRAP) profiles of rice root cell populations^36^ as an annotation-independent reference. With rice excluded from dictionary training, the expected population ranked first for 3 of 5 evaluable programs, and two of these, phloem companion cells and pericycle, also exceeded the distributions of 1,000 random programs (*P* = 0.004 and 0.034; Extended Data Fig. 8b, Supplementary Table 8 and Supplementary Note 3). Phylogenetic support varied among programs (Fig. 4d, Supplementary Fig. 10 and Supplementary Table 9): conserved cores, the ortholog groups that discriminated the corresponding cell type in at least one monocot and one dicot species, made up 69% of the endodermis program, 43% of the procambium program and 20% of the leaf vasculature program, compared with 1–10% for expression-matched random ortholog groups (Extended Data Fig. 8c).

**Fig 4.**
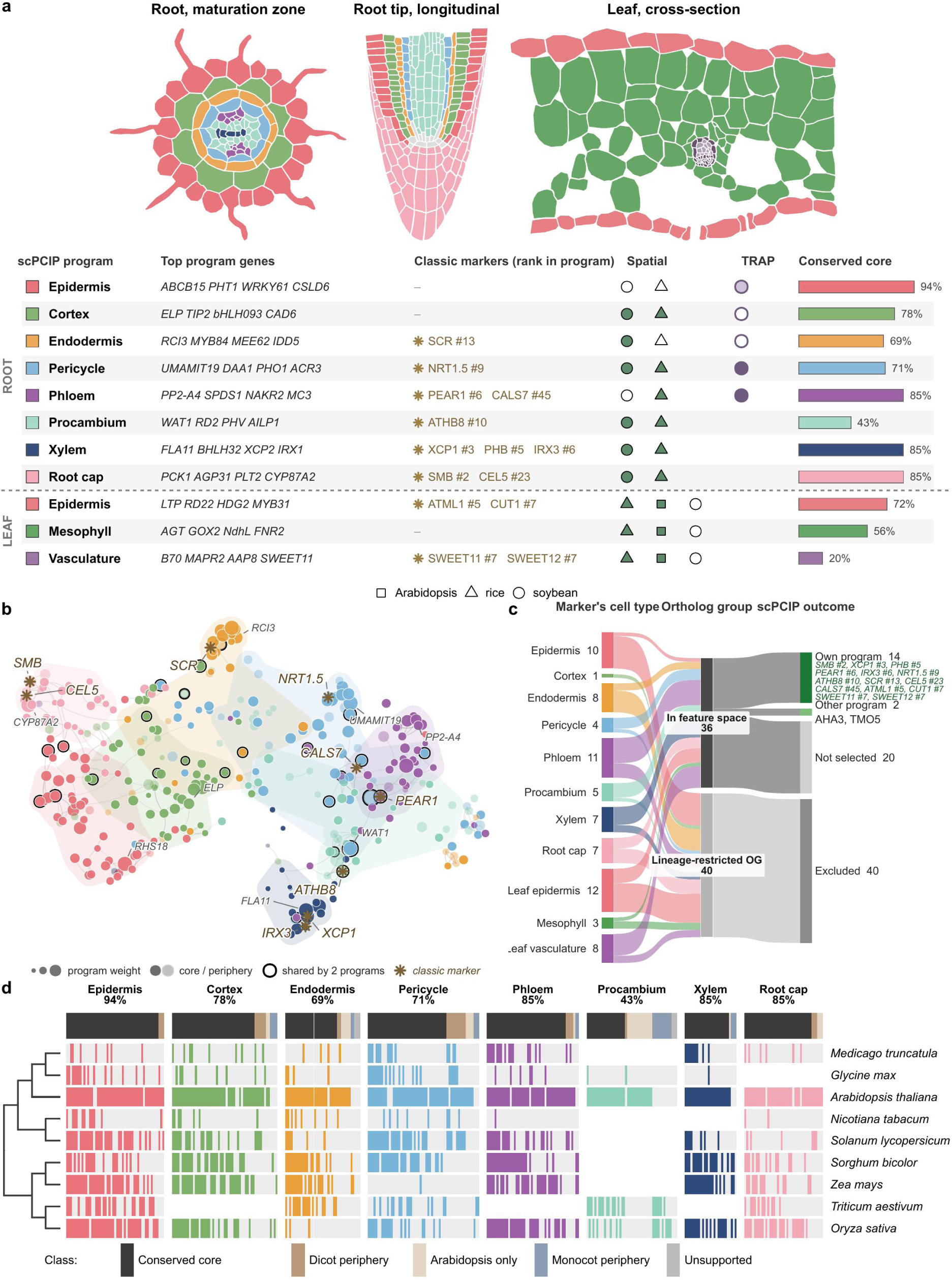
Cell-identity programs contain classic markers and differ in phylogenetic support. a, Top, schematics of a root maturation-zone cross-section, a longitudinal root tip and a leaf cross-section coloured by cell-identity program. Bottom, summary of each program: its highest-weighted genes, classic markers selected by the program with their rank in it, spatial and TRAP support, and the proportion of its ortholog groups in the conserved core. Spatial column: symbol shape, species; filled, the program distinguished its tissue with mean AUROC > 0.5 and Benjamini–Hochberg-adjusted q < 0.05 against random programs; open, these criteria were not met. TRAP column: filled, the target population ranked first with *P* < 0.05 against random programs; light, ranked first only; open, not ranked first; blank, not evaluated. b, Root program ortholog groups arranged by co-expression across training cell types. Colour indicates program assignment. c, Distribution of 76 classic *Arabidopsis* cell-type marker records across the shared feature space and cell-identity programs. d, Phylogenetic support for ortholog groups in root programs. Columns represent ortholog groups and rows represent species. Filled cells indicate AUROC ≥ 0.6 for the corresponding cell type in that species; grey, evaluated but below this threshold; white, not tested. Top bars indicate support categories. Percentages, conserved-core share of each program.

### Cross-species cell-type recognition is carried mainly by conserved program cores

Conserved regulatory cores are thought to stabilize animal cell types while effector genes turn over^32^, and root cell types of distant plant species can reuse the same genes^37^. We therefore asked whether cross-species recognition depends on program components whose discrimination is shared across lineages. Of six hypotheses pre-specified in an internal protocol (Supplementary Table 10), only the contrast between conserved cores and program peripheries remained significant after family-wise correction: in species excluded from training, cores distinguished their cell types better than peripheries of equal size by a mean AUROC of 0.054 (Holm-adjusted *P* = 0.003), in all 11 held-out species and in 33 of 47 species–program combinations (Fig. 5a and Extended Data Fig. 9a). The advantage was largest in roots (0.089, positive in all seven species), led by the endodermis program. Cores also discriminated more strongly in the training species, and in an exploratory analysis matching cores and peripheries on training AUROC removed the difference (Supplementary Note 3): discrimination already shared between monocots and dicots in training predicts recognition in unseen species.

**Fig 5.**
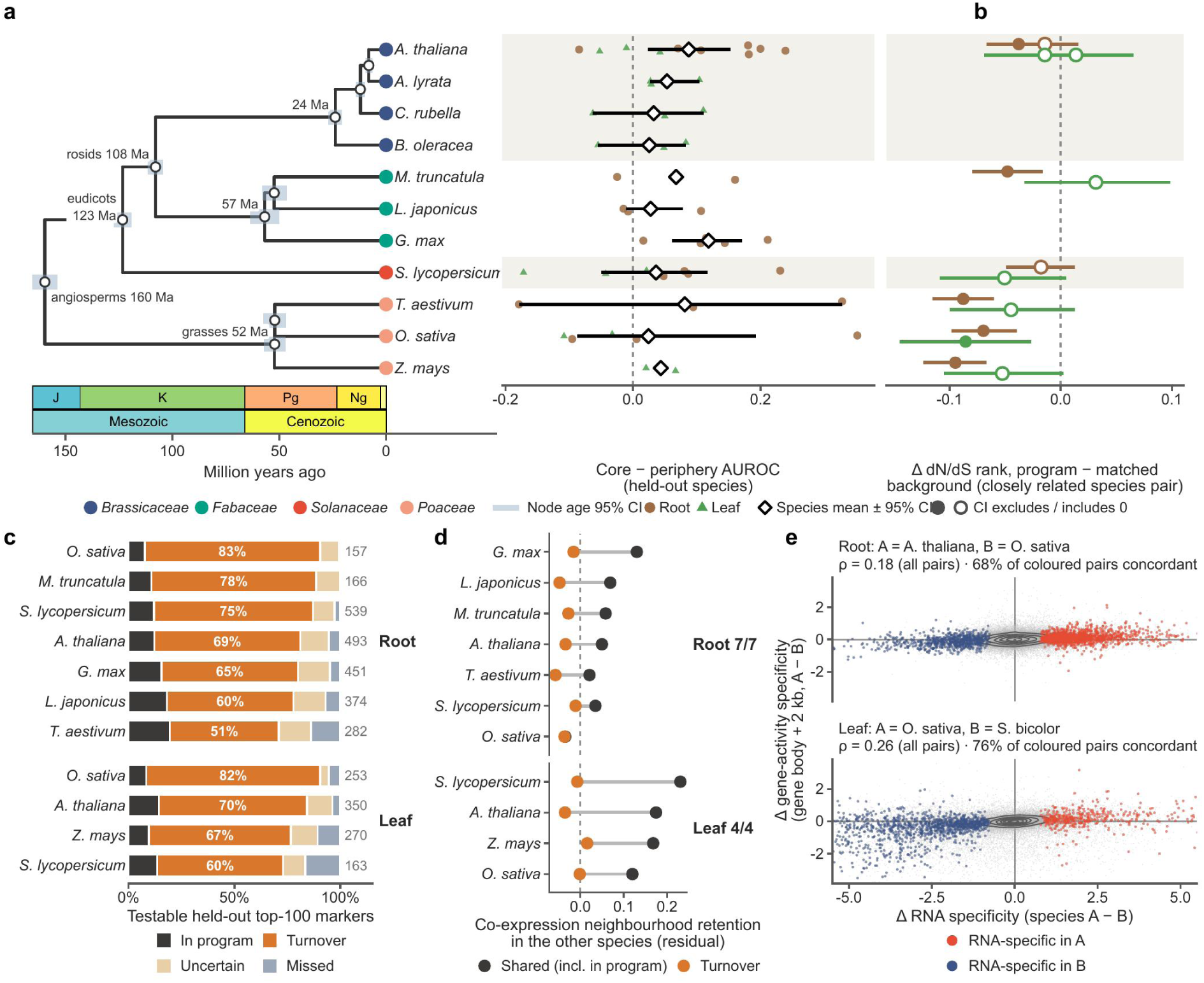
Program cores conserved between monocots and dicots carry cell-type identity across held-out species, whereas individual markers turn over. **a**, Species tree with TimeTree node ages and core-minus-periphery AUROC differences for each program at equal ortholog-group set sizes in species excluded from training. Points, the 47 species–program combinations; diamonds and bars, species means with bootstrap 95% CIs for species with at least three combinations; the formal test was a one-sided sign-flip test across the 11 species. b, Differences in dN/dS percentile rank between program ortholog groups and matched background groups for seven pairs of closely related species, plotted on the row of the species carrying the ortholog-group assignment. Points, mean rank differences; lines, 95% bootstrap CIs across program ortholog groups. c, Classification of testable marker records among the top 100 markers per cell type in each held-out dataset, pooled within species; numbers, testable marker records rather than unique genes. Markers outside the corresponding program are classified by their mean training-species AUROC: turnover, < 0.55; uncertain, ≥ 0.55 and < 0.60; missed, ≥ 0.60. d, Cross-species retention of co-expression neighbourhoods for shared markers, with training AUROC ≥ 0.60 and including program members, and for turnover markers as defined in c, expressed as residuals against matched reference ortholog groups. Points, species means; numbers, species in which turnover markers showed lower retention. e, Between-species differences, species A minus B, in RNA specificity and gene-activity specificity for ortholog-group–cell-type pairs in single-nucleus multiome data. Coloured pairs were selected on RNA specificity alone, 2,072 in roots and 1,556 in leaves, independently of the turnover definition in c; Spearman ρ is calculated across all evaluated pairs and percentages give the coloured pairs with concordant signs.

Program genes were also under stronger sequence constraint: their dN/dS percentile ranks were lower than those of background ortholog groups matched for expression, family size and protein length in all seven closely related root species pairs (pooled difference −0.053; 95% CI −0.075 to −0.031) and in 5 of 7 leaf pairs (−0.029; 95% CI −0.068 to 0.011), most strongly in grasses (Fig. 5b, Extended Data Fig. 9b and Supplementary Table 11). Individual markers behaved differently (Supplementary Fig. 11). Of 3,498 testable top-marker records from held-out datasets, ’turnover’ markers, outside the programs and with a mean training AUROC below 0.55 for their cell type, made up 51–83% in roots and 60–82% in leaves (19–29% and 25–41% below 0.50; Fig. 5c, Extended Data Fig. 9c and Supplementary Table 12a,b). In roots, this loss exceeded that between studies of the same species (73% versus 47%; species-bootstrap 95% CI of the difference excluding zero), implicating species divergence, whereas in leaves the two were similar (62% versus 49%; 95% CI of the difference −0.05 to 0.29; Supplementary Table 5h,i). Turnover markers were mostly ancient, at most 6% being younger than the angiosperm stratum (Supplementary Fig. 12), and retained their co-expression neighbourhoods in other species less well than shared markers in all seven root and four leaf species (Fig. 5d, Supplementary Fig. 13 and Supplementary Table 12c).

In an exploratory analysis of single-nucleus multiome data from *Arabidopsis*^38^ and rice^2^ roots and rice and sorghum leaves^8^, between-species differences in RNA specificity and gene-activity specificity agreed in direction for 68% of 2,072 root and 76% of 1,556 leaf ortholog-group–cell-type pairs with species-restricted RNA specificity, compared with 55% and 59% of the remaining pairs; after matching on the size of the RNA difference, the excess persisted in leaves (76% versus 65%; Fig. 5e, Extended Data Fig. 9d and Supplementary Table 12d). Cross-species cell-type information thus resides in conserved program cores, whereas many individual markers lose specificity across species and studies.

### Leaf and root cell-type programs correspond by tissue system, linking bundle sheath to root endodermis

Because all dictionaries share one feature space, cell types of different organs can be compared directly. The leaf bundle sheath provides a test case: SHORT-ROOT and SCARECROW, which act in root endodermis development, also contribute to bundle-sheath specification in *Arabidopsis*^39^, although their roles differ across lineages^40–42^. Marker sets from nine leaf and 11 root study blocks of 11 species showed that leaf and root cell types correspond by tissue system (Fig. 6a–c, Supplementary Figs. 14 and 15 and Supplementary Table 13): in held-out blocks, epidermis matched epidermis in 5 of 7, vasculature matched vasculature in 7 of 7 and bundle sheath matched the inner root layers, mostly the pericycle, in 8 of 9, all significant after Holm correction, whereas mesophyll matched cortex in only 3 of 9. These correspondences draw on gene families largely established in seed plants: across 89 species from green algae to flowering plants, families of both shared and organ-specific markers were present in 93–95% of core angiosperms and 73% of gymnosperms but in 51–61% of ferns, lycophytes and bryophytes and in fewer algae (Fig. 6a). Shared markers had modestly lower dN/dS ranks than organ-specific markers in *Arabidopsis* and maize comparisons, but not in tomato (Fig. 6d and Supplementary Fig. 16). Bundle-sheath and inner-layer markers shared eight ortholog groups, including groups associated with sulfur metabolism and potassium transport and a GRAS-family group distinct from SCARECROW (Fig. 6e and Supplementary Fig. 17).

**Fig 6.**
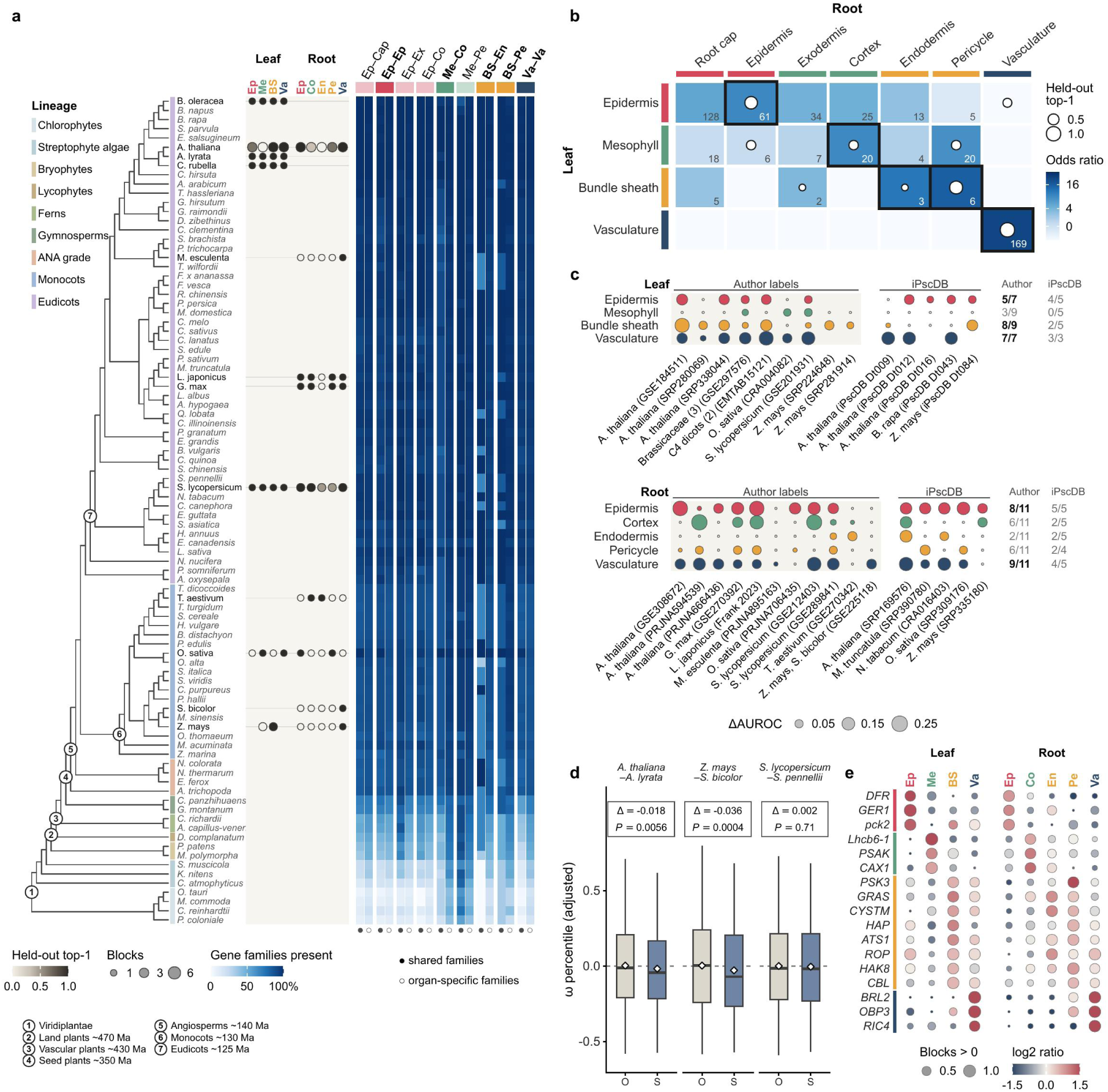
Leaf–root cell-type correspondence across tissue systems. a, Species tree from the Open Tree of Life, with branch lengths not to scale; bold species contributed single-cell data, and the two C4 dicots are omitted. Dot colour, proportion of held-out study blocks in which the pre-specified cell-type partner ranked first; dot size, number of blocks. Heat map, presence of shared and organ-specific marker families across 89 species for nine leaf–root pairs; bold columns, pre-specified pairs. Ep, epidermis; Me, mesophyll; BS, bundle sheath; Va, vasculature; Co, cortex; En, endodermis; Pe, pericycle; Ex, exodermis; Cap, root cap. b, Odds ratios of leaf–root marker overlap after removing vascular markers from non-vascular types, so that vascular–non-vascular overlaps are zero by construction. Outlines, pre-specified pairs; numbers, shared marker ortholog groups. c, Replication across nine held-out leaf study blocks and the corresponding root blocks. Coloured symbols, first-ranked pre-specified partners; size, AUROC difference from the best alternative; numbers, successful blocks; bold rows met the pre-specified criteria and remained significant after Holm correction. d, Adjusted dN/dS percentile ranks of shared and organ-specific markers in *A. thaliana*–*A. lyrata*, *Z. mays*–*S. bicolor* and *S. lycopersicum*–*S. pennellii*; ω, dN/dS; S, shared; O, organ-specific. Boxes, medians and interquartile ranges; whiskers, 1.2 × interquartile range; diamonds, means; outliers omitted. Δ, shared-minus-organ-specific coefficient adjusted for tissue system; *P*, stratified permutation test with 5,000 permutations. e, Mean log₂ expression ratios, shown by colour, and proportions of blocks with positive ratios, shown by symbol size, for eight ortholog groups shared between bundle-sheath and inner-root-layer markers and three representative groups from each other tissue system. The GRAS-family group is distinct from SCARECROW.

To ask which root program the bundle sheath most resembles, we scored leaf cells with the released root dictionary, which was trained without leaf data (Extended Data Fig. 10a and Supplementary Table 14a). Among eight root programs, endodermis best distinguished bundle-sheath cells in 8 of 12 species-by-study datasets (one-sided binomial *P* = 1.8 × 10^−5^ against a chance rate of 1 in 8) and in 7 of 9 species, spanning Brassicaceae, rice and two C4 dicots (Extended Data Fig. 10d). It also ranked first in 5 of 9 non-overlapping study blocks (*P* = 0.0025) and in all five species absent from root training (Supplementary Table 15g), and the result held after exclusion of all C4 species (endodermis first in 6 of 8 datasets; Supplementary Table 14d) and of vascular-like bundle-sheath cells (Supplementary Note 3 and Supplementary Table 15c). The endodermis program exceeded detection-matched random gene sets in 8 of 12 datasets (Extended Data Fig. 10c and Supplementary Table 14b), with vascular programs close behind (median AUROC difference of 0.05; Extended Data Fig. 10b and Supplementary Fig. 18), consistent with the position of the bundle sheath around the vein; root cortex correspondingly ranked first for leaf mesophyll in 11 of 12 datasets (Extended Data Fig. 10e and Supplementary Table 14c). Root programs therefore recognized the corresponding leaf tissue systems, supporting the hypothesis that differentiated bundle-sheath cells deploy components of an endodermis-like expression program.

## Discussion

Our first question was whether cell-type programs learned across studies recognize cells of species excluded from training. They did: compact programs over shared ortholog groups recovered published annotations across species, laboratories and organs, exceeded OMG, SPmarker, scPlantLLM and scPlantAnnotate in macro-F1 and placed spatial spots in their anatomical tissues without a same-species reference, a starting point that tissue morphology can refine. Supervised classifiers developed for human and animal data reached similar macro-F1 when trained on our labelled cross-species atlas, a resource that does not exist for most plant species. Each call is traceable to a small, weighted set of named ortholog groups and query genes, the confidence margin tracked annotation accuracy in root evaluations, and the atlas lists the member genes of every program in 12 species, for example to design small gene panels for in situ or imaging-based validation. Conformal prediction^43^ could turn the margin into calibrated uncertainty, and more resolved homology representations, potentially informed by protein language models^44^, could extend the feature space to lineage-restricted genes.

Our second question was what transfers between species. In every held-out species, components whose discrimination is shared between monocots and dicots distinguished cell types better than the rest of each program, reflecting discrimination already shared in training, whereas many individual top markers lost specificity. This extends the concept of cell-type foundational genes in vascular-plant shoot apices^9^ from gene lists to weighted programs whose cross-species recognition can be tested directly, and is consistent with the view that cell identities rest on conserved regulatory cores while effector genes turn over^32^. In roots, marker loss exceeded variation between studies of the same species, pointing to evolutionary divergence such as that documented among grasses^6,8^, whereas in leaves study-specific effects contributed comparably; program ortholog groups were also under stronger sequence constraint than matched genes, most clearly in roots. The transferable features of plant cell types are thus carried by combinations of orthologous genes rather than by single markers.

Our third question was whether programs connect cell types across organs. The relationship between leaf bundle sheath and root endodermis has long attracted attention: beyond the shared roles of SHORT-ROOT and SCARECROW^39^, endodermal tissues with Casparian bands surround leaf veins in many Pinaceae^45^, and SCARECROW is expressed in bundle-sheath cells and their progenitors in C4 *Flaveria*^46^. Our analyses add a genome-wide, cross-species view: leaf and root cell types corresponded by tissue system, and among root programs trained without leaf data, endodermis most often distinguished bundle-sheath cells, including in species absent from root training. Because marker overlap pointed more often to the pericycle and vascular programs scored closely behind, the bundle sheath is best described as sharing an inner-root-layer program with a prominent endodermal component. Bundle-sheath regulation differs among *Arabidopsis*, rice, C3–C4 intermediate Brassicaceae and maize^8,47–49^, and the root endodermis is itself heterogeneous^50^, so whether these shared components reflect common developmental origin or convergent deployment remains open. The eight ortholog groups shared by bundle-sheath and inner-layer markers, including those associated with sulfur metabolism, potassium transport and a GRAS family distinct from SCARECROW, are concrete candidates for cell-type-resolved reporter and perturbation experiments across lineages.

Several limitations define the scope of this work. Training and evaluation labels are published or database-curated and derive from clustering and marker knowledge, so agreement with them measures concordance with expert annotation, although spatial anatomy and independent translatomes provide label-independent support. Organ coverage is uneven: the flower dictionary derives from four Brassicaceae species of one study and the seed dictionary from three species, and a stem dictionary awaits more consistent datasets. Ortholog-group aggregation excludes lineage-restricted genes, including many classic *Arabidopsis* markers. Finally, scPCIP assigns every cell to a known type and does not reliably flag cell types absent from its dictionaries (Supplementary Note 2), a capability that open-set methods could add. As plant single-cell atlases expand across species, organs and developmental stages^51,52^, comparing cell types at the level of programs, between individual markers and whole transcriptomes, offers annotation that transfers without references and remains traceable to genes, and turns links between cell types into experimentally testable hypotheses about plant cell-type conservation.

## Methods

### Training data and cell-type labels

We assembled published single-cell and single-nucleus RNA-seq datasets of plant roots, leaves, flowers and seeds from public repositories including CNGB/STOmicsDB^53^ (Supplementary Table 2). Datasets were included only if they provided raw counts and per-cell labels; duplicate deposits of the same cells and cells labelled as unknown were removed, and root cell subtypes with fewer than 150 cells were not modelled. iPscDB supplied training annotations for 22 root and 22 leaf datasets, one sorghum root dataset used scPlantDB labels, one tomato leaf dataset used marker-recovered labels, and all other labels came from the original studies. With fixed label mappings (Supplementary Table 2c), root annotations were mapped through the Plant Ontology of Cell Types^14^ to eight coarse cell types, leaf annotations to epidermis, mesophyll and vasculature, and flower and seed annotations to five and three classes, respectively. Throughout, a dataset denotes one annotated expression matrix, a study one publication or data release, and a source one study–organ combination. The primary benchmark comprised 26 held-out datasets from 18 sources and 16 studies; cross-organ analyses additionally distinguish species-by-study datasets from study blocks, which pool species within multispecies studies.

### Ortholog-group representation and program learning

Proteomes were assigned to Viridiplantae ortholog groups using eggNOG-mapper v2.1.13 and eggNOG 5.0.2^54,55^. Raw counts of genes assigned to the same group were summed into a shared feature for cell c and ortholog group o:

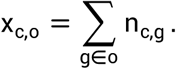

Normalized expression, where needed, was calculated as log CP10k. Before the root, leaf and flower dictionaries were trained, ortholog groups annotated with proliferation-, translation- or stress-related functions were excluded (446, 764 and 626 groups, respectively; Supplementary Table 1); the seed dictionary was trained without this filter, and query data are never filtered. Training features were restricted to ortholog groups present in every training species and converted to within-cell percentile ranks. Each program was learned for one cell type against all others in two steps. First, cells were partitioned into stability groups (species for roots, datasets for the other organs), and the ortholog groups ranking highest by median rank contrast between the target and other types across stability groups became candidates. Second, candidates were refined by stability selection^56^ over 30 rounds, each leaving out one stability group, resampling cells and fitting an L1-penalized logistic regression of the target type against all others with scikit-learn^57^; groups selected in at least 60% of rounds for root and seed, or 40% for leaf and flower, with a positive median coefficient entered the program with weights rescaled within it (Supplementary Methods and Supplementary Table 16). This yielded 19 coarse cell-type programs of 25–60 weighted ortholog groups each (Supplementary Table 3); the contribution of each learning step was assessed by ablation (Supplementary Note 1). The training cells of each released dictionary are listed in Supplementary Table 2. An exploratory stem dictionary, not part of this release, was evaluated by leave-one-species-out across three author-labelled stem datasets, each fold trained with the same procedure on the other two (Supplementary Note 3).

### Program scoring and cell-type annotation

For each query cell, scPCIP converts ortholog-group counts to within-cell percentile ranks and scores each cell-type program by the weighted mean of its available groups,

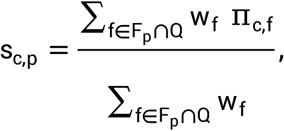

where π_c,f_ is the feature rank and Q the features present in the query. Type scores are centred on the cells that the raw argmax did not assign to that type,

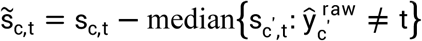

and each cell receives the call and confidence margin

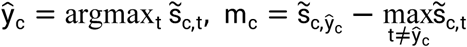

Every cell receives a call. The margin is not a calibrated probability; instead, each dictionary stores reliability estimates (per-type F1 and accuracy by margin quartile) obtained by leaving out each training species in turn. A single refinement step then trains a logistic regression on the first 30 principal components of the 2,000 most variable query ortholog groups, using cells at or above the 75th margin percentile of their called type; this model relabels all cells, and calls are harmonized within Leiden^58^ clusters computed with Scanpy^59^ when a cluster’s majority label covers at least half of its cells. Final assignments therefore depend on both the fixed programs and query-specific features. Each call is reported with its margin, program scores, ranked ortholog groups and contributing query genes with their expression summaries (Extended Data Fig. 2). Because refinement results depend on the number of linear-algebra threads, BLAS and OpenMP are fixed to one thread in the software and in all analyses.

### Annotation benchmarks and statistical analysis

Leave-one-species-out dictionaries were evaluated on 18 leaf, flower and seed test datasets, and the released root dictionary on eight root datasets excluded from its training; because all but one of these root query species were represented by other training datasets, root leave-one-species-out tests were also performed. The benchmark datasets were also used during method development. Evaluation labels came from the original studies, except for five leaf datasets and the Medicago root dataset (scPlantDB curation). Two leaf datasets were excluded, one because its labels were marker-recovered and one because marker expression did not support its labels, and root labels inconsistent with marker evidence were removed; these exclusions applied to every method, and retaining them did not change the main comparisons (Supplementary Note 3). The primary metric was macro-F1^60^, with accuracy, balanced accuracy and, where available, mean one-versus-rest AUROC as additional measures. Primary inference used mean paired macro-F1 differences across 18 sources, or seven root sources for the SPmarker-derived baseline, with percentile bootstrap CIs and two-sided Wilcoxon signed-rank tests, Holm-corrected^61^ over seven competitors; comparisons were repeated across 16 independent studies and, with Holm correction over six competitors, across the 10 leave-one-species-out sources. Divergence times were median TimeTree^62^ estimates to the closest training species. Competitors, all scored on the same test cells, were OMG, using Seurat^63^ clustering and OrthoFinder^64^ orthogroups for species absent from its bundled reference; an SPmarker-derived baseline scoring the published Arabidopsis marker list with the scPCIP rank score; scPlantLLM; scPlantAnnotate; CellTypist; SingleR; and scANVI with scArches query mapping^65,66^. CellTypist, scANVI and SingleR were trained on the assembled cross-species atlas; scPlantLLM and scPlantAnnotate were retrained with their released code on subsamples of up to 10,000 cells per training unit, scPlantLLM with the single epoch per training stage set in its tutorial; and the marker baselines used their published resources. For CellTypist majority voting and OMG clustering resolution, the setting that performed better on the test labels was used. Eight further sources, previously evaluated with earlier dictionary versions, were re-evaluated with the released dictionaries. Finally, after the pan-grass root study GSE225118 and data overlapping with it were withheld, the retrained root dictionary was compared with the best single ortholog group per type, 20 expression-matched random dictionaries, the majority class and OMG; macro-F1 is reported as the main measure for consistency with the benchmark, alongside accuracy, the primary measure specified for this test (Supplementary Table 4f).

### Spatial transcriptomic evaluation

Dictionaries trained without the corresponding query species were applied to published spatial transcriptomes of soybean, rice, Arabidopsis and wheat. In seven organ datasets, program scores were evaluated by AUROC against annotated spatial domains and compared with 100 expression-matched random programs, with Benjamini–Hochberg correction^67^ across 31 program tests. Annotation performance was assessed by pooled-spot macro-F1 against a majority-class baseline and 20 expression-matched random dictionaries processed with the same refinement, and without refinement as a sensitivity analysis. To assess anatomy without author cell-type labels, the characteristic position of each root type in soybean and wheat sections was defined from its highest-scoring spatial bins and compared with anatomical radial order by Spearman correlation, sign tests and 1,000 random dictionaries. Because the pre-specified radial criteria were not met in either species (Extended Data Fig. 6d and Supplementary Table 6d), we report the direction of correlation in each section and the median correlation against random dictionaries, with sections filtered by an author-labelled stele-bin criterion (Supplementary Note 3). Sections are not independent biological replicates.

### Program interpretation and evolutionary conservation

Program composition was examined with UMAP^68^ layouts of within-dataset-centred pseudobulk co-expression, literature-curated Arabidopsis marker records and independent rice root translatomes scored with a dictionary trained without rice. The conserved core of each program comprised ortholog groups with cell-type AUROC ≥ 0.6 in at least one monocot and one dicot training species. Six hypotheses on program conservation were pre-specified in an internally recorded protocol and tested with Holm correction. In the main core–periphery test, cores and peripheries were redefined in each leave-one-species-out fold, compared at equal feature-set sizes and summarized across 11 held-out species and 47 species–program units with a one-sided exact sign-flip test. Because covariate imbalance made the pre-specified matching control uninformative, matching on expression, detection, family size and training discrimination is reported as an exploratory analysis. Marker turnover was characterized from held-out marker AUROCs in training species, within-species between-study comparisons, cross-species preservation of co-expression neighbourhoods and exploratory RNA–chromatin associations. Gene age was assessed with hierarchical orthogroups, and sequence constraint with dN/dS of reciprocal-best-hit orthologs computed with PLAZA^69^, DIAMOND^70^, MAFFT^71^ and yn00^72,73^; for each organ, program-minus-background differences were averaged within each of seven species pairs and then across pairs, with 95% CIs from 10,000 bootstrap resamples of program ortholog groups drawn jointly across pairs.

### Cross-organ cell-type correspondence

Leaf–root correspondence was examined in the common ortholog-group space with two complementary analyses. First, marker groups reproducibly enriched in each organ were compared by Fisher’s exact test after vascular markers had been excluded from non-vascular types, and proposed cell-type correspondences were assessed by leave-one-study-block-out ranking across nine leaf and 11 root study blocks with Holm-corrected Poisson-binomial tests. Marker families were mapped from A. thaliana to 89 species with OrthoFinder and to rice through eggNOG, and shared and organ-specific marker groups were compared for family presence and sequence constraint (Fig. 6 and Supplementary Table 13). Second, the released root dictionary was applied without retraining to 12 species-by-study leaf datasets from nine species, with rice mestome sheath counted as bundle sheath. Each root program’s AUROC for annotated bundle-sheath cells against other leaf cells was computed from the exported centred type scores before refinement; because centring subtracts one constant per type within a dataset, these AUROCs and rankings equal those from uncentred weighted-rank scores. The pre-specified primary statistic was the number of datasets in which endodermis ranked first among eight root programs, tested against a chance rate of 1 in 8 with a one-sided binomial test; because datasets share species and studies, non-overlapping study blocks and lineages were summarized as secondary units (Supplementary Table 15g). Controls comprised detection-matched random gene sets computed on uncentred scores, comparison with the best vascular program and sensitivity exclusions of the two C4 dicots and of all C4 species including maize (Supplementary Table 14d). Marker–program comparisons, filtering of vascular-like bundle-sheath cells and analyses of published rice and maize chromatin accessibility were exploratory (Supplementary Table 15); accessibility data were not used to train the dictionaries.

### Pre-specification, deviations and reproducibility

‘Pre-specified’ denotes criteria recorded in internal protocols before the corresponding computation; the study was not externally registered. Confirmatory and exploratory analyses are distinguished throughout, and protocol departures are listed in Supplementary Note 3: the benchmark label exclusions, the main measure reported for the GSE225118 test, the re-evaluation of eight sources, the spatial section filter and dictionary version, the matching controls of the core–periphery and bundle-sheath tests, and the C4 classification in the cross-organ analysis.

## Use of AI tools

Claude and Codex were used to assist with writing. The authors checked all analyses, results and text and take full responsibility for the content.

## Data availability

Accession numbers, original publications and label sources are provided for every training, held-out benchmark and spatial transcriptomic dataset (Supplementary Tables 2, 4 and 6, respectively). Training data were obtained from GEO, CNGB/STOmicsDB, figshare and Zenodo^2,3,5,6,51,74–80^ and from datasets re-processed by iPscDB^33^, with the original studies given in Supplementary Table 2. The held-out benchmark used root datasets GEO GSE198394^81^, figshare 10.6084/m9.figshare.23986200^82^, GEO GSE226149^83^, GEO GSE314252^84^, GEO GSE270342^7^, NCBI BioProject PRJNA1218210 / CNCB PRJCA035692^85^, CNCB PRJCA011245^86^ and the root samples of GEO GSE232863^2^; leaf datasets GEO GSE297576^74^, CNCB CRA004082^87^ and NCBI SRA SRP224648, SRP280069, SRP281914 and SRP338044^88–91^, with labels curated by scPlantDB^92^ where the authors did not provide per-cell labels; flower datasets GEO GSE297576^74^; and seed datasets GEO GSE232863, GSE270392 and GSE295007^2,5,51^. The eight re-evaluation sources were CNGB CNP0002614^93^, GEO GSE224539 and figshare^94^, GSE240098^95^, GSE303996^96^, GSE273033^97^ and GSE313726, which had no associated publication at the time of analysis. The independent study was GEO GSE225118^6^. Spatial transcriptomes were STOmicsDB STDS0000168^4^, STDS0000421^3^, STDS0000104^34^ and STDS0000332 / GEO GSE270392^5^, and GEO GSE271725^7^. Rice root TRAP-seq data were GEO GSE180100^36^, and single-nucleus multiome data were GEO GSE235510, GSE232863 and GSE248919^2,8,38^. Chromatin accessibility atlases were GEO GSE252040^98^ and GSE155178^99^. Leaf datasets in the correspondence analysis also included GEO GSE184511^91^, GSE201931^75^ and ArrayExpress E-MTAB-15121^100^. Stem datasets collected for the unreleased stem dictionary are listed in Supplementary Table 2. The trained dictionaries and gene-to-ortholog-group maps are available from Hugging Face at https://huggingface.co/shipengyang/scPCIP. The scPCIP program atlas v1.0, listing the 19 programs with their weights and member genes in 12 species, is provided as Supplementary Table 3c and is archived with the released dictionaries and the gene-to-ortholog-group map at Zenodo (https://doi.org/10.5281/zenodo.24411388). Source data are provided with this paper.

## Code availability

scPCIP v1.0.0 is available under the MIT licence from PyPI, installed with pip install scpcip, and from GitHub at https://github.com/Shipeng-Yang/scPCIP (release tag v1.0.0) as a Python package and command-line tool, with optional R figures, a container image, an MCP server and an agent skill for use with AI assistants. Analysis scripts that generate every figure and table will be made publicly available on publication.

## Supporting information

scPCIP_Supplementary_Information

scPCIP_Supplementary_table

## Author contributions

S.Y. conceived the study, designed the methods, curated the data, developed the scPCIP software, performed the analyses, prepared the figures and wrote the manuscript. C.S. contributed to data curation and analysis and revised the manuscript. Y.W., Z.S., C.Y., X.S., L.W. and W.L. contributed to data collection and interpretation and revised the manuscript. Q.Z. supervised the project, acquired funding and revised the manuscript. All authors read and approved the final manuscript.

## Acknowledgements

We thank the authors of the public single-cell and spatial transcriptomic datasets analysed here for making their data available, and the developers of iPscDB, scPlantDB and ggPlantmap.

---

**Extended Data Fig. 1.**
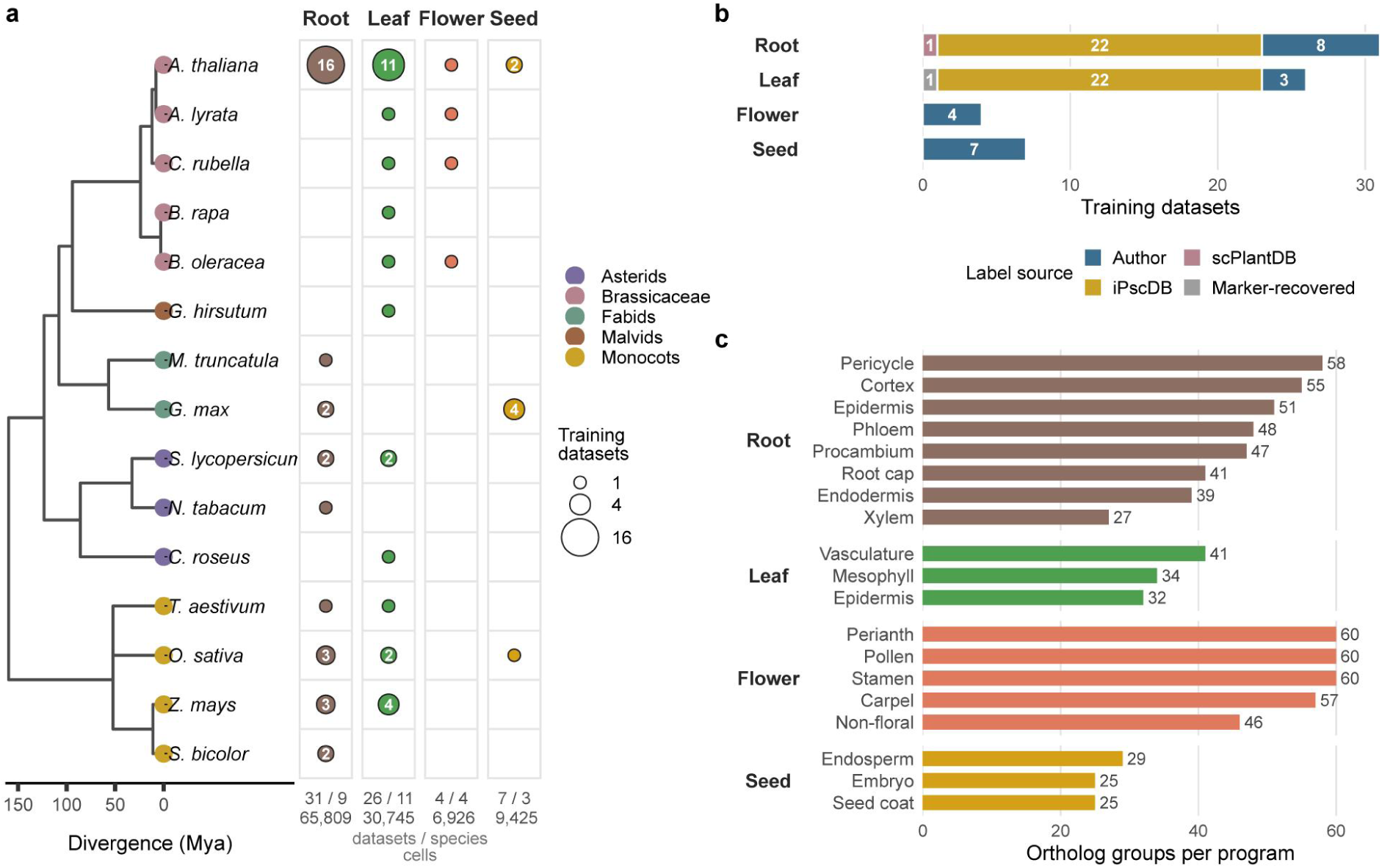
Training compendium. a, Number of training datasets per species and organ, shown by dot size, on a TimeTree species tree. b, Source of training labels. c, Ortholog groups per program.

**Extended Data Fig. 2.**
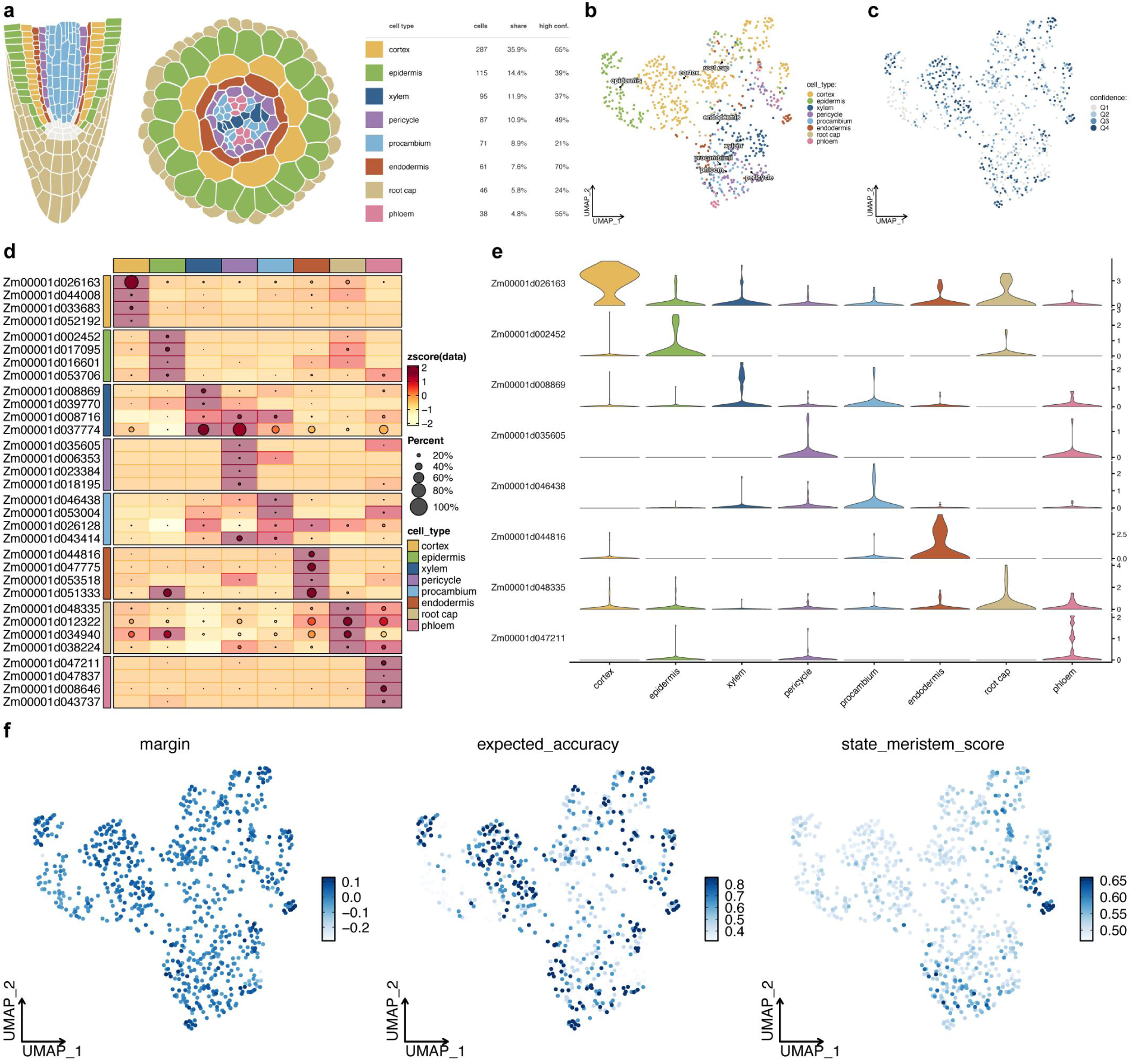
scPCIP output for an example dataset. Maize root example distributed with the software. a, Anatomy schematic coloured by call frequency. b, UMAP coloured by call. c, Confidence quartile. d, Top candidate marker genes per type. e, Expression of the top marker of each type. f, Confidence margin, expected accuracy and meristem state score.

**Extended Data Fig. 3.**
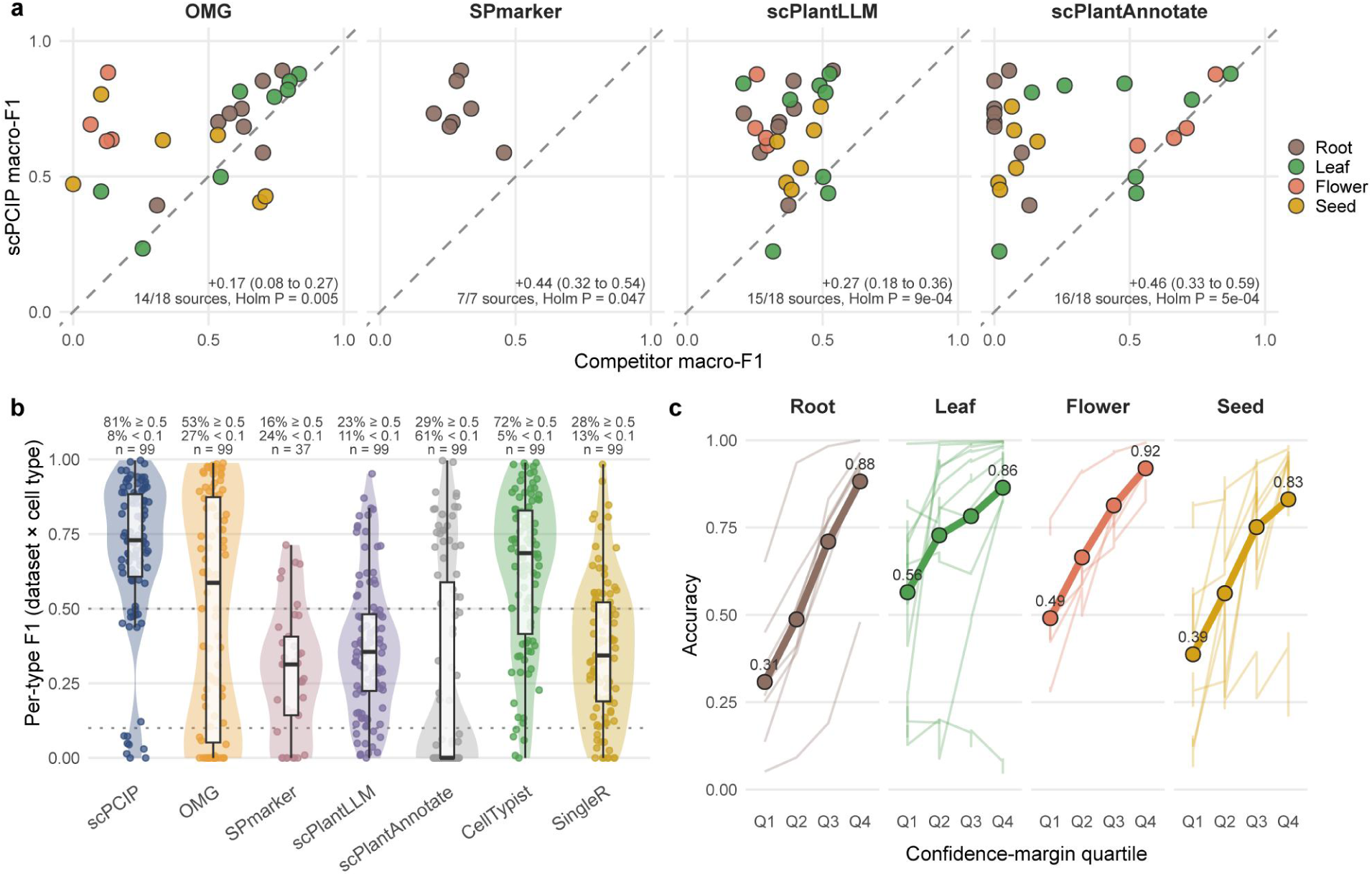
Benchmark against reference-free methods. a, Macro-F1 of scPCIP against each method, one dot per dataset; dashed line, equality; insets, mean difference with 95% bootstrap CI and Holm-corrected Wilcoxon *P*. b, Per-type F1 of every dataset–cell-type pair; boxes, median and interquartile range; whiskers, 1.5 × interquartile range; dotted lines, F1 = 0.5 and 0.1. c, Accuracy by confidence-margin quartile; thin lines, datasets; thick line, mean.

**Extended Data Fig. 4.**
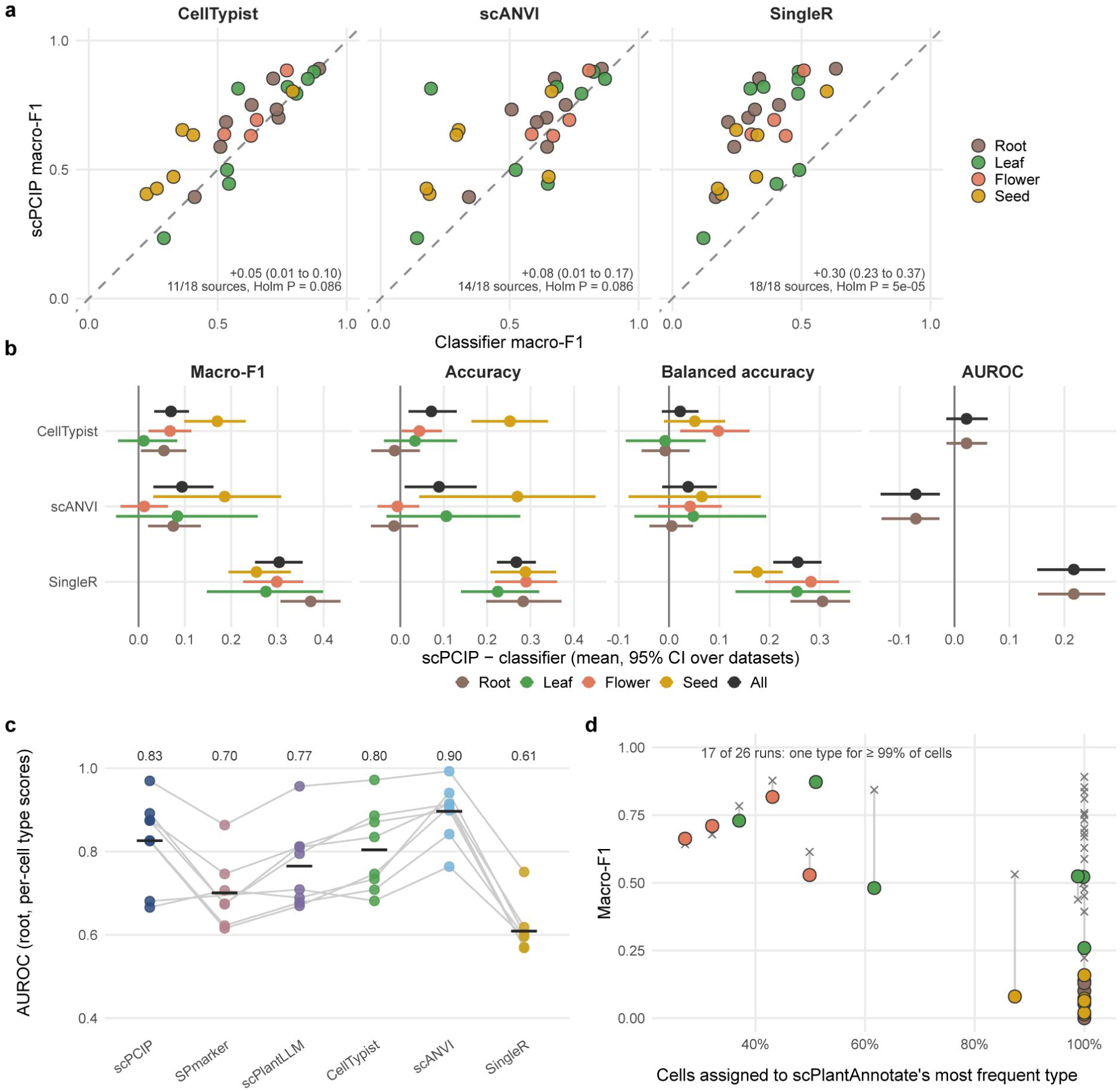
Comparison with supervised classifiers. a, Macro-F1 of scPCIP against CellTypist, scANVI and SingleR, one dot per dataset; dashed line, equality; insets, mean difference with 95% bootstrap CI and Holm-corrected Wilcoxon *P*. b, Difference in macro-F1, accuracy, balanced accuracy and AUROC per organ. c, Per-type AUROC in root datasets; points, datasets; bars and numbers, means. d, Share of cells in the most frequent scPlantAnnotate type against macro-F1; circles, scPlantAnnotate; ×, scPCIP on the same dataset.

**Extended Data Fig. 5.**
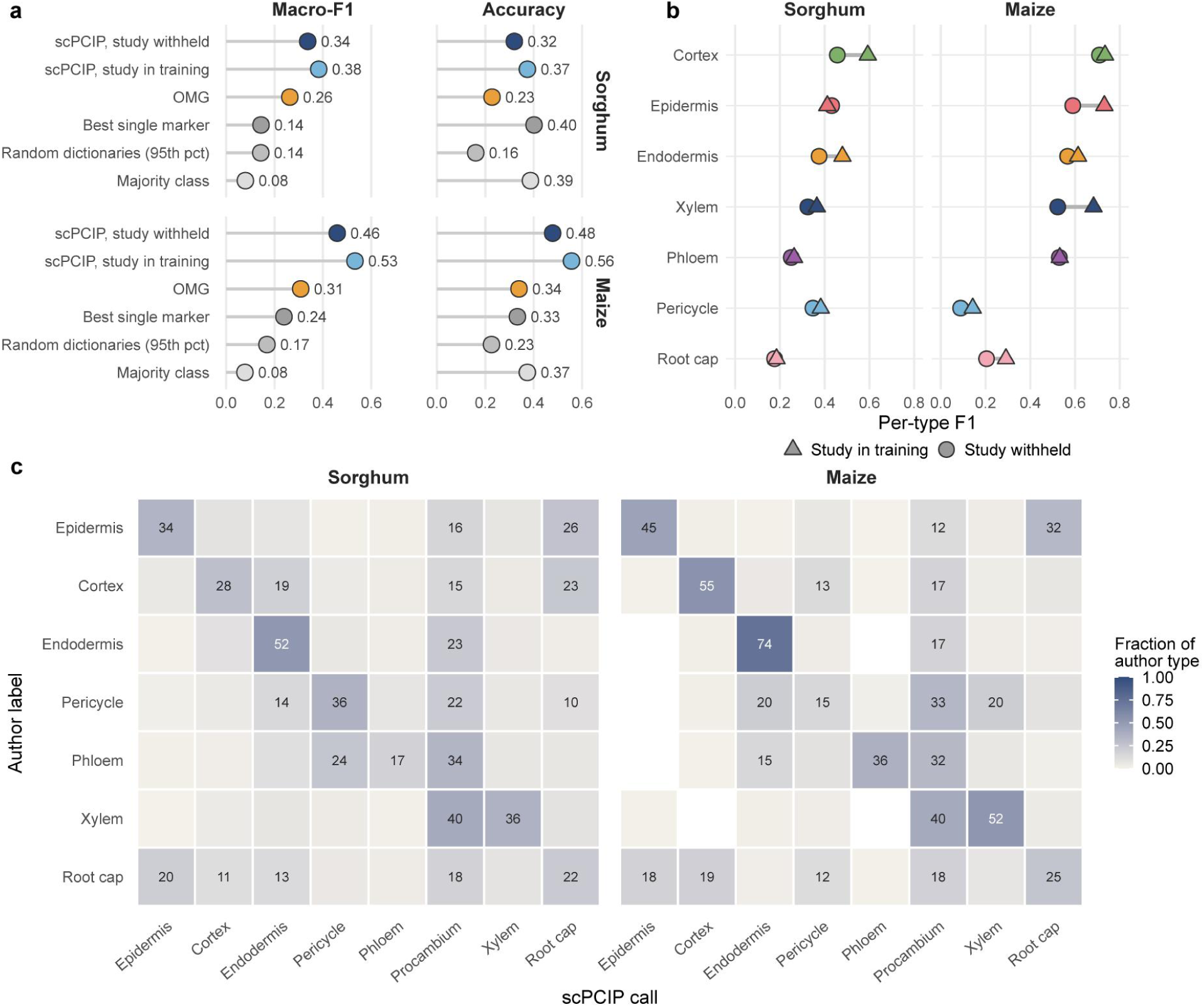
Transfer to a withheld study. a, Macro-F1 and accuracy on sorghum and maize with the study withheld. b, Per-type F1 with the study withheld or included. c, Confusion matrices.

**Extended Data Fig. 6.**
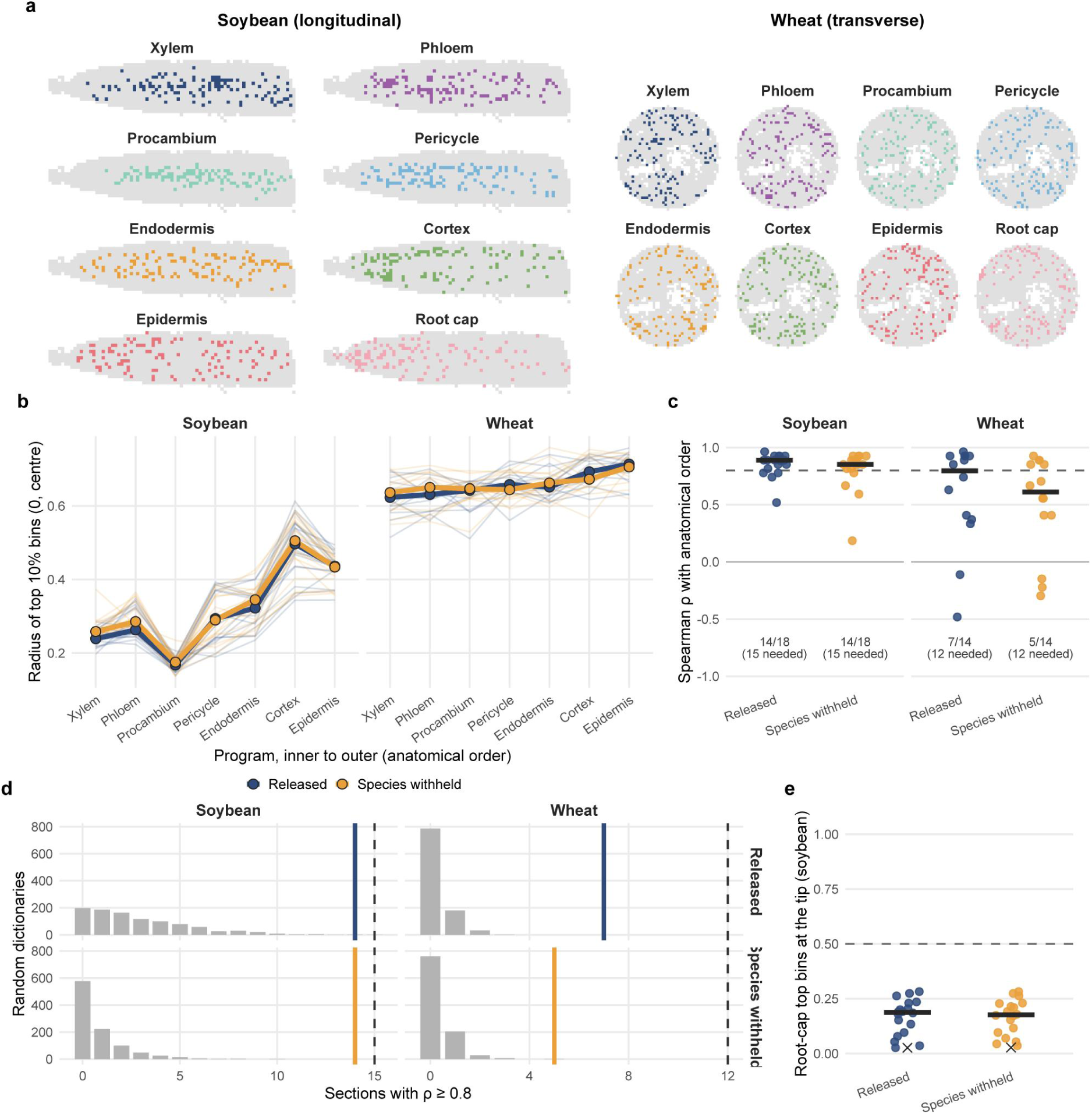
Label-free radial order of root programs. a, Bins in the top 10% for each program, scored with the species-withheld dictionary; grey, other bins. b, Mean radius of the top 10% bins per program; thin lines, sections; thick lines, median. c, Spearman ρ between observed and anatomical program order per section; bars, medians; dashed line, pre-specified threshold. d, Number of sections with ρ ≥ 0.8; grey, random dictionaries; colour, observed; dashed line, required number. e, Share of top root-cap bins at the tip; bars, medians; dashed line, required proportion; ×, null 95th percentile.

**Extended Data Fig. 7.**
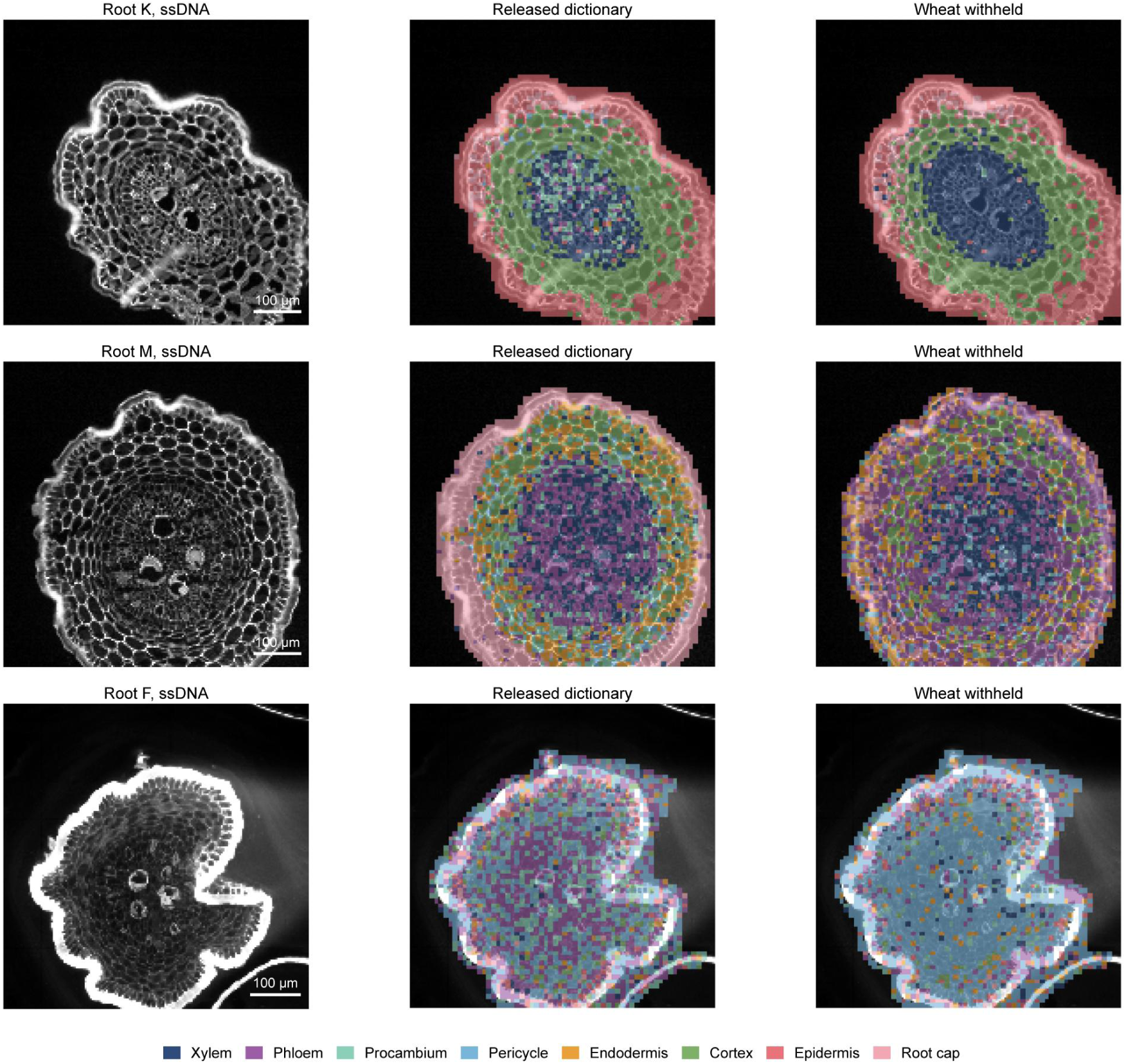
Wheat roots on the tissue image. Transverse wheat root sections. Left, ssDNA image; middle, scPCIP calls of 10-µm bins within the tissue mask from the released dictionary; right, calls from the wheat-withheld dictionary. Scale bar, 100 µm.

**Extended Data Fig. 8.**
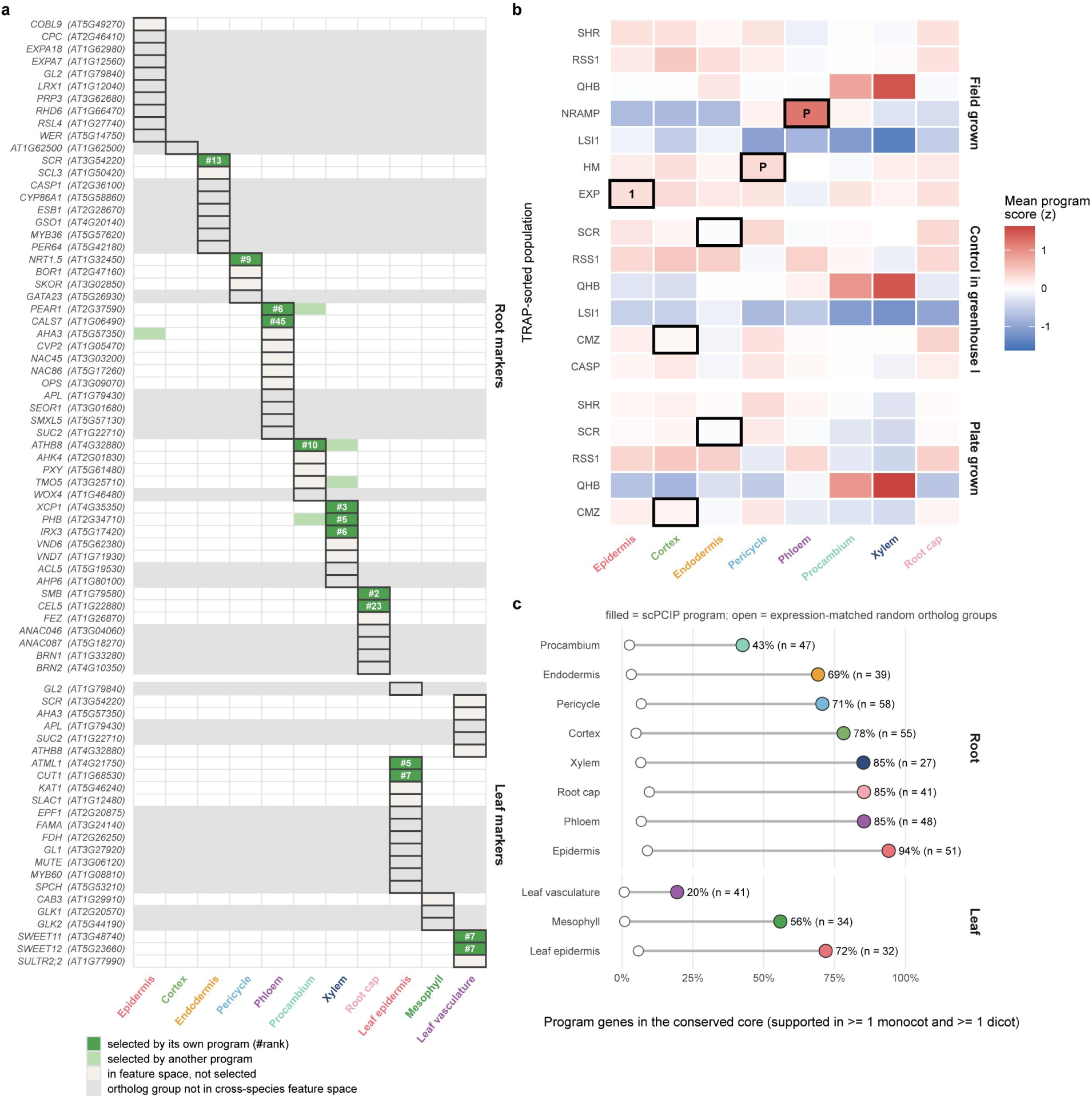
Classic markers, TRAP populations and conserved cores. a, Fate of classic *Arabidopsis* markers in the root and leaf programs. b, Mean program score of rice root TRAP populations; outlined, target population; P, ranked first with *P* < 0.05 against 1,000 random programs; 1, ranked first only. c, Conserved-core share of each program.

**Extended Data Fig. 9.**
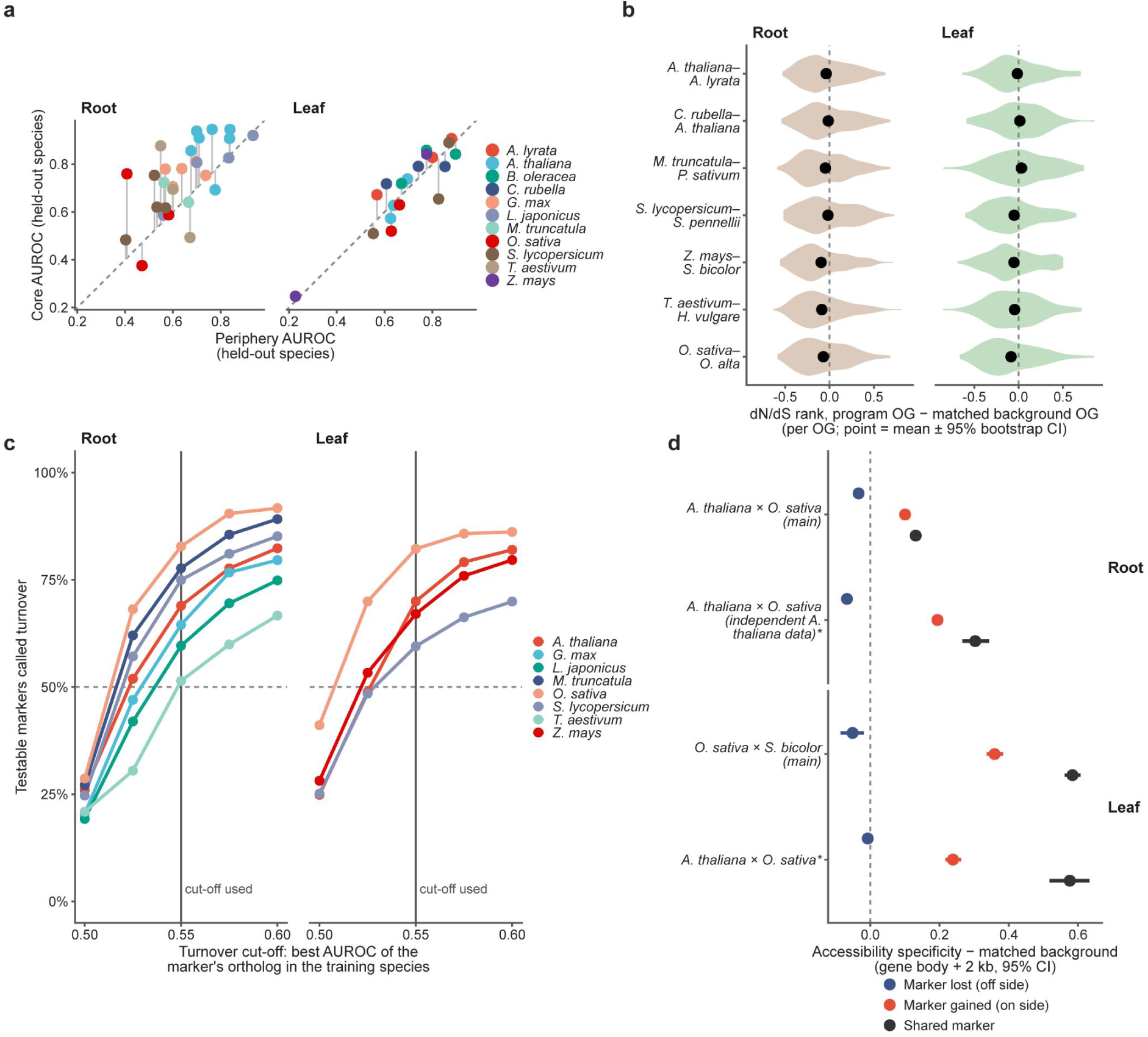
Cores, selection, turnover and accessibility. a, AUROC of the conserved core against the periphery per species–cell-type unit; dashed line, equality. b, dN/dS rank difference from matched background per sister-species pair. c, Share of turnover markers as a function of the cut-off. d, Accessibility of lost, gained and shared markers relative to matched background; *, replication with independent *A. thaliana* data.

**Extended Data Fig. 10.**
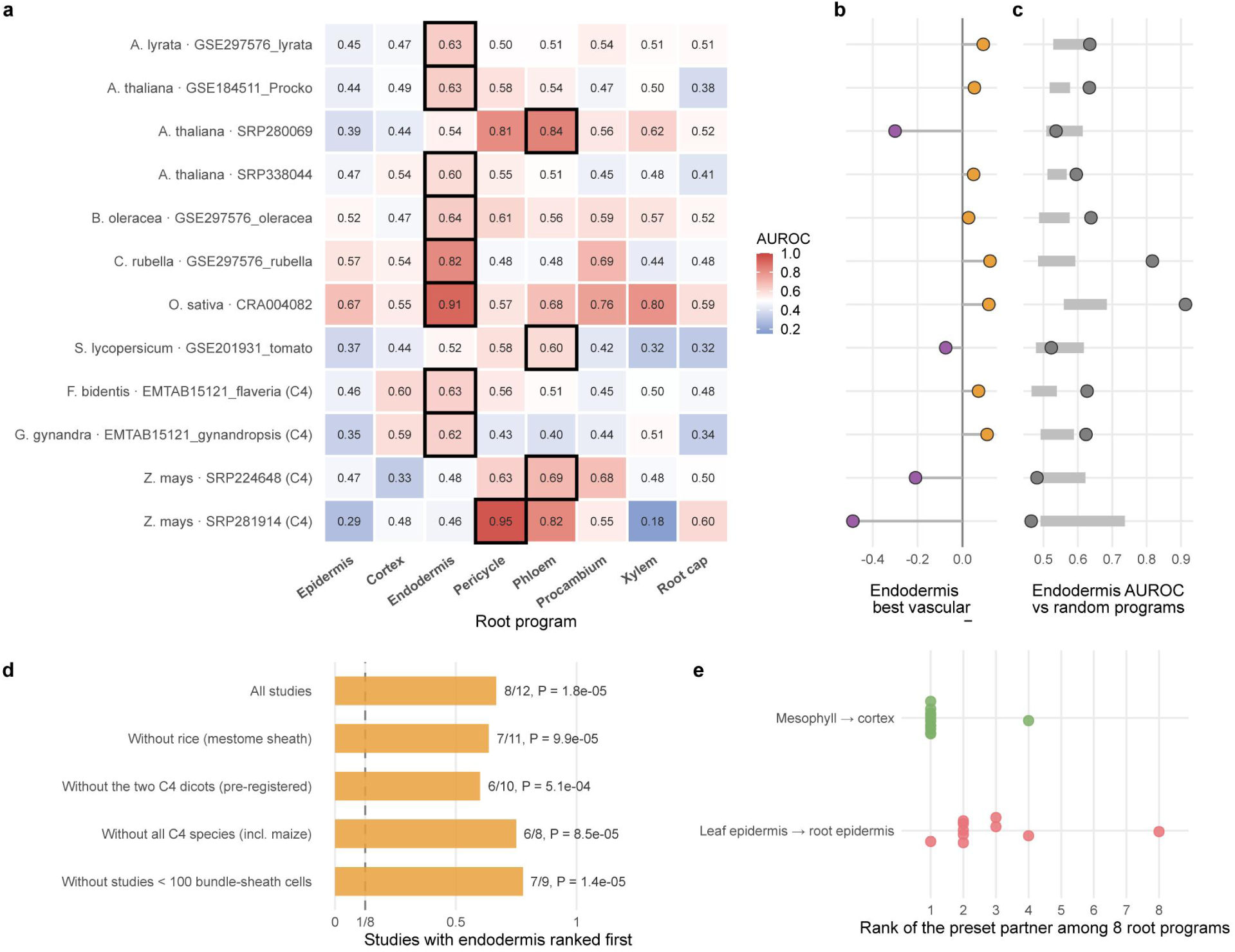
Program-level test of bundle-sheath cells with the root dictionary. Leaf cells of 12 species-by-study datasets from nine species, scored with the released root dictionary; studies are labelled in the panels. a, AUROC of each root program for bundle-sheath cells against all other leaf cells of the same dataset; outlined, highest program. b, Endodermis AUROC minus that of the best of the pericycle, phloem, procambium and xylem programs per dataset; orange, endodermis higher; purple, a vascular program higher; one-sided Wilcoxon signed-rank test. c, Endodermis AUROC against 1,000 detection-matched random gene sets; dots, endodermis, filled when above the 95th percentile; bars, median to 95th percentile of the random sets. d, Datasets in which endodermis ranked first among eight root programs; dashed line, 1 in 8 expected by chance; one-sided binomial *P*. e, Rank of the pre-specified partner program among eight root programs.

