## Supplementary material for "Cross-species gene programs enable reference-free annotation of plant cells in single-cell and spatial transcriptomes": scPCIP_Supplementary_Information

### Supplementary Methods

### Training data and cell-type labels

We assembled published single-cell and single-nucleus RNA-seq datasets of root, leaf, flower and seed with cell-type annotations from GEO, CNGB/STOmicsDB[53], the authors or the re-processed collection of iPscDB[33] (Supplementary Table 2). Training labels came from the original studies or from the iPscDB re-annotation, which supplied the labels of 22 of the 31 root and 22 of the 26 leaf training datasets, 44 of 68 in total. One sorghum root dataset, with BioProject accession PRJNA935359, used scPlantDB-provided labels, and one tomato leaf dataset, with GEO accession GSE201931, used labels recovered from marker expression. All flower and seed labels came from the original studies.

Root labels were mapped to the Plant Ontology of Cell Types (POCT)[14] and from there to eight coarse types: epidermis, cortex, endodermis, pericycle, phloem, procambium, xylem and root cap. Where possible, they were also mapped to fine types. Exodermis was kept as a fine type within cortex, meristematic and quiescent-centre initials were kept as a state, and fine types with fewer than 150 cells were not modelled. Leaf labels were collapsed into epidermis, mesophyll and vasculature, with guard cells included in epidermis and bundle sheath, xylem and phloem included in vasculature; maize bundle-sheath labels were excluded from training. Flower labels for pollen, stamen, carpel, perianth and non-floral tissue, and seed labels for embryo, endosperm and seed coat, were mapped with fixed tables (Supplementary Table 2c).

The units used throughout are as follows. A dataset is one annotated expression matrix. A study is one publication or data release, and a source is a study–organ combination. In the benchmark, 26 held-out datasets formed 18 sources from 16 independent studies. In the bundle-sheath analyses, a species-by-study dataset is one species within one study, and a study block pools the species of a multi-species study. Section-level statistics in spatial data treat tissue sections as units and do not represent biological replication.

### Ortholog-group features

Proteomes were annotated with eggNOG-mapper v2.1.13 against eggNOG 5.0.2[54,55] at the Viridiplantae level, taxon 33090, keeping one ortholog group per gene by choosing the highest-scoring assignment across isoforms. The feature value of ortholog group $o$ in cell $c$ is the summed raw count of its member genes,

$$x_{c,o} = \sum_{g \in o} n_{c,g}. \qquad (1)$$

Where normalized values were needed for refinement, marker statistics and plots, we used $\log(1 + 10^4 x_{c,o} / \sum_{o'} x_{c,o'})$, which denotes log CP10k. Before the root, leaf and flower dictionaries were trained, ortholog groups whose eggNOG descriptions matched terms for ribosomal proteins, translation initiation or elongation, histones, cyclins or kinesins, heat-shock proteins or DNA replication were removed: 446 groups for the root, 764 for the leaf and 626 for the flower dictionary (Supplementary Table 1). Matching used all descriptions of a group, so some removed groups have other functions. The seed dictionary was trained without this filter, and query data are never filtered.

### Learning cell-identity programs

Training cells were subsampled per dataset and label, with up to 250 per coarse-and-fine label for the root and up to 500 per coarse label for other organs. They were restricted to ortholog groups present in all training species and converted to within-cell rank percentiles $p_{c,o}$, with average ranks for ties, including zeros.

Programs were learned for each cell type $t$ against all others. Cells were grouped into stability groups by species for root and by dataset for the other organs. For group $d$, the contrast of feature $f$ is

$$C_d(f) = \bar{p}^{(t)}_{d,f} - \frac{1}{|T_d| - 1} \sum_{r \in T_d \setminus \{t\}} \bar{p}^{(r)}_{d,f}, \qquad (2)$$

where $\bar{p}^{(r)}_{d,f}$ is the mean rank percentile of $f$ in type $r$ of group $d$. Features with $C_d(f) > 0$ in at least two groups and a positive median contrast were ranked by the median, and the top 300 for root and seed or 1,000 for leaf and flower became candidates. On the candidates, we ran stability selection[5 6]: in each of 30 rounds, one stability group was left out at random when at least three contained type $t$, 400 cells per type were resampled, and an L1-penalized logistic regression of type $t$ versus the rest was fitted using scikit-learn[5 7]. A feature entered the program if it was selected in at least 60% of rounds for root and seed or 40% for leaf and flower with a positive median coefficient $\widetilde{\beta}_f$. Programs were capped at 60 features, and weights were

$$w_f = \widetilde{\beta}_f / \max_{f' \in F_t} \widetilde{\beta}_{f'}. \qquad (3)$$

The 19 coarse programs contained 25–60 ortholog groups, with 339, 107, 280 and 79 distinct groups in the root, leaf, flower and seed dictionaries, respectively, 835 program–group entries and 727 distinct groups overall (Supplementary Table 3). Fine root programs were learned within each coarse type, and xylem and phloem programs were learned within leaf vasculature and reported as a secondary subtype only. The released dictionaries were trained on all data: 65,809 root cells from 31 datasets of 9 species, 30,745 leaf cells from 26 datasets of 11 species, 6,926 flower cells from 4 datasets of 4 species and 9,425 seed cells from 7 datasets of 3 species (Supplementary Table 16).

An exploratory stem dictionary was evaluated without release by leave-one-species-out across three author-labelled stem datasets (*Arabidopsis* inflorescence stem, soybean hypocotyl and rice culm), each fold trained on the other two with the development stem recipe of 1,000 candidates and a selection-frequency threshold of 0.4. Its five classes were cambium, epidermis, ground tissue, trichome and vasculature (Supplementary Table 2). It was not used in the spatial analyses and will be released when more stem data become available (Supplementary Note 3).

### Scoring and classification

In a query cell, ortholog groups are ranked over all features present as $\pi_{c,o} \in [0,1]$, and the score of program $p$ with features $F_p$ is the weighted mean rank of the features found in the query, $Q$:

$$s_{c,p} = \frac{\sum_{f \in F_p \cap Q} w_f \ \pi_{c,f}}{\sum_{f \in F_p \cap Q} w_f}. \qquad (4)$$

The type score $s_{c,t}$ is the maximum over the programs of type $t$. Because the typical rank of a program differs between datasets, type scores are centred on the cells that the raw argmax did not assign to that type,

$$\tilde{s}_{c,t} = s_{c,t} - median\{s_{c',t} : \hat{y}_{c'}^{raw} \neq t\}, \qquad (5)$$

and each cell receives the call and confidence margin

$$\hat{y}_c = \operatorname{argmax}_t \tilde{s}_{c,t}, \qquad m_c = \tilde{s}_{c,\hat{y}_c} - \max_{t \neq \hat{y}_c} \tilde{s}_{c,t}. \qquad (6)$$

Every cell receives a call; the margin, all type scores and a confidence quartile with its expected accuracy are reported alongside the call. The margin is not a calibrated probability.

Calls are refined once. Cells at or above the 75th margin percentile of their called type are used to train a logistic regression on the first 30 principal components of the query, calculated from log CP10k values for the 2,000 most variable features, which relabels all cells; each Leiden[58] cluster calculated with Scanpy[59] at resolution 2.0 whose majority label covers at least half of its cells then takes that label. Final calls therefore depend on variable features of the query as well as on program members. Because refinement depends on the number of linear-algebra threads, the software fixes BLAS and OpenMP to one thread, and all analyses were run this way. In a test on 3,426 maize root cells, multithreading changed 0.93% of refined calls and none of the unrefined ones.

For each called type, scPCIP reports the program's ortholog groups with their weights, the query genes in each group, their detection and mean expression inside and outside the type, and $\log_2$ fold changes (Extended Data Fig. 2). When a sample column and an integrated-cluster column are supplied, a reproducibility diagnostic compares call distributions of matched clusters across samples by total-variation distance: consistent at ≤ 0.20, to be checked at ≤ 0.30 and otherwise not reproducible. Reliability estimates stored with each dictionary comprise per-type F1 and accuracy by margin quartile and come from leaving out each training species in turn.

### Evaluation

**Designs.** For leaf, flower and seed, each of the 7 leaf, 4 flower and 3 seed species was held out in turn and scored by a dictionary trained on all other species, with three training seeds, yielding 18 test datasets. For root, eight datasets not used in training were scored by the released root dictionary. Except for *Lotus japonicus*, the species in these datasets were represented in training by other datasets, so this arm primarily tests transfer across datasets. Root cross-species performance was estimated separately by leaving out each of seven root species. Up to 6,000 labelled cells per test dataset were scored by every method. These folds were also used during development to select training recipes and prediction settings for root, leaf and flower.

**Evaluation labels and exclusions.** Evaluation labels came from the original studies, except that the labels of five leaf datasets with accessions CRA004082, SRP224648, SRP280069, SRP281914 and SRP338044, and the *Medicago* root dataset were obtained through scPlantDB curation (Supplementary Table 4). Two leaf datasets were excluded from evaluation: tomato GSE201931, whose labels were marker-recovered, and rice SRP286275, which was removed after results were examined because marker expression did not support its labels. In the root benchmark, selected labels identified as inconsistent with marker evidence were excluded for all methods: rice epidermis; wheat pericycle, phloem and root cap; and tomato xylem; these checks were prompted by low scPCIP scores and made before the competitors were run.

**Metrics.** Macro-F1, the unweighted mean of per-type F1 over types with at least 20 test cells, was the primary measure[60]; we also report accuracy, balanced accuracy and the mean one-versus-rest AUROC for the eight root datasets, where per-cell scores were available. Cells whose labels were outside the dictionary's types were not evaluated; predictions outside the type set counted as errors. Accuracy by margin quartile was evaluated in the root leave-one-species-out folds.

**Statistics.** The primary unit was the source, with 18 sources. For each competitor we computed per-source mean differences in macro-F1, their mean with a percentile bootstrap 95% CI from 10,000 resamples of sources, the number of sources favouring scPCIP and a two-sided Wilcoxon signed-rank test, with Holm correction[61] across the seven competitors. The ten leave-one-species-out sources were tested in the same way, with Holm correction across six competitors because SPmarker applies only to root. To account for studies contributing more than one source, we repeated the comparison on 16 independent studies by averaging all datasets within a study, with Holm correction across seven competitors (Supplementary Table 5). Source-level accuracy and the sensitivity analyses retaining SRP286275 or excluding the six datasets with scPlantDB-curated labels were corrected across the seven competitors in the same way. Organ-specific and dataset-level comparisons are reported without multiplicity correction. Dataset-level paired differences were analysed in the same way (Extended Data Fig. 3 and Supplementary Table 4). Per-type F1, set to 0 for a type a method never predicted, was compared between scPCIP and each method over shared dataset–cell-type pairs by a two-sided Wilcoxon signed-rank test after discarding zero differences and without multiplicity correction; these comparisons were treated as descriptive because pairs are nested within datasets and sources. Divergence times were median TimeTree[62] estimates to the closest training species, with groups below 25, at 50–60 and above 100 million years. No equivalence test was performed. All statistics used SciPy 1.17.1.

**Competing methods.** All methods were scored on the same test cells with the settings recorded for each method (Supplementary Table 4); deviations from defaults are stated here.

- *OMG*[17] v0.1.0: clustering with Seurat[63] v5.5.0 in R 4.5.3 at resolutions 0.3, 0.5 and 1.0, positive cluster markers, and OMG with its bundled reference and defaults. Orthogroups for species absent from the bundle were built with OrthoFinder[64] v2.5.5 from the OMG reference proteomes. For each dataset we report the resolution most favourable to OMG by macro-F1. The bundled reference included several query species, so this evaluation was not strictly leave-one-species-out.
- *SPmarker*[16] baseline: the published *Arabidopsis* root markers, mapped to ortholog groups and scored as unweighted programs with the scPCIP scorer, because SPmarker has no cross-species implementation. Pericycle and procambium could not be predicted. Tomato SlarAR was

excluded because its labels had been assigned with SPmarker markers, leaving seven root sources. Soft voting across the marker sets was tested as a sensitivity analysis.

- *scPlantLLM*[26] : the authors' tutorial notebook and preprocessing were run as released, changing only paths, the number of classes and the GPU; the pipeline comprised masked-language-model pretraining, loading of the released pretrained weights, classification training and prediction fine-tuning, each with the single epoch set in the tutorial. Its gene vocabulary is *Arabidopsis*-based, so each ortholog group was represented by its member in that vocabulary with the highest mean expression in the *Arabidopsis* datasets of the organ, and groups without such a member were dropped. Zero-shot use was not possible because the zero-shot head covers only *Arabidopsis* root types.
- *scPlantAnnotate*[27] at commit dc7d7b1: the released code; runs that assigned nearly all cells to one type were kept.
- For scPlantLLM and scPlantAnnotate, training used the same cells, sampled from each training unit in proportion to label up to 10,000 cells; root and leaf units were capped, whereas flower and seed units were used in full.
- *CellTypist*[18] v1.7.1, *SingleR*[20] v2.12.0 and *scANVI*[19],[65] using scvi-tools v1.1.6.post2 with scArches[66] query mapping: trained on the same leaf, flower and seed cells and labels as the corresponding scPCIP dictionary, using several thousand ortholog-group features and without the non-identity filter. The primary root training set contained 62,840 rather than 65,809 cells because exodermis and meristem labels were not included. For CellTypist, the better of annotation with or without majority voting on the test labels was reported, as pre-specified; SingleR used the "classic" marker method because the pre-specified "wilcox" option could not be installed.

**Unified root training cells.** CellTypist, scANVI and SingleR were retrained on the 65,809 root cells used by scPCIP, with exodermis merged into cortex and meristem kept as a ninth class used only during training; meristem predictions were reassigned to the highest-scoring of the eight evaluated types. Statistics were recomputed at source and study level as above. scPlantLLM and scPlantAnnotate were not rerun. These criteria were written before computation.

**Re-evaluation on eight further sources.** Eight study–organ sources from six studies outside the training data had been set aside on 14 October 2026 for a pre-specified confirmatory comparison using earlier dictionaries: root, leaf and flower of CNP0002614; GSE224539, GSE240098 and GSE303996 in root; and GSE273033 and GSE313726 in leaf. Their results had been examined before the dictionaries were finalized, so the re-evaluation of the released dictionaries was not blind. The criteria for this re-evaluation were written before it was computed. CNP0002614 counts were the authors' corrected non-integer values, given identically to all methods. As pre-specified, the tests were one-sided Wilcoxon signed-rank tests for scPCIP greater than CellTypist and scANVI with Holm correction across these two comparisons, and were repeated with the three CNP0002614 organs merged into one source; SingleR and OMG were reported descriptively. OMG terminated with an error on GSE240098.

**Independent study.** The GSE225118 study[6], a related sorghum project and an iPscDB maize dataset sharing 2,408 barcode cores with it were removed from root training, the dictionary was retrained under criteria fixed beforehand, and 6,000 sorghum cells and 4,944 maize cells were scored. Baselines were the majority class, the best single ortholog group per type as determined by training AUROC and 20 random dictionaries in which each feature was replaced by one from the same expression decile. Differences from OMG used cell-level bootstrap 95% CIs. The dictionary containing the study was scored on the same cells to estimate the effect of withholding it. The pre-specified primary measure was accuracy, with support defined as exceeding the best single marker in both species; macro-F1 is reported as primary to match the main benchmark, a choice made after seeing both.

### Spatial transcriptomics

Program and annotation tests used seven datasets: Stereo-seq of soybean root tips and leaf[4], Visium of soybean seed[5], Stereo-seq of rice root, leaf and seed[3] and single-cell Stereo-seq of *Arabidopsis* leaf[34]. The radial analysis also used Stereo-seq of transverse wheat roots[7], with accessions given in Supplementary Table 6. Integer counts were reconstructed where only log-normalized values were distributed. Each dataset was scored with a dictionary trained without its species; soybean leaf was scored with the released leaf dictionary, which contains no soybean. Soybean stem and hypocotyl sections from the same studies were not analysed, because no stem dictionary is released.

**Program test.** In each section, the score of each program was used to separate spots that the authors had assigned to the program's tissue from all other labelled spots, and AUROCs were averaged over sections, with bootstrap 95% CIs for datasets with at least three sections. The null was 100 random programs per section with each feature replaced by one from the same expression decile, keeping weights. Empirical $P$ values were corrected by the Benjamini–Hochberg procedure[6, 7] across the 31 tests, and a program met the criteria if $q < 0.05$ and AUROC > 0.5. Spot-level ROC curves used spots pooled across sections with DeLong 95% CIs (Fig. 3c). The rice seed curve pooled 16 sections, including fertilization and germination samples, whereas other rice seed panels used 10.

**Annotation test.** Spots were called as in the released software, including refinement. Macro-F1 was computed over spots pooled across sections and compared with a majority-class baseline defined by the most frequent label across sections and with 20 expression-matched random dictionaries per section, processed with the same refinement and pooled by draw; a dataset passed if scPCIP exceeded both the majority class and the 95th percentile of the random dictionaries. Soybean root was analysed in the radial test and not included, leaving six datasets. As sensitivity analyses, the comparison was repeated without refinement for both scPCIP and the random dictionaries, and on per-section means. For soybean seed and *Arabidopsis* leaf, a supervised within-dataset reference was computed by leave-one-section-out: an L2-regularized logistic regression with C = 0.1 in scikit-learn on z-scored log CP10k ortholog-group expression was trained on the author labels of all other sections and evaluated on the held-out section, and macro-F1 was computed over the pooled held-out spots.

**Radial order.** Each bin received a relative radial position $r_i \in [0,1]$: for soybean, the distance from the midline of the section's long axis, normalized within slabs along that axis; for wheat, the distance from a manually placed root centre normalized by the root radius. The characteristic position of type $t$ in a section is the mean position of its top-scoring decile of bins,

$$\bar{r}_t = mean\{r_i : s_{i,t} \geq Q_{0.9}(s_{\cdot,t})\}, \qquad (7)$$

and agreement with anatomy is the Spearman correlation between $\bar{r}_t$ and the expected order of vascular types, pericycle, endodermis, cortex and epidermis. Pre-specified criteria (Supplementary Table 6) required ρ ≥ 0.8 in at least five-sixths of sections, a one-sided sign test that ρ is positive across sections against a null probability of 0.5, more sections with ρ ≥ 0.8 than the 95th percentile of 1,000 random dictionaries matched by mean-expression decile and, in soybean, a median of at least 50% of the top-decile root-cap bins located in the terminal 10% of the section's long axis. Tip localization could not be assessed in wheat transverse sections. Author labels were not used to infer radial position. The pre-specified soybean test of all 24 sections used an earlier dictionary and did not pass; restricting the test to the 18 sections with at least five author-labelled stele bins to define the expected order and updating to the current dictionary were introduced after that result. The comparison of median ρ with random dictionaries by a one-sided empirical *P* was not pre-specified (Fig. 3e). For the wheat type maps, tissue was segmented from the authors' registered single-stranded DNA image, and bins with at least 50 transcripts were scored, one chip per query, with the released root dictionary and with the dictionary of the wheat-held-out fold (Fig. 3b and Extended Data Fig. 7). The 14 wheat sections came from two chips.

### Interpretability analyses

**Co-expression layout.** Pseudobulk profiles of the root training data, calculated per dataset and cell type, were centred within datasets, and correlations between program ortholog groups were embedded with UMAP[6,8]. Because programs were selected for specificity on the same pseudobulks, the layout is descriptive and no statistic is derived from it.

**Classic markers.** We curated 76 records of *Arabidopsis* markers from 70 genes, comprising 53 root and 23 leaf records, from primary literature without reference to the dictionaries (Supplementary Table 7). A record was testable if its ortholog group lay in the shared training feature space and was counted as selected if that group belonged to the program of its cell type or, in a separate analysis, to any other program.

**Rice TRAP.** Translatomes of rice root cell populations[36] , with three replicates per population, were summed into ortholog groups, log-transformed and z-scored within growth batches, and programs of a dictionary trained without rice were scored as weighted means. A program was supported if its target population scored highest and its target-versus-other AUROC exceeded that of 1,000 expression-matched random programs at $P < 0.05$ (Supplementary Table 8). We also report Bonferroni correction across the five evaluable programs. The field-grown and greenhouse-control batches were separate primary batches, and the plate-grown batch was a replication.

**Conserved core.** An ortholog group supports a program in a training dataset if its one-versus-rest AUROC is at least 0.6; groups supported by at least one monocot and one dicot species form the conserved core (Supplementary Table 9). The same definition was applied to expression-matched random ortholog groups; because program members were selected for discrimination, this comparison is descriptive.

### Conservation analyses

Six confirmatory hypotheses were specified in an internal hash-stamped protocol before analysis and Holm-corrected together (Supplementary Table 10). For H5a, core and periphery were re-derived within each leave-one-species-out fold across 11 held-out species and 47 species–program units with at least five core and five periphery groups. The larger set was subsampled 50 times to the size of the smaller, each subset was scored with the fold weights, and AUROCs were averaged over the species' datasets. The test statistic is

$$\bar{\Delta} = \frac{1}{S}\sum_{s=1}^{S}\frac{1}{|U_s|}\sum_{u\in U_s}\left(\bar{A}_u^{core} - \bar{A}_u^{peri}\right), \qquad (8)$$

over the $S = 11$ species with units $U_s$, tested with an exact one-sided sign-flip test over species. Organ-specific and endodermis-excluded values are descriptive summaries of the same units.

The pre-specified secondary control compared core groups with nearest-neighbour groups of the same training discrimination from the non-program pool; it could not be interpreted because matched sets were unbalanced in 46 of 47 units. In an exploratory analysis, core and peripheral groups were instead matched on expression, detection, family size and training AUROC using

Mahalanobis cost on percentile-ranked, z-scored covariates, one-to-one assignment, a caliper of 0.25 and at least five pairs per unit, with a species-level bootstrap CI from 10,000 resamples.

Gene ages for H1 and H2 came from hierarchical orthogroups from OrthoFinder[64] v3.1.5 across 61 species, with background groups matched on expression, detection, family size and protein length. For dN/dS, reciprocal-best-hit orthologues of seven sister-species pairs identified using PLAZA 5.0[69], DIAMOND[70] and MAFFT[71] were analysed with yn00[72,73]. Each ortholog group received the median ω of its gene pairs and a percentile rank within the species pair, and program groups were compared with background groups matched for expression, family size and protein length, with bootstrap CIs across program groups (Supplementary Table 11).

Marker classification used the top 100 markers per cell type in each held-out dataset. Records were testable if their ortholog groups could be evaluated in the training species, and program-selected markers were classified separately. Other records were classified by the AUROC of their ortholog group averaged within and then across training species: turnover at < 0.55, uncertain at ≥ 0.55 and < 0.60, missed at ≥ 0.60, and shared markers at ≥ 0.60 regardless of program membership. Thresholds were varied from 0.50 to 0.60 (Supplementary Table 12). “Turnover” is an operational label. For a between-study baseline written before computation, records that could also be scored in another study of the same species were compared at thresholds of 0.55 and 0.50. Cross-species and within-species proportions, regardless of program membership, were balanced across species, with species-level bootstrap CIs from 10,000 resamples, and a sensitivity analysis assessed each training species separately (Supplementary Table 5).

Co-expression neighbourhood conservation was the AUROC with which a group’s correlation ranking in a training dataset recovers its 50 strongest partners in the held-out dataset, minus the mean AUROC of reference non-marker groups matched by detection decile in both datasets. A sensitivity analysis excluded program-selected markers from the shared set. For chromatin, gene activity, defined as peaks over the gene body plus 2 kb upstream summed across member genes, and RNA specificity were compared between species in single-nucleus multiome data of *Arabidopsis*[38] and rice[2] roots and rice and sorghum leaves[8]. Pairs were selected on RNA specificity alone, requiring RNA detection ≥ 5 counts per million in both species and $\log_2$ specificity ≥ 1 in one species and ≤ 0.25 in the other. Remaining pairs passing the same expression filter formed the background, which was also reweighted to the decile distribution of the absolute RNA difference of the selected pairs, with ortholog-group bootstrap CIs from 2,000 resamples. These analyses were exploratory.

### Leaf–root correspondence

**Marker overlap.** Markers per cell type were ortholog groups with a mean $\log_2$ specificity of at least 0.5, positive in at least 70% of study blocks, within a universe of 11,675 groups detected in more than 1% of cells in at least half of the blocks in each organ. Marker discovery required at least three blocks and held-out evaluation required at least five markers; overlap was tested with Fisher's exact test (Supplementary Table 13). The 12 leaf datasets of the program test formed 9 study blocks, because the three species of GSE297576 and the two species of E-MTAB-15121 were pooled by study. Epidermis included guard cells, and senescent, dividing, stressed and precursor states were excluded. Each of the 9 leaf blocks and 11 root blocks was held out in turn: in the held-out block, each marker set of the other organ was scored by AUROC on that block's type specificity, a hit was recorded when the hypothesized partner ranked first, and the number of hits was tested against fold-specific chance rates using a Poisson-binomial tail. Discovery vascular markers were removed from non-vascular types for the held-out test and for the odds ratios (Fig. 6b and Supplementary Table 13d). Without this filter, bundle-sheath markers still overlapped most with pericycle and endodermis markers, with odds ratios of 20.3 and 12.5, against 4.6 for root vascular markers (Supplementary Table 13c). *P* values were Holm-adjusted over the four leaf rows; the same rows were confirmed when adjusting over all nine leaf and root rows, the family used for the reverse direction (Supplementary Table 15k). iPscDB blocks were evaluated separately as supplementary evidence (Supplementary Table 13a). Labels for GSE184511 and GSE201931 were marker-recovered; without these two blocks, bundle-sheath cells matched the inner root layers in 6 of the remaining 7 (Supplementary Table 13b).

**Family presence and sequence constraint.** Marker-family presence used OrthoFinder v3.1.5 orthogroup gene counts over 89 species, anchored on *A. thaliana*, with an orthogroup counted as present when a species had at least one copy. Rice, which was not among the 89, was added from eggNOG membership. For the sequence-constraint comparison (Fig. 6d), the ω percentile rank from yn00 was modelled per gene, with shared-marker status, marker strength, tissue system and log leaf and root expression as covariates, in *A. thaliana*–*A. lyrata*, *Z. mays*–*S. bicolor* and *S. lycopersicum*–*S. pennellii*. Significance used two-sided permutation of shared status within system and strength-tertile strata with 5,000 permutations, and CIs used a stratified gene bootstrap with 2,000 resamples (Supplementary Fig. 16).

**Program-level test.** The program-level test followed criteria frozen before computation. For 12 leaf species-by-study datasets from 9 species with labelled bundle sheath (Supplementary Table 14), all cells were scored with the released software and root dictionary, and the AUROC for bundle sheath versus all other cells was computed before refinement from the exported type scores, which are the weighted-rank program scores centred as in equation (5). Centring subtracts one constant per type within a dataset, so these AUROCs and the ranking of programs equal those from uncentred scores; the largest difference across the 12 datasets and eight programs was $5 \times 10^{-9}$. The primary statistic was the number of datasets in which endodermis ranked first, tested with a one-sided binomial test against a chance of 1/8; species were ranked by the median over their datasets. Because datasets share species and studies, we also report non-overlapping study blocks and lineages, using the median rank within each unit and the same one-sided binomial test against 1/8 (Supplementary Table 15g); these were not specified in the criteria. As a descriptive specificity check, we counted how often endodermis ranked first for other labelled cell types with at least 20 cells, excluding unknown, contaminant, proliferating, stressed and hydathode labels; this comparison did not calibrate the 1 in 8 null. Cell bootstrap intervals from 200 resamples within bundle-sheath and other cells gave uncertainty in AUROC and in the margin over the next program. The random-program control was computed for endodermis only on uncentred weighted-rank scores using 1,000 draws, each sampling one ortholog group with replacement for each program group from the same detection-rate bin while excluding endodermis-program groups. This matching on detection rate replaced the expression matching named in the frozen criteria, and the Bonferroni threshold for eight programs was post hoc. The vascular control compared endodermis with the best of the pericycle, phloem, procambium and xylem programs using a one-sided Wilcoxon signed-rank test for endodermis greater, which gave $P = 0.485$. Sensitivity analyses removed rice, C4 species or datasets with fewer than 100 bundle-sheath cells. The pre-specified C4 set coded maize as non-C4 and removed only the two C4 dicots, giving 6 of 10 with $P = 5.1 \times 10^{-4}$. We also report the corrected set without all C4 species, giving 6 of 8 (Extended Data Fig. 10d and Supplementary Table 14d). Datasets from the C4 species *Z. mays*, *Flaveria bidentis* and *Gynandropsis gynandra* were compared with C3 datasets by whether endodermis ranked first, using a two-sided Fisher's exact test. Labels for GSE184511 and GSE201931 were recovered from marker expression rather than supplied as author annotations; these datasets were included as specified in the criteria, and without them endodermis ranked first in 7 of 10 datasets, with a one-sided binomial $P$ of $4.0 \times 10^{-5}$ (Supplementary Table 15f,g). The root dictionary was trained on 31 datasets from 9 species, 4 of which were also evaluated in leaf: *A. thaliana*, tomato, maize and rice; rice accession CRA004082 contributed separate root and leaf cells, with no shared cell identifiers.

**Follow-up analyses.** Follow-up analyses were exploratory. Their criteria were written before computing the follow-up results, but after the primary and control results were known and after inspecting dataset label names and epidermis-program gene symbols. For the epidermis control, ortholog groups annotated from the literature or Gene Ontology as root-hair or nutrient-uptake genes, 16 in total, were removed from the epidermis program, the uncentred weighted-rank score was recomputed, and the gain was compared with 200 random removals of the same size. For vascular contamination, in each of the 10 datasets with at least 50 separately labelled vascular cells, vascular cells and mesophyll or epidermal reference cells were randomly divided into discovery and held-out subsets after excluding precursor labels. We selected the 50 ortholog groups with the largest discovery difference in mean log1p(CP10k), requiring detection in at least 10% of discovery vascular cells and excluding endodermis-program groups; their equal-weight mean within-cell percentile rank defined the vascular score. Bundle-sheath cells at or above the 25th percentile of the held-out vascular cells were called vascular-like and removed, and AUROCs were recomputed with the exported type scores. For gene function, single ortholog-group AUROCs of bundle sheath versus other cells were computed for endodermis-program groups detected in at least 1% of cells; a group was called enriched when its AUROC exceeded 0.55 in at least half of the testable datasets. To explore associations with gene set and statistic, marker sets and program gene sets were each evaluated by the held-out gene-level AUROC and by a cell-level AUROC of an equal-weight rank score, with the phloem, xylem and procambium programs merged to match the single root vascular marker set (Supplementary Table 15h). The sensitivity analysis re-evaluated the exported type scores under alternative target-label definitions; it did not reproduce the background-cell filtering of the marker analysis (Supplementary Table 15i).

**Chromatin accessibility.** This analysis was exploratory, with criteria written before computation. Cell-type-level accessibility tables of 500-bp accessible chromatin regions (ACRs) were obtained for rice from GEO accession GSE252040 on the MSU7 assembly and for maize from GEO accession GSE155178 on the AGPv4 assembly. Each ACR was assigned to a gene when its centre lay between 2 kb upstream of the transcription start site and the 3’ end of the gene, using the MSU7 annotation for rice and the Ensembl Plants release 44 annotation for maize. Genes were mapped to eggNOG ortholog groups as for the expression data, with rice RAP identifiers converted to MSU identifiers. Within a fixed set of differentiated leaf cell types, or root cell types for the positive control, each ACR was converted to a percentile rank across cell types; clusters with the same label were averaged first. Gene scores were the mean over ACRs, ortholog-group scores the mean over genes, and program scores the dictionary-weighted mean over ortholog groups. Programs were not adjusted using ATAC data. For bundle sheath, each program was compared with 1,000 random draws with seed 0, each sampling one ortholog group with replacement for each scored program group from joint quintile bins of ACR number and mean accessibility, with dictionary weights randomly reassigned; $P$ = (1 + number of random sets at or above the observed score) / 1,001. Species were analysed separately and $P$ values were not combined or corrected across programs. Rice leaf was the primary analysis and rice seedling a sensitivity analysis. Both species are grasses represented in root dictionary training; the tables provide no replicate-level inference, and proximity assignment does not establish regulation.

### Supplementary analyses

Both analyses followed criteria fixed before computation (Supplementary Notes 1 and 2).

**Program-learning ablation.** The released learner was compared with four simpler ways of filling the same programs, each scored with the same scorer and refinement on the 26 benchmark datasets: the 50 or 25 ortholog groups ranked highest by a one-versus-rest Wilcoxon test with weights set to 1, the first step of the released learner alone, and the released programs with all weights set to 1. The primary contrast was the released learner versus the top-50 set at the source level across 18 sources, with a percentile bootstrap 95% CI and Holm correction over the four comparisons. The same contrast was repeated across the seven root species, each left out in turn, with three training seeds.

**Cell types absent from the dictionary.** Each of the eight root types was removed from training in turn, and the eight root test datasets were annotated with the resulting dictionary and with CellTypist trained on the same cells. For each dataset, we computed the AUROC of the negative confidence, defined as the scPCIP margin or the CellTypist maximum probability, for cells of the removed type against all other evaluable cells; a method was considered to flag unseen types if the 95% CI of this AUROC exceeded 0.5 for at least six types and its mean was at least 0.70.

### Pre-specification and deviations

We use “pre-specified” for criteria recorded before the corresponding computation in internal protocols; the study was not registered externally. The departures from these criteria were as follows. Analyses without pre-specified criteria are labelled exploratory in the text.

- Benchmark: the use of macro-F1 as primary for GSE225118; the post hoc removal of SRP286275; and the marker-based root label exclusions; retaining these labels gave a root mean macro-F1 of 0.65, still above OMG, CellTypist and scANVI.
- Re-evaluation: released dictionaries replaced the earlier dictionaries named in the original confirmatory criteria, so the analysis was not blind.
- Spatial: the soybean section filter and dictionary update after the first radial result, and the median-ρ comparison.
- Conservation: the uninterpretable secondary control for H5a, the post hoc matched analysis and the use of a detection-matched rather than expression-matched control in the program-level bundle-sheath test.
- Bundle sheath: the coding of maize as non-C4 in the pre-specified C4 set, the addition of study blocks and lineages, and the post hoc Bonferroni threshold.

### Reproducibility

Analyses used scPCIP 1.0.0, Python 3.11 with numpy 2.4, SciPy 1.17, scikit-learn 1.8 and Scanpy 1.11, and R 4.5.3 (Supplementary Table 16). Seeds are fixed in the code. All refined scoring used single-threaded BLAS, and repeated single-threaded runs reproduced calls line for line. Each figure can be regenerated from the scripts and source data in its package, as described in the Code availability section.

### Supplementary Note 1. Program-learning ablation

This ablation locates where the accuracy comes from: the ortholog groups that enter each program, rather than the procedure used to select them.

The released learner (A0) was compared with four arms that keep every other step unchanged, including the training cells, labels, non-identity filter, scorer, background centring and refinement. The arms were A1, the 50 ortholog groups with the highest one-versus-rest Wilcoxon score in pooled training cells, using log CP10k and positive fold change, and weights set to 1; A2, as A1 with 25 groups; A3, the consensus-contrast ranking that forms the first step of the released learner, truncated to the length of the A0 program, with weights set to 1; A4, the A0 programs with all weights set to 1. Replaying A0 inside the arm framework reproduced the released benchmark exactly, with a maximum absolute deviation of 0.0 over 26 datasets.

**Supplementary Table N1. Macro-F1 difference, A0 minus arm, at the source level across 18 sources.**

| Arm | Mean difference | 95% CI | Sources favouring A0 | Holm P |
|---|---|---|---|---|
| A1, Wilcoxon top 50 | +0.014 | −0.020 to 0.046 | 11 of 18 | 0.57 |
| A2, Wilcoxon top 25 | +0.040 | 0.016 to 0.063 | 12 of 18 | 0.040 |
| A3, contrast ranking only | +0.040 | 0.001 to 0.080 | 14 of 18 | 0.13 |
| A4, released genes, weights 1 | +0.004 | −0.023 to 0.029 | 10 of 18 | 0.57 |

The pre-specified primary contrast of A0 versus A1 gave no evidence that the released learner outperforms a top-50 differential-expression set; at the dataset level, the mean difference was +0.030, and the released learner scored higher in 17 of 26 datasets. The released learner was better than the shorter top-25 set (Holm-adjusted P = 0.040); its advantage over its own first step alone was not significant after Holm correction (P = 0.13), and learned weights added little over equal weights.

**Root, species absent from training.** The A0 − A1 contrast was repeated in root leave-one-species-out runs across 7 species and 3 training seeds, using species-level means. A0 replay deviated by 0.0 from the published runs. The mean difference was +0.035, with a 95% CI of −0.011 to 0.090, a positive difference in 4 of 7 species and Wilcoxon $P$ = 0.47; this did not meet the pre-specified criterion of mean > 0, CI lower bound > 0 and $P$ < 0.05. Per-species differences were: *Arabidopsis thaliana* +0.169, *Medicago truncatula* +0.103,

*Solanum lycopersicum* +0.033, *Lotus japonicus* +0.018, *Oryza sativa* −0.013, *Glycine max* −0.030, *Triticum aestivum* −0.034.

Conclusion: annotation accuracy depends on which ortholog groups enter each program, as shown in the Results, but not measurably on the specific learner used to select them, provided that enough informative groups are chosen.

### Supplementary Note 2. Cell types absent from the dictionary

For each of the eight root types, the type was removed from all training cells and the eight root test datasets were annotated; cells of the removed type can then only be mis-called. Low confidence, measured by the scPCIP margin or CellTypist maximum probability, should rank them below other cells. Datasets with fewer than 20 cells of the removed type were skipped.

**Supplementary Table N2. AUROC of low confidence for cells of the removed type versus all other evaluable cells, given as the mean over datasets with a bootstrap 95% CI.**

| Removed type | Datasets | scPCIP | CellTypist |
|---|---|---|---|
| Cortex | 8 | 0.69 (0.63–0.76) | 0.61 (0.55–0.68) |
| Endodermis | 7 | 0.66 (0.63–0.70) | 0.60 (0.54–0.66) |
| Epidermis | 7 | 0.58 (0.51–0.64) | 0.56 (0.48–0.66) |
| Pericycle | 4 | 0.68 (0.62–0.74) | 0.55 (0.46–0.64) |
| Phloem | 5 | 0.62 (0.52–0.69) | 0.57 (0.46–0.67) |
| Procambium | 3 | 0.71 (0.59–0.85) | 0.70 (0.65–0.79) |
| Root cap | 3 | 0.67 (0.51–0.77) | 0.60 (0.50–0.69) |
| Xylem | 4 | 0.59 (0.53–0.65) | 0.60 (0.53–0.67) |
| Mean of types | | 0.65 | 0.60 |

For scPCIP the confidence interval excluded 0.5 for all eight types, and its mean of 0.65 exceeded the CellTypist mean of 0.60, with scPCIP higher in seven of the eight types; both means nevertheless fell below the pre-specified 0.70, so neither method met the criterion for flagging unseen types. Cells of a removed type were mostly called as a related type of the same lineage; for example, 87% of *Arabidopsis* cortex cells were called as endodermis. The margin therefore indicates unreliable calls but should not be used to discover cell types that are missing from the dictionary.

### Supplementary Note 3. Additional benchmark, spatial and evolutionary details

This note collects the design choices, sensitivity analyses and exceptions that bear on how the benchmark, spatial and evolutionary results should be read. Some strengthen the reported comparisons: the marker-turnover difference is positive in all six root species, and the highlighted leaf–root correspondences remain significant after Holm correction (Fig. 6c). Others mark where a result is narrower in scope than the headline or does not hold.

**Training data.** Flower results reflect within-family transfer, because the flower dictionary was derived from four Brassicaceae species in one study. Some ortholog groups contribute to multiple programs. An exploratory stem dictionary was evaluated by leave-one-species-out on *Arabidopsis* inflorescence stem, soybean hypocotyl and rice culm datasets, each fold trained on the other two, with three random seeds; its mean macro-F1 of 0.16 was below that of random gene sets (0.19) and of the best single marker per cell type (0.22), so no stem dictionary was released. A stem dictionary will be released when more stem data become available.

**Benchmark design.** The SPmarker baseline could not predict pericycle or procambium, which its marker list does not cover. scPlantLLM and scPlantAnnotate were trained on about 10,000 cells: 10,000 root cells and all flower and seed cells. OMG's bundled reference included several query species, so its evaluation was not strictly leave-one-species-out. scPlantAnnotate was included in the seven-method Holm correction.

**OMG and other marker-based comparisons.** In the unadjusted organ-specific comparisons with OMG, only root reached $P < 0.05$, and the flower comparison involved a single study. At the globular and heart stages of soybean seed, OMG achieved macro-F1 values of 0.69 and 0.71, compared with 0.41 and 0.43 for scPCIP. Conversely, scPCIP achieved higher overall accuracies of 0.53 and 0.57, compared with 0.30 and 0.39 for OMG at the resolution selected for its macro-F1; at heart stage, OMG accuracy reached 0.58 at resolution 0.3. scPCIP assigned many seed-coat cells to embryo, reducing its macro-F1. One further seed dataset contained only embryo as an evaluable class. When the SPmarker baseline used soft voting across its marker sets, the difference was 0.37, with Holm-adjusted $P = 0.094$. scPlantAnnotate results in which at least 99% of cells were assigned to a single type were retained in the benchmark; in the remaining five sources, the difference from scPlantAnnotate was 0.16, with $P = 0.31$.

**Supervised classifiers: sensitivity analyses.** With SRP286275 retained, the difference against CellTypist was 0.04, with a 95% CI of −0.002 to 0.09, and against scANVI was 0.06, with a 95% CI of −0.02 to 0.15; the Holm-adjusted $P$ was 0.27 for both. The difference against OMG was 0.16, with a Holm-adjusted $P$ of 0.011. Excluding the six datasets with scPlantDB-curated labels gave a difference of 0.20 against OMG, with Holm-adjusted $P = 0.027$, and differences of 0.06 against CellTypist and 0.07 against scANVI, with Holm-adjusted *P* = 0.10 for both. Overall accuracies across eight root datasets were 0.61 for scANVI and 0.60 for scPCIP.

**Unified root training.** To remove the difference in root training cells, we retrained the three classifiers on the 65,809 root cells used by scPCIP, merging exodermis with cortex and keeping meristem as a class used only during training. Mean root macro-F1 was unchanged at 0.64 for CellTypist and decreased from 0.62 to 0.61 for scANVI and from 0.33 to 0.30 for SingleR. Source-level differences became 0.05 against CellTypist, with Holm-adjusted *P* = 0.11; 0.09 against scANVI, with Holm-adjusted *P* = 0.047; and 0.31 against SingleR. The mean difference against scANVI changed by less than 0.01, but its adjusted *P* value crossed 0.05.

**Eight re-evaluation sources.** These sources were set aside on 14 October 2026. OMG terminated with an error on GSE240098. The Wilcoxon tests against CellTypist and scANVI were one-sided. Merging the three organs of one study gave three of six sources against each, with Holm-adjusted *P* = 0.58 and 0.44, respectively. The positive means were driven by a rice leaf dataset and an *Arabidopsis* flower dataset in which the classifiers performed poorly, whereas they scored higher in five *Arabidopsis* and *Medicago* root and leaf datasets. Results were unchanged when the root classifiers were trained on the unified 65,809 cells: three of eight sources, with Holm-adjusted *P* = 0.74 for both.

**Label retention.** With the excluded root labels retained, OMG performed better in rice, with 0.62 versus 0.43 for scPCIP, and slightly better in wheat, with 0.49 versus 0.47.

**Per-type F1.** In the two maize zero-F1 pairs, CellTypist also scored low, with F1 values of 0.10 and 0.13, whereas OMG reached 0.95 in one of these datasets and 0 in the other.

**Withheld pan-grass study.** The sorghum and maize datasets contained 6,000 and 4,944 cells. The labels of this atlas were assigned to clusters after cross-species integration, so this is not label-free evidence. Accuracy was the pre-specified primary metric, with support defined as exceeding the best single marker in both species; in sorghum, the majority-class baseline reached an accuracy of 0.39 and cortex cells accounted for 39% of the dataset. Including the withheld study in training increased macro-F1 by 0.05–0.07.

**Spatial evaluations.** Spatial data were obtained from GEO and STOmicsDB. Root, leaf and seed datasets used species-excluded dictionaries, whereas soybean leaf used the released leaf dictionary, which contains no soybean data. The rice seed embryo result used spots pooled over 16 sections. The soybean root epidermis program did not discriminate its class, with AUROC 0.53 and q = 0.63. Two additional evaluations performed significantly better than their random references but had AUROCs below 0.5 and therefore did not meet both criteria: rice root epidermis and soybean leaf vasculature. All approaches were evaluated on spots pooled across sections. Soybean root was evaluated separately in the radial analysis and was not included in the macro-F1 comparison. Soybean leaf reached a macro-F1 of 0.59 compared with 0.62 for the random-dictionary 95th percentile. When refinement was omitted from both scPCIP and the random dictionaries, only four of six datasets exceeded both reference thresholds.

**Radial analysis.** The pre-specified soybean analysis of all 24 sections using an earlier dictionary reached Spearman $\rho \geq 0.8$ in 12 sections, which was below the required 20 and did not pass. Six sections contained fewer than five stele bins in the author annotations, so the expected layer order was undefined; this filter and the update to the current dictionary were introduced after that result. With the earlier dictionary, the same 18 soybean and 14 wheat sections reached $\rho \geq 0.8$ in 12 and 3 sections, also below the thresholds. The wheat sections came from two chips, so these statistics are at the section level and do not represent biological replication. Author-annotated root-cap spots themselves reached only 33% on the root-tip measure, suggesting that the 50% threshold was too stringent. Root-tip localization could not be assessed in wheat transverse sections.

**Rice TRAP.** The cortex and endodermis programs scored a proliferating cell population higher than their corresponding target populations, including in the plate-grown replication batch. Each cell population was represented by three replicates.

**Conserved cores.** The matched analysis used 25 combinations in eight species, and the pre-specified secondary control could not be interpreted because its nearest-neighbour matched sets were unbalanced in 46 of 47 combinations. The seven dN/dS species pairs were *A. thaliana–A. lyrata* and *C. rubella–A. thaliana*, both plotted on the *A. thaliana* row, *M. truncatula–Pisum sativum*, *S. lycopersicum–S. pennellii*, *Z. mays–Sorghum bicolor*, *T. aestivum–Hordeum vulgare* and *O. sativa–O. alta*. Each ortholog group was assigned the median dN/dS, denoted $\omega$, of its reciprocal-best-hit gene pairs and ranked within the species pair.

**Marker turnover.** Program-selected markers were classified separately. The between-study comparison used 2,742 records that could also be scored in another study of the same species, regardless of program membership. The root difference was positive in all six species and was 0.18 without *Arabidopsis*; the leaf difference was positive in three of four species. At AUROC < 0.50, the differences were smaller and their CIs included zero, with differences of 0.06 in roots and 0.04 in leaves, although a sensitivity analysis assessing training species separately gave positive differences of 0.16, with a 95% CI of 0.07–0.25, and 0.10, with a 95% CI of 0.02–0.18, so this comparison depends on aggregation. After excluding program-selected markers from the shared set, the neighbourhood-retention difference remained in the same direction in all four leaf species and five of six evaluable root species. Neighbourhood-retention residuals were calculated as neighbourhood-retention AUROC minus the mean AUROC of reference ortholog groups matched by detection decile in both datasets, averaged across training datasets, training species and held-out datasets.

**RNA and chromatin.** The RNA and chromatin measures were weakly correlated across all evaluated pairs, with Spearman ρ = 0.18 in roots and 0.26 in leaves. Gene activity was calculated from peaks over the gene body and 2 kb upstream, summed across member genes. Coloured pairs required RNA detection ≥ 5 counts per million (CPM) in both species and $\log_2$ specificity ≥ 1 in one species and ≤ 0.25 in the other.

**Leaf–root correspondence.** Sensitivity analyses tested whether the correspondences depended on particular genes or cells (Supplementary Table 15a–e,j,l). Removing 16 root-hair and nutrient-uptake ortholog groups from the epidermis program did not restore the leaf epidermis–root epidermis correspondence: epidermis ranked first in 2 of 10 datasets, compared with 1 of 10 before removal, and the gain exceeded the 95th percentile of random removals in one dataset. After bundle-sheath cells resembling vascular cells were removed, endodermis remained first in 7 of the 8 datasets in which it had ranked first. Only 4 of 39 endodermis-program ortholog groups were individually enriched in bundle sheath, two of them class III peroxidases including RCI3, and these groups were over-represented for detoxification terms. In cell-level bootstrap resampling, endodermis ranked first in at least 80% of resamples in each of these 8 datasets and in none of the resamples in the other 4. Endodermis ranked first for bundle sheath in 8 of 12 datasets but for only 4 of 84 other leaf cell labels. Excluding two blocks with marker-recovered labels retained the bundle-sheath correspondence in 6 of 7 blocks. In the reverse direction, endodermis matched in 2 of 11 blocks and pericycle in 6 of 11. No significant dN/dS difference was detected specifically for the bundle-sheath–inner-layer class. The equal-ranking benchmark is approximate, and dataset independence is not established. Endodermis AUROC was below 0.5 in both maize datasets. The endodermis program exceeded random gene sets in five datasets after Bonferroni correction, although this control did not adjust for selecting the highest-ranking of eight programs. The pre-specified C4 exclusion inadvertently retained maize, and endodermis then ranked first in 6/10 datasets. The observed exceptions did not support a systematic C3–C4 difference, with Fisher's exact $P = 0.55$. Differences in gene sets, scoring and evaluation units also affected these correspondences. The pairs shown in bold had a hit rate ≥ 2/3, unadjusted Poisson-binomial $P < 0.05$ and median ΔAUROC > 0, and remained significant after Holm correction, with $P < 0.01$ (Fig. 6c).

### Supplementary Figures

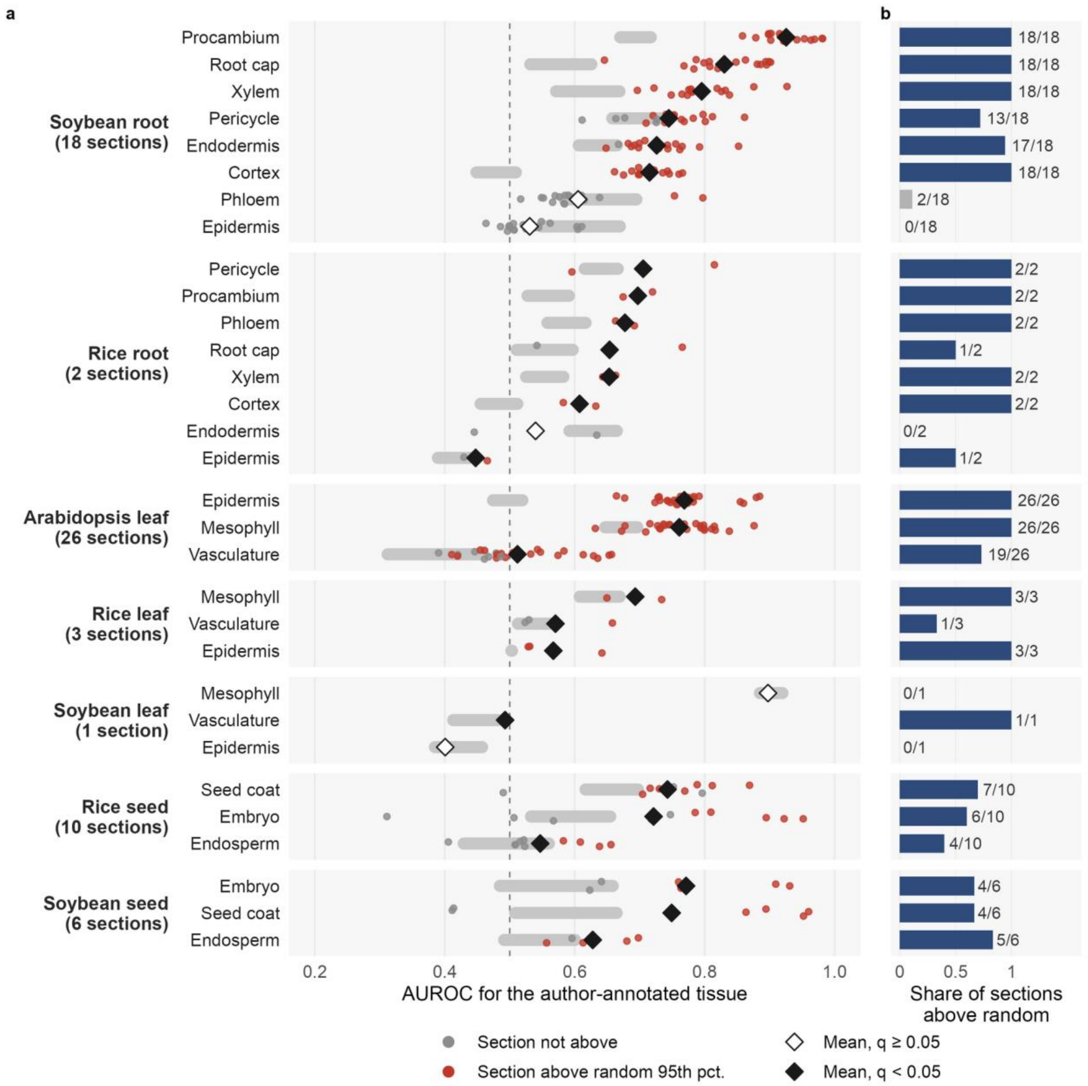


**Supplementary Fig. 1 | Program AUROC in every labelled spatial dataset. a**, AUROC of each program for its author-annotated tissue; points, sections; grey bar, mean to 95th percentile of expression-matched random programs; q, Benjamini–Hochberg adjusted. **b**, Share of sections above the random 95th percentile; bar colour, q < 0.05.

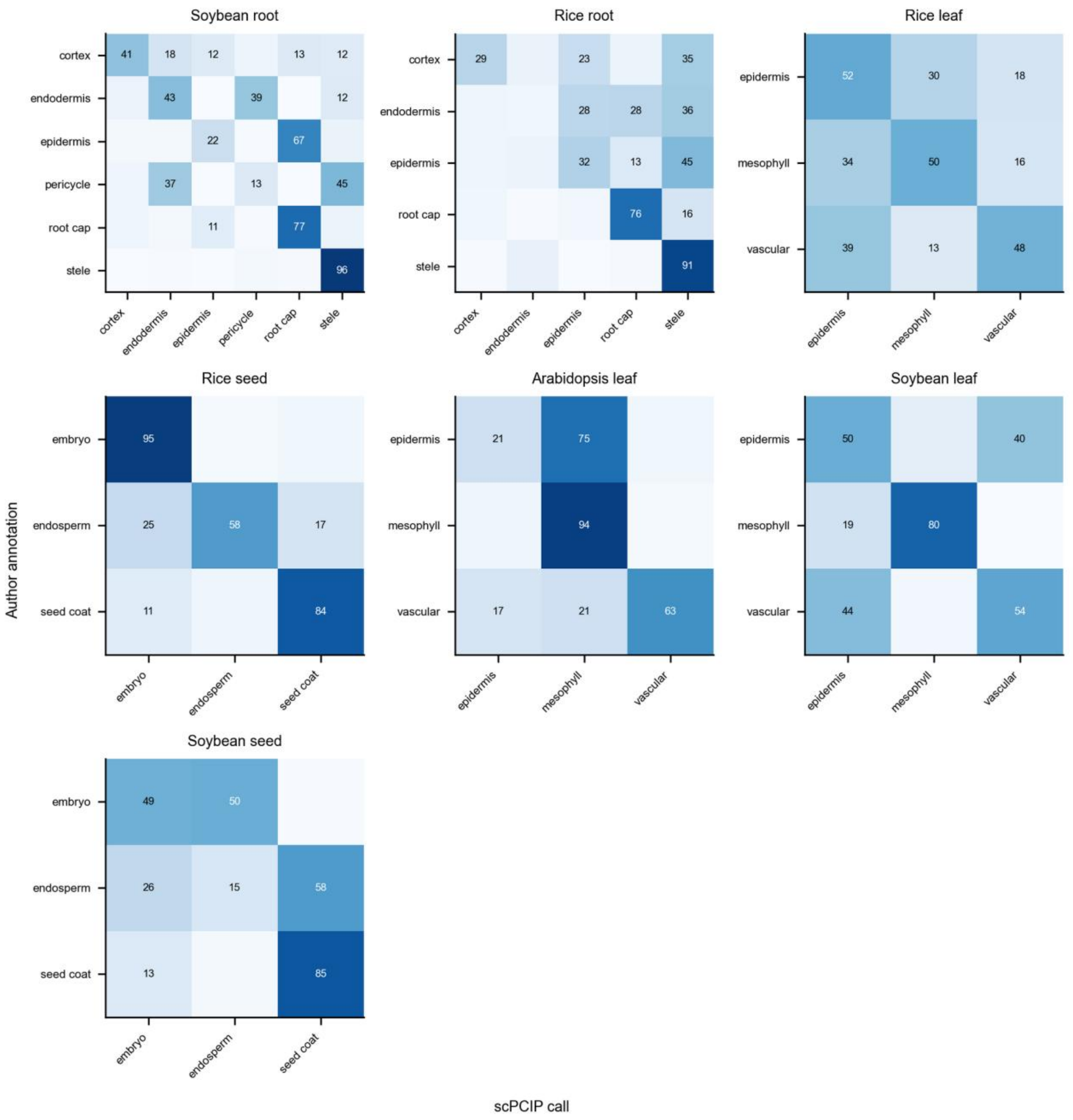


**Supplementary Fig. 2 | Confusion matrices of the spatial datasets.** Numbers, percentage of spots of each author type.

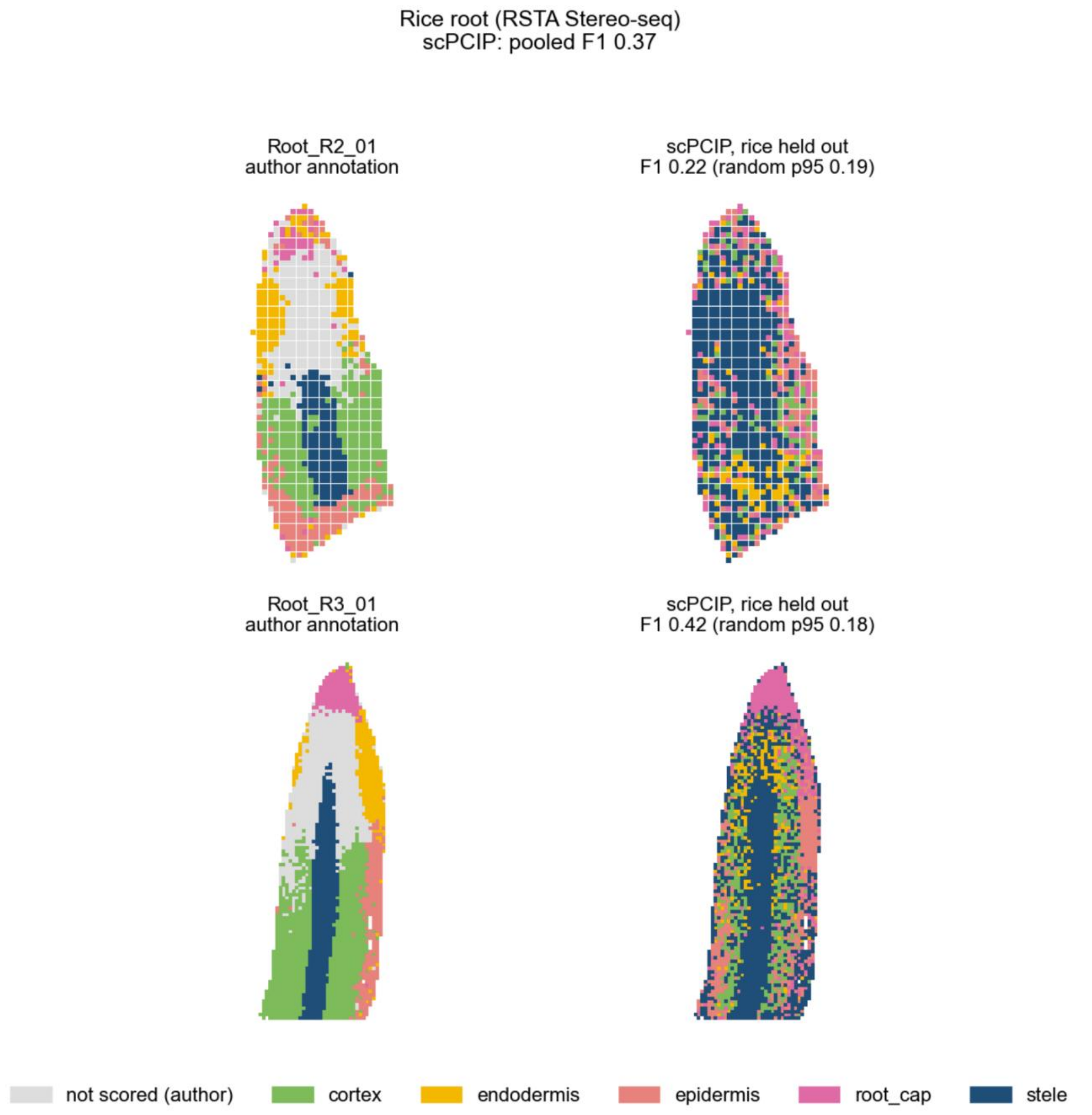


**Supplementary Fig. 3 | Author annotation and scPCIP calls on spatial sections of rice root.** In each pair, the author annotation is on the left and the scPCIP calls from a dictionary trained without the species are on the right. Random p95, 95th percentile of 20 expression-matched random dictionaries for the same section.

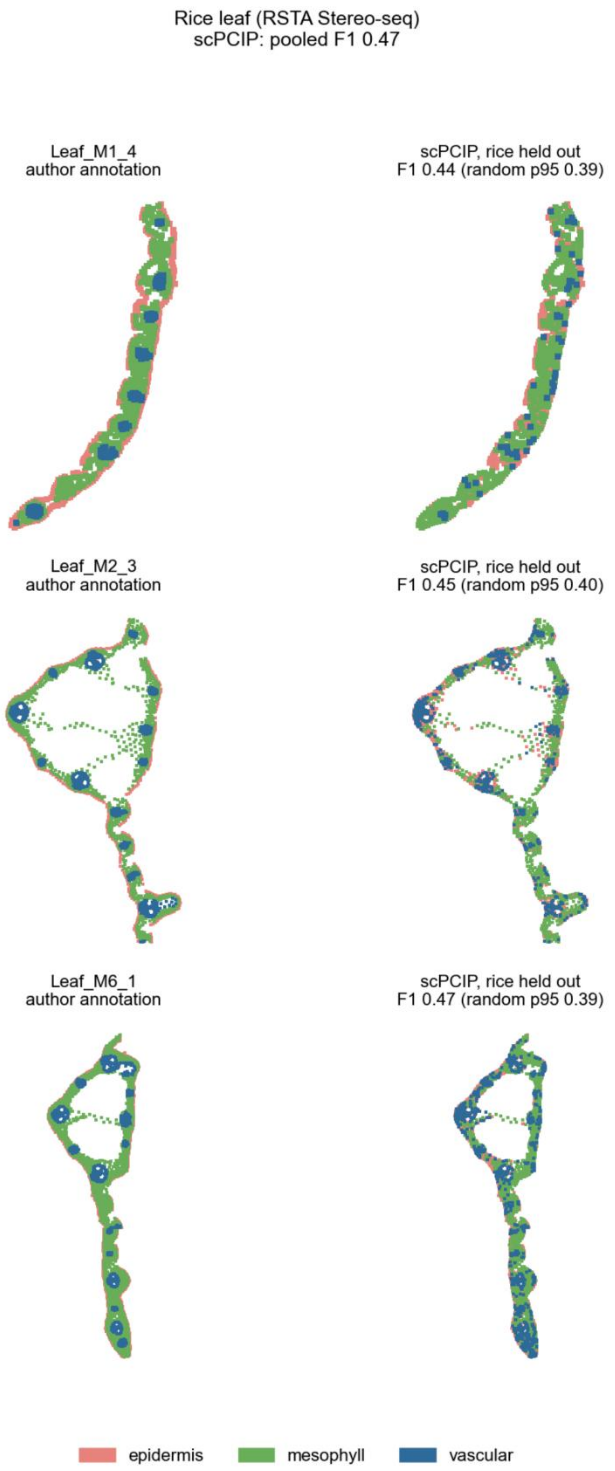


**Supplementary Fig. 4 | Author annotation and scPCIP calls on spatial sections of rice leaf.** In each pair, the author annotation is on the left and the scPCIP calls from a dictionary trained without the species are on the right. Random p95, 95th percentile of 20 expression-matched random dictionaries for the same section.

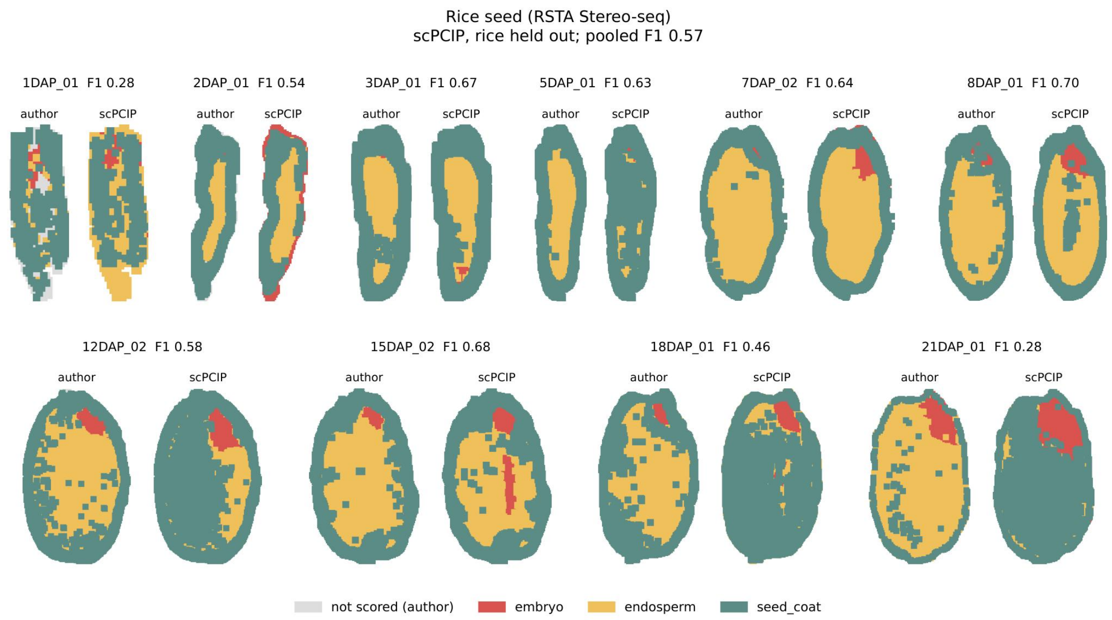


**Supplementary Fig. 5 | Author annotation and scPCIP calls on spatial sections of rice seed.** In each pair, the author annotation is on the left and the scPCIP calls from a dictionary trained without the species are on the right. Random p95, 95th percentile of 20 expression-matched random dictionaries for the same section.

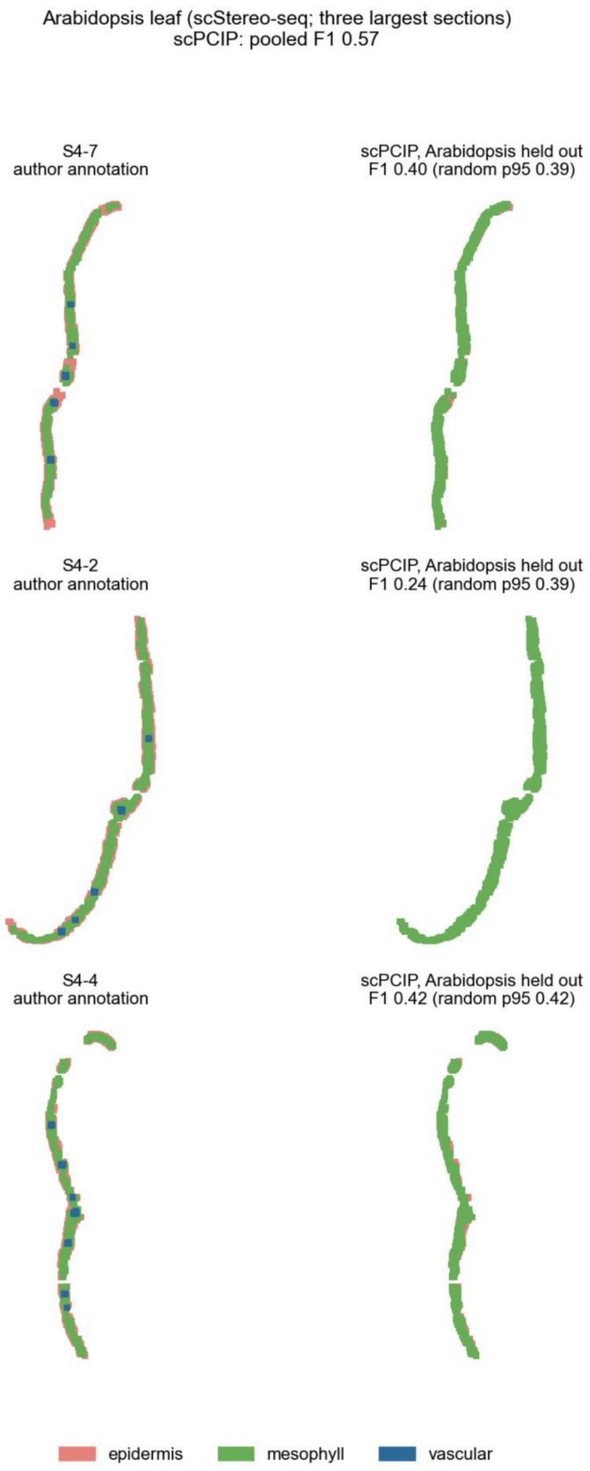


**Supplementary Fig. 6 | Author annotation and scPCIP calls on spatial sections of *Arabidopsis thaliana* leaf.** In each pair, the author annotation is on the left and the scPCIP calls from a dictionary trained without the species are on the right. Random p95, 95th percentile of 20 expression-matched random dictionaries for the same section.

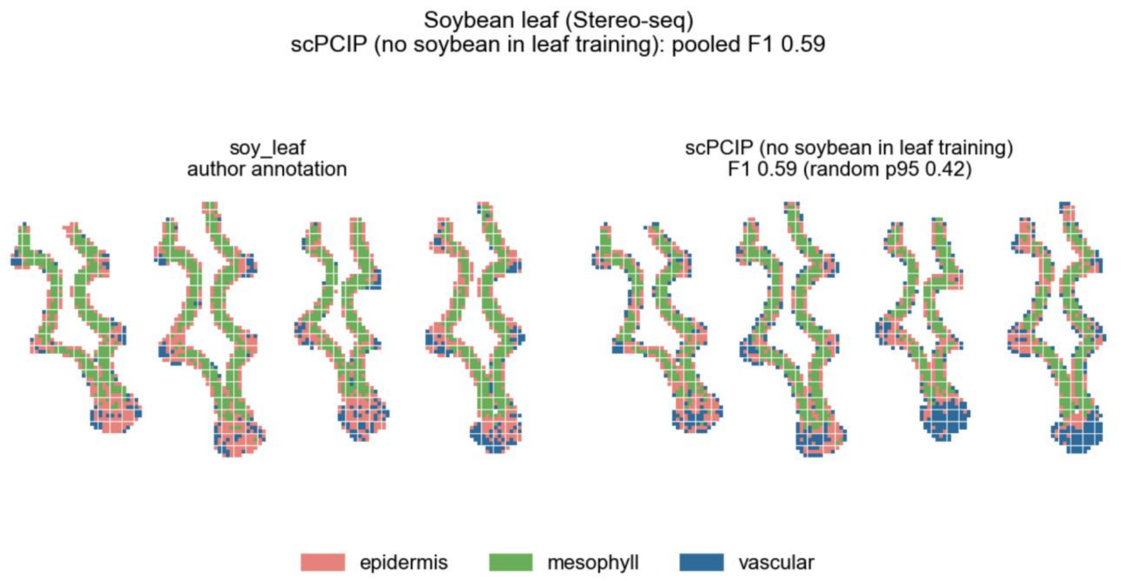


**Supplementary Fig. 7 | Author annotation and scPCIP calls on spatial sections of soybean leaf.** In each pair, the author annotation is on the left and the scPCIP calls from a dictionary trained without the species are on the right. Random p95, 95th percentile of 20 expression-matched random dictionaries for the same section.

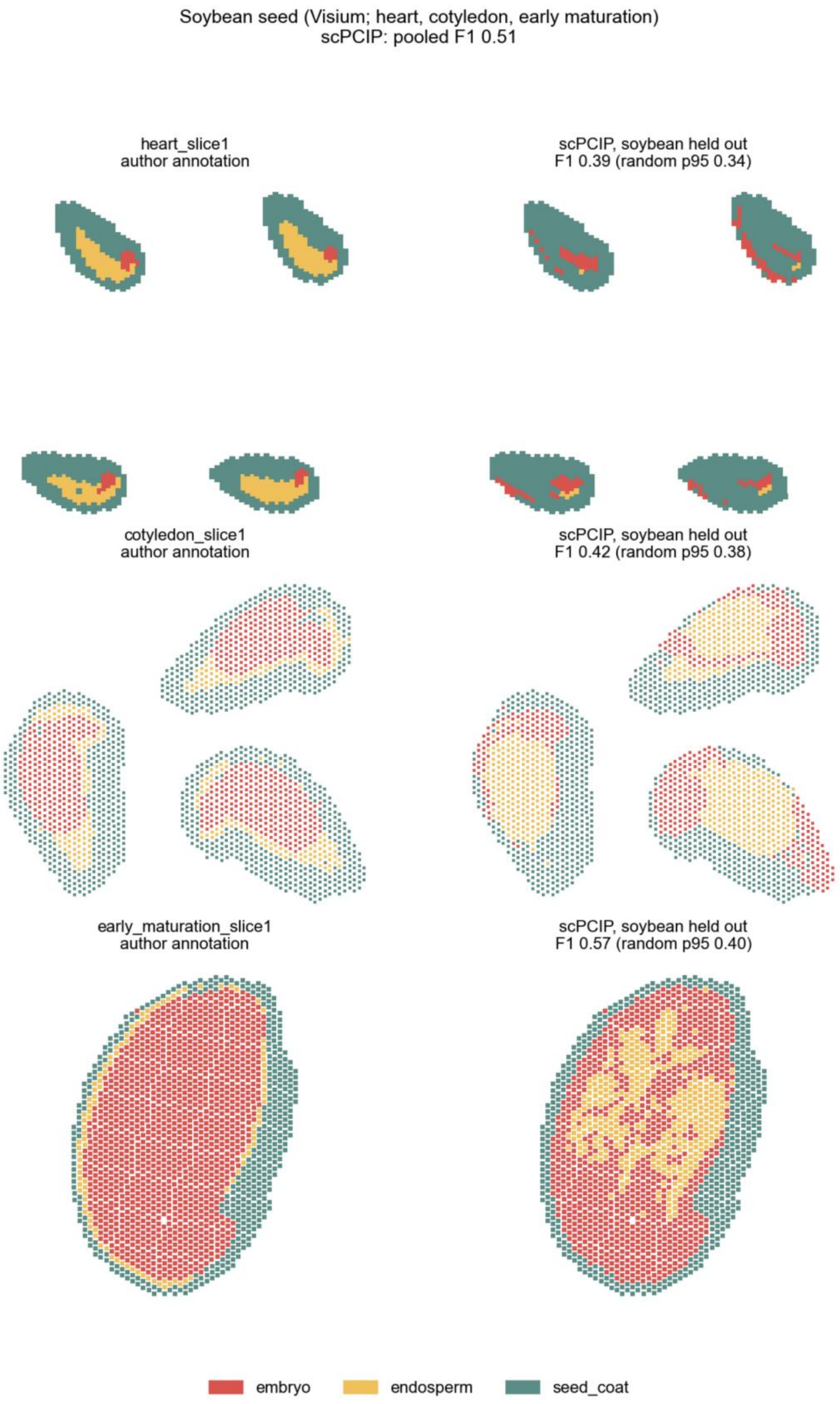


**Supplementary Fig. 8 | Author annotation and scPCIP calls on spatial sections of soybean seed.** In each pair, the author annotation is on the left and the scPCIP calls from a dictionary trained without the species are on the right. Random p95, 95th percentile of 20 expression-matched random dictionaries for the same section.

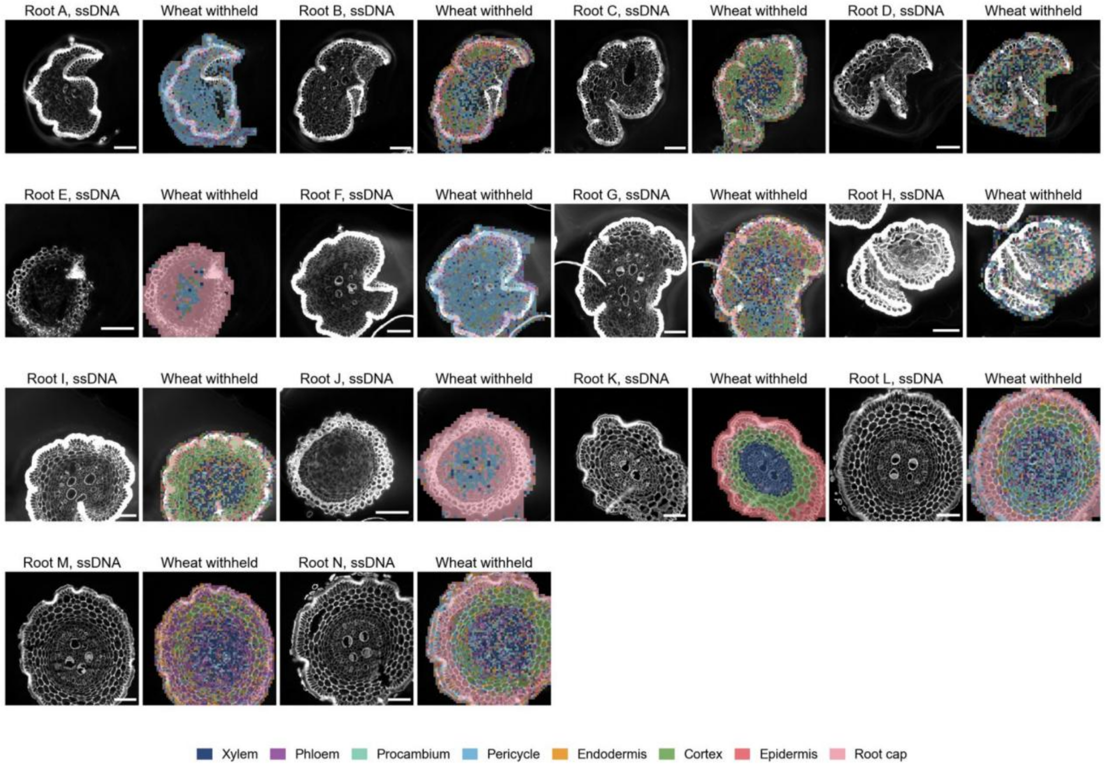


**Supplementary Fig. 9 | All wheat root sections.** ssDNA image and scPCIP calls from the wheat-withheld dictionary for each root. Scale bars, 100 μm.

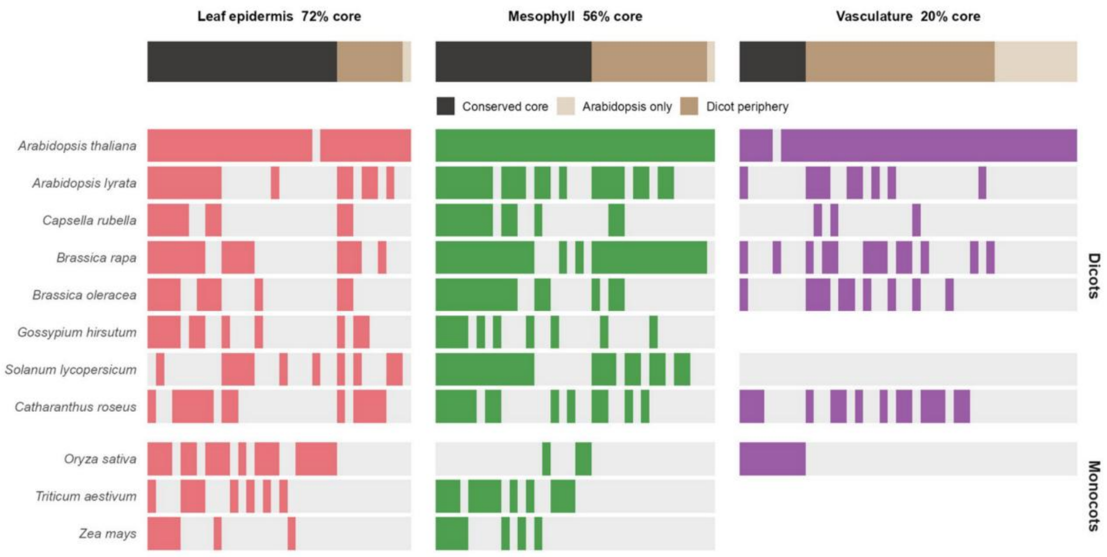


**Supplementary Fig. 10 | Phylogenetic support of leaf program ortholog groups.** Leaf programs, drawn as for the root programs (Fig. 4d).

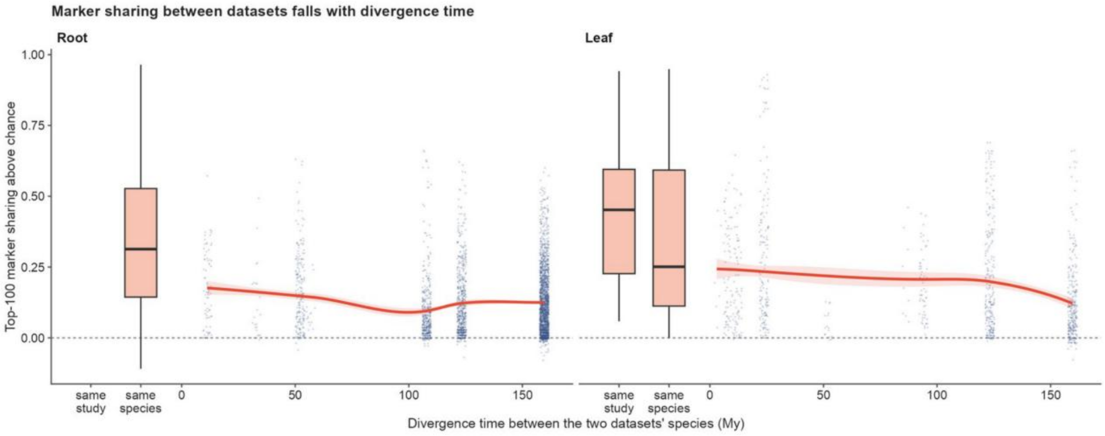


**Supplementary Fig. 11 | Marker sharing and divergence time.** Marker sharing between dataset pairs against species divergence time. Boxes, median and interquartile range for dataset pairs from the same study or from other studies of the same species; whiskers, 1.5 × interquartile range; points, pairs from different species; line, loess fit with 95% CI.

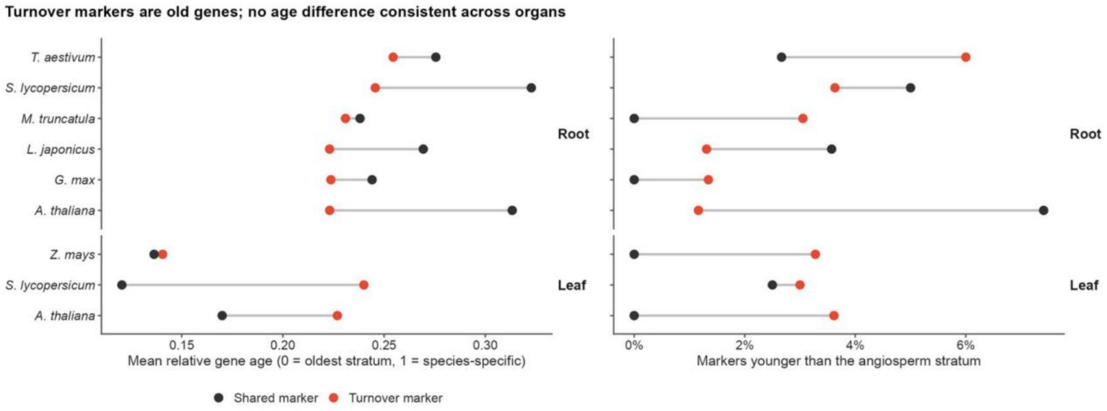


**Supplementary Fig. 12 | Gene age of turnover and shared markers.** Age stratum of turnover and shared markers in root and leaf.

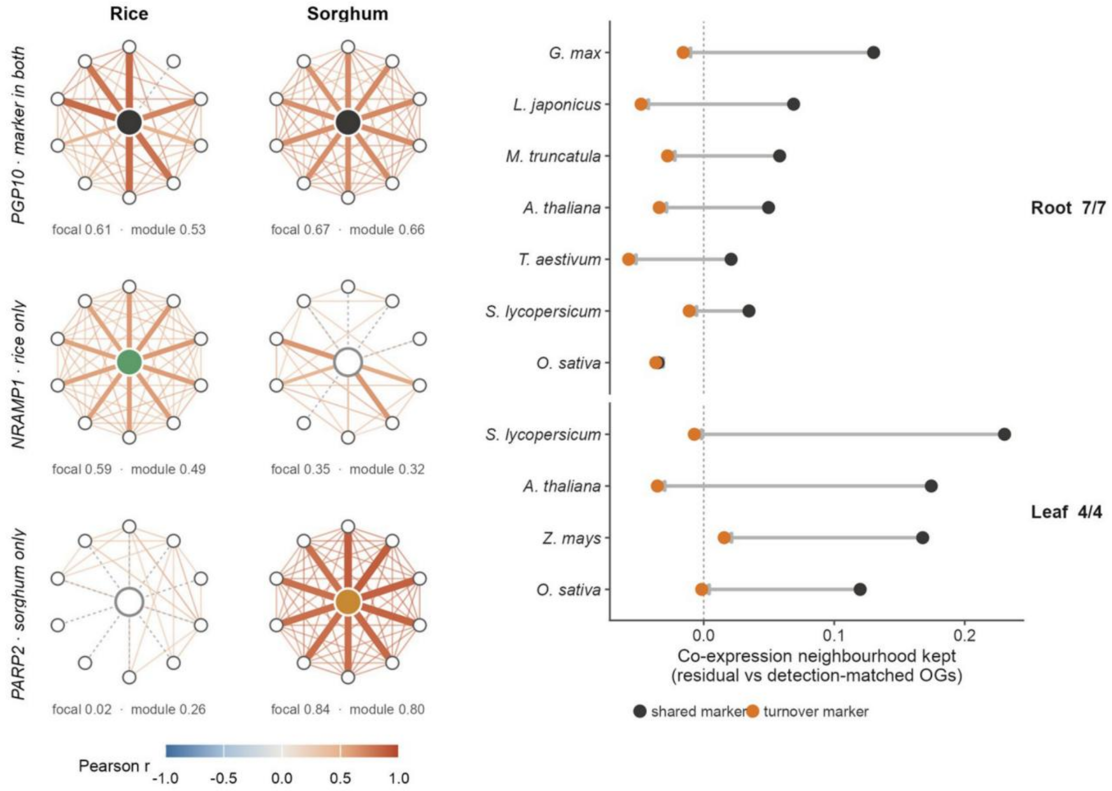


**Supplementary Fig. 13 | Co-expression neighbourhoods of exemplar ortholog groups.** Ego networks in rice and sorghum leaf and per-species neighbourhood retention. Focal, mean Pearson *r* of the edges to the focal ortholog group; module, mean *r* of the other edges.

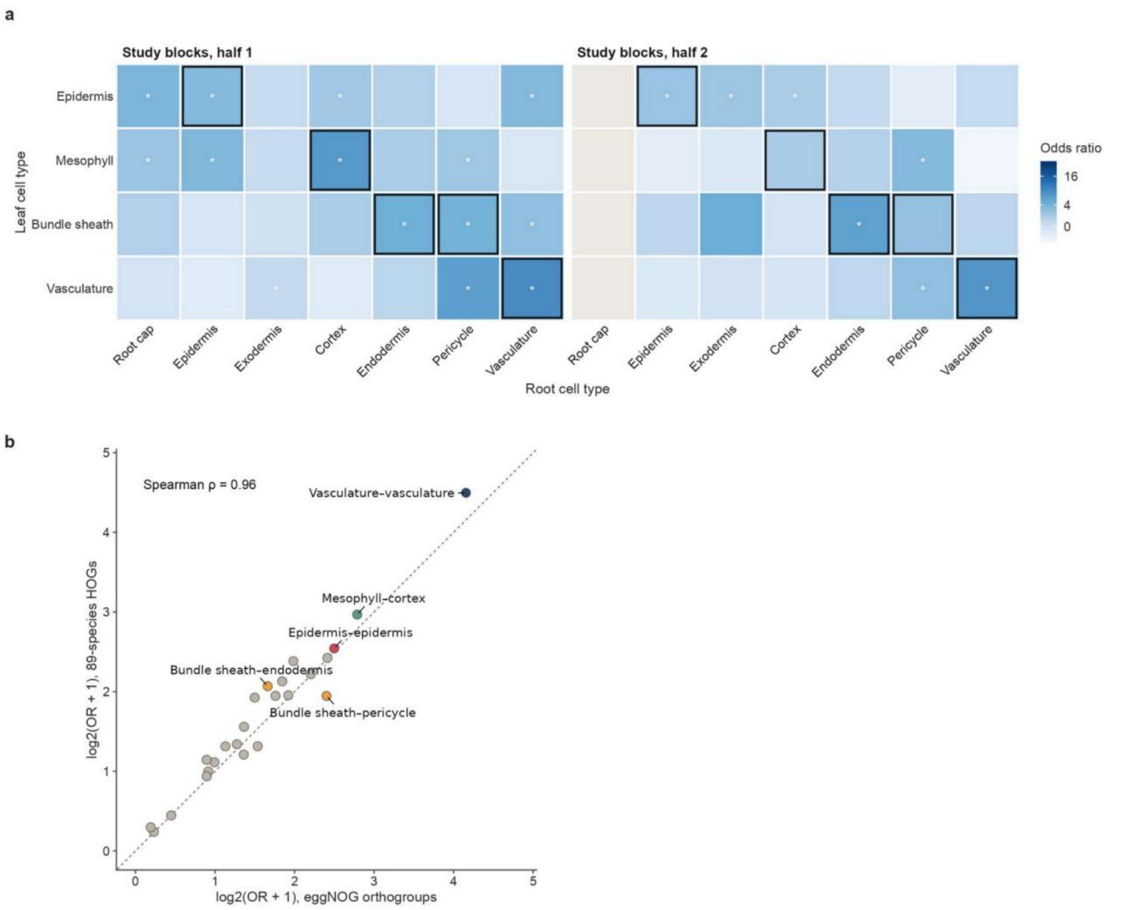


**Supplementary Fig. 14 | Robustness of the leaf–root correspondence. a**, Odds ratios in two halves of the studies; outlined, pre-specified pairs; asterisks, *P* < 0.001. **b**, Odds ratios at the eggNOG and hierarchical-orthogroup levels.

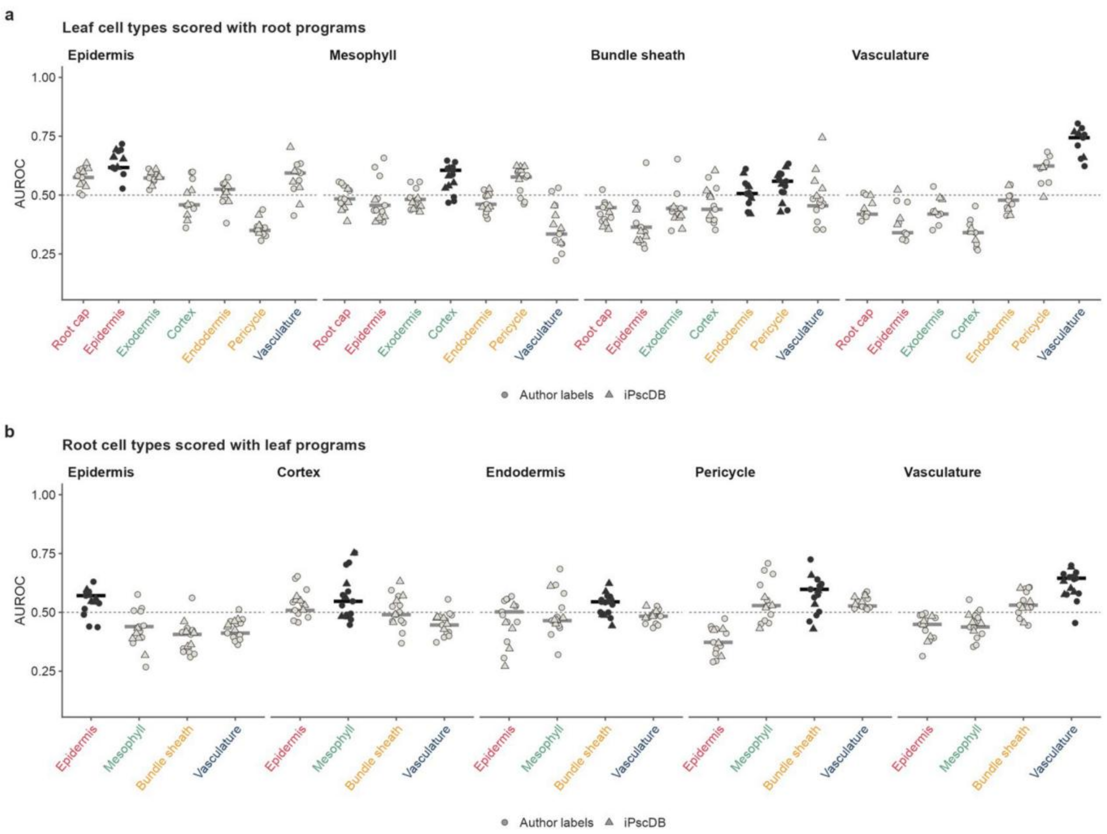


**Supplementary Fig. 15 | Held-out AUROC of every candidate partner.** Dark, preset partner; line, median.

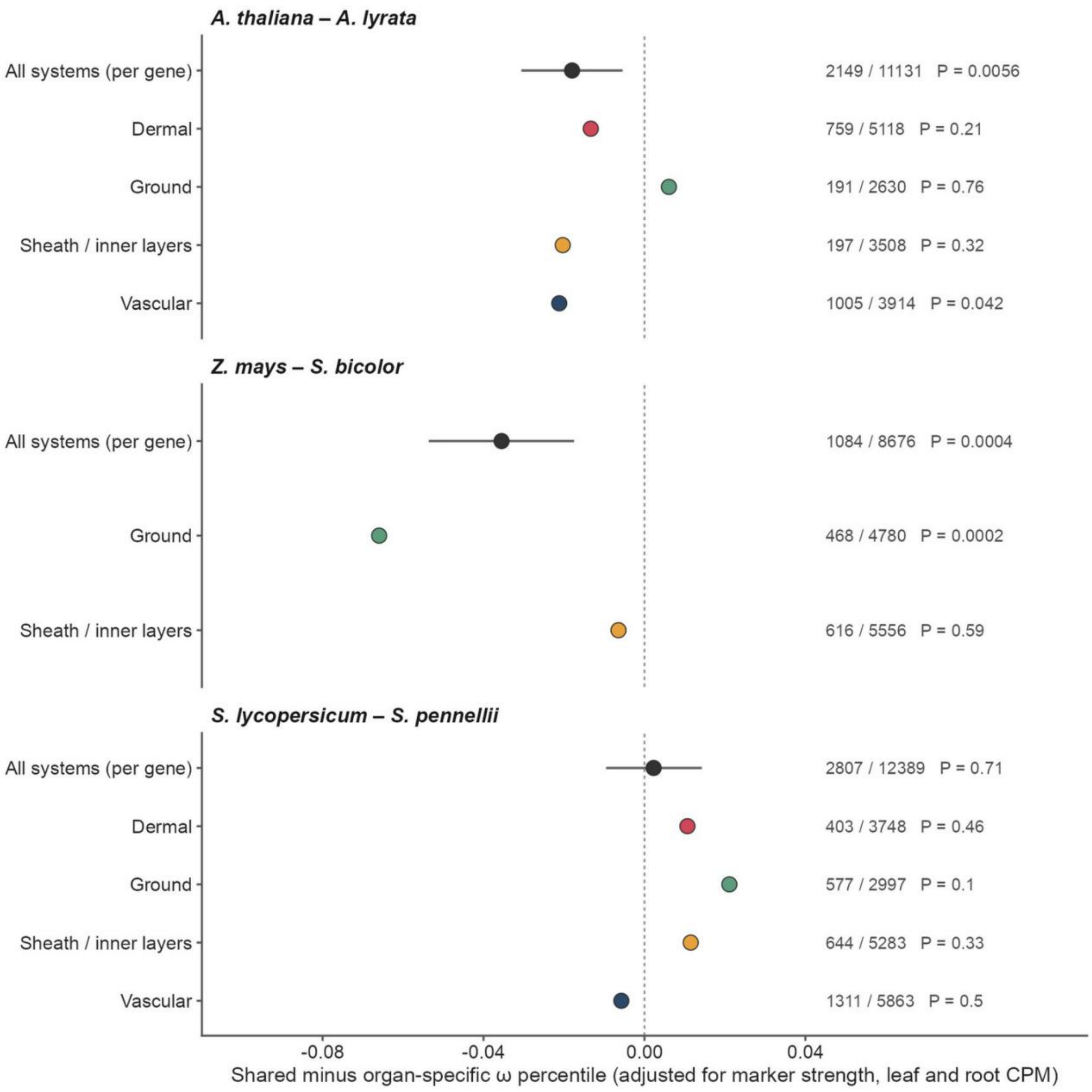


**Supplementary Fig. 16 | Sequence constraint in all species pairs.** Shared minus organ-specific ω percentile per species pair and tissue system, with bootstrap 95% intervals. Numbers, shared markers / all markers and stratified permutation *P*.

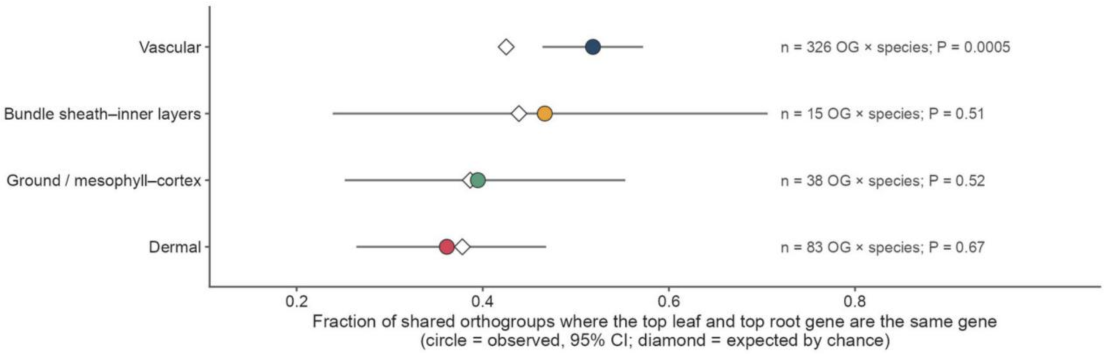


**Supplementary Fig. 17 | Same-gene reuse across organs.** Proportion of shared ortholog groups in which the top gene is the same in leaf and root, against random expectation; *P*, one-sided, from simulated random choice of the top gene. The bundle sheath–inner layer row covers 7 of the 8 ortholog groups shared by bundle-sheath and inner-layer markers (Fig. 6e); the eighth (CBL) had fewer than two detected member genes in each of the four species tested (Arabidopsis, rice, maize and tomato). n, ortholog group × species combinations.

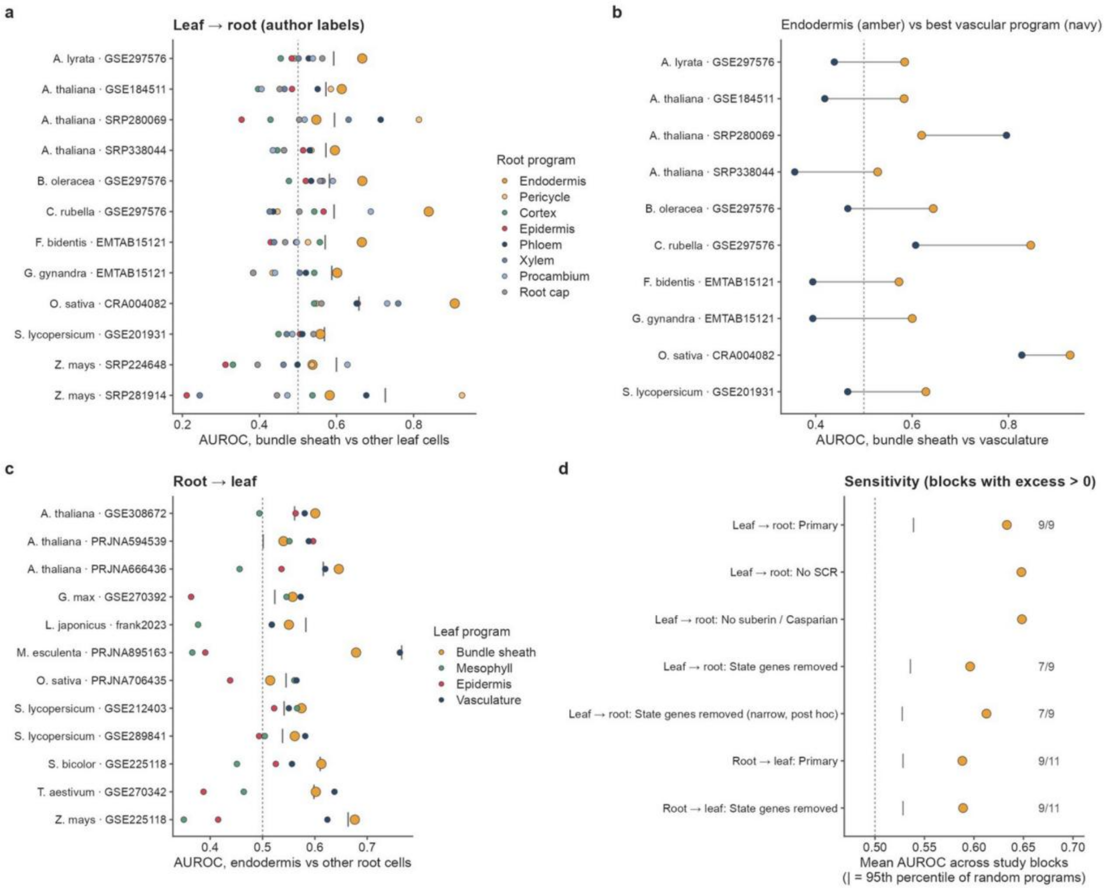


**Supplementary Fig. 18 | Bundle sheath and root inner layers. a**, AUROC of the endodermis and control root programs for bundle sheath. **b**, Endodermis against the best vascular program. **c**, Root to leaf. **d**, Sensitivity to removing gene subsets. Vertical ticks in **a** and **c**, 95th percentile of random programs.
